# Effects of DNA Origami-Based Nanoagent Design on Apoptosis Induction in a Large 3D Spheroid Model

**DOI:** 10.1101/2025.02.12.637807

**Authors:** Johann M. Weck, Riya Nair, Merve-Z. Kesici, Xiaoyue Shang, Cornelia Monzel, Amelie Heuer-Jungemann

## Abstract

The extrinsic activation of programmed cell death by FasR/CD95 is a promising minimally invasive strategy for cancer treatment. This can be leveraged using high-precision nanoscale therapeutics: Utilizing DNA origami for precise Fas ligand (FasL) presentation resulted in over 100 times more potent apoptosis induction in single cells. However, treating large, solid tumors poses challenges for DNA origami-based therapeutics, including drug distribution and altered cellular behavior. Here, we addressed these challenges using a 3D spheroid model. First, we assessed DNA origamis’ ability to penetrate tumor tissue, finding that penetration is influenced by the DNA origami size rather its structural flexibility. Second, we evaluated the apoptosis induction efficacy by DNA origami-FasL nanoagents within the spheroid model. The most potent nanoagents were able to completely eradicate all cells in the spheroid. Results indicated that apoptosis induction depended strongly on FasL attachment strategy rather than DNA origami design. Notably, only a rigid neutravidin linker for FasL attachment, rather than a flexible dsDNA linker, halted spheroid growth and fully eradicated all cancer cells. This study offers critical insights into designing DNA-based therapeutics for complex cellular environments and significantly advances DNA origami nanotherapeutic development, highlighting the impact of nanoagent design on cell fate decisions.

## 1. Introduction

Programmed cell death (apoptosis) plays several important roles in multicellular organisms^[1]^. In embryonic development, apoptosis is crucial for the development of organs and limbs, regressing, for example, the interdigital web in animals^[2]^. In defense against pathogens, cells undergo apoptosis upon pathogen infection^[3]^. Similarly, the induction of apoptosis is an important defense mechanism against potentially malignant tumor cells^[4]^. Apoptosis can either be triggered intracellularly, or induced extracellularly, through an external trigger like the binding of the Fas ligand (FasL/CD95L) to the Fas receptor (FasR/CD95) on the cell membrane.^[5]^ Explicitly this interaction was suggested to require specific oligomerization of the receptors around the ligands^[6]^. Controlling, and selectively triggering these interactions, by mimicking the specific configurations, and thus introducing cell death with minimal invasiveness, has the potential to be an effective cancer therapy^[7]^.

As ligand-receptor interactions take place on the nanoscale, control over them presupposes nanometer precision. One effective way to achieve this is by employing DNA nanotechnology, and especially the DNA origami method^[8]^. It offers control over the size, shape, and functionality of structures on the nanoscale, with almost full addressability. Owing to these unique properties, DNA origami have been used to construct nanoscale rulers^[9]^, molecular motors^[10]^, enact forces on the nanoscale^[11]^, probe plasmonic assemblies^[12]^ as well as biological systems^[13]^. Most recently, the potential of DNA origami as biomedical agents is being explored^[13d, 14]^, sparking the question of how this new category of therapeutics affect complex biological systems.

In a prior study, we already successfully utilized DNA origami to decipher the crucial dependence of nanoscale positioning of FasL for apoptosis induction in a two-dimensional (2D) cell culture model^[13a]^. Only if ligands were placed in hexameric arrangements with ∼ 10 nm inter-ligand distance (ILD), mimicking the proposed FasR cluster pattern^[5]^, were our so-called DNA origami-FasL “nanoagents” over 100-fold more effective at inducing apoptosis compared to the soluble ligand. Larger or smaller ILDs strongly impeded signaling. Smaller oligomers, like dimers, at the correct ILD were also able to induce apoptosis, albeit not as efficiently as the hexameric arrangement^[6, 13a]^. Similar results were found for a TRAIL-mimicking peptide^[13b]^. The ability to drastically enhance the efficacy of the ligand through programmed pre-patterning promises great potential for nanoagents in clinical applications.

However, all of the aforementioned studies were conducted in less physiologically-relevant conditions with cells in 2D, adhered to a surface, with the origami-protein or origami-peptide nanoagents added onto the substrate or into the solution, directly presenting the ligands to the cells. Many cancers *in vivo*, on the other hand, form large, solid tumors in three dimensions (3D). This adds several challenges: Firstly, the transportation to and distribution within the tumor plays a major role in the therapeutics’ efficacy. Secondly, tumors form a different microenvironment, with gradients of pH, nutrients, and metabolic waste, and form necrotic cores at their center, all changing cell behavior in 3D, as opposed to 2D^[15]^. Thus, nanoagents effective in 2D might not yield the same effectivity in 3D, but could even have an inverse effect^[16]^. Many prior studies^[13a, 13b, 14b]^, investigating the feasibility of DNA origami-based cancer therapeutics, fell short in studying essential aspects of nanoagent design affecting their effectivity. Other promising studies^[17]^, investigating different proteins and the influence of their spatial arrangement with DNA origami, also largely disregarded the effects of protein attachment and the underlying DNA origami structure. Therefore, before more elaborate *in vivo* studies can be conducted, it is essential, also from an ethical point of view, that the principal effects of nanoagents on cells in 3D and their apoptosis efficacy are investigated.

In this study, we examined the influencing factors of the overall nanoagent design, from the underlying DNA origami (size/flexibility) to the ligand conjugation strategy, on their efficacy in penetrating into and inducing apoptosis in a solid tumor model. For this, we cultivated large 3D spheroids with diameters of ∼0.5 mm. Using confocal microscopy and fluorescence-activated cell sorting (FACS), we found that smaller DNA origami penetrated into the spheroid much more efficiently, however, all DNA origami nanoagents, independent of the underlying DNA origami design, were efficient at completely inhibiting tumor growth, if ligands were attached *via* a rigid neutravidin-biotin linker. On the other hand, none of the nanoagents were able to halt tumor growth if ligands were conjugated via a more flexible dsDNA linker. Finally, we show that the most effective nanoagents were able to completely eradicate the tumor, leaving no viable cells. With this study, we provide important insights into the intricate design requirements of DNA origami-based nanotherapeutics in complex environments and provide a nanoagent specifically acting only on cancer cells overexpressing CD95, therefore minimizing adverse cytotoxicity.

## 2. Results & Discussion

### 2.1 DNA origami design and characterization

We initially sought to identify the most suitable DNA origami platform, capable of penetrating deep into the spheroid. For this, we examined two parameters of DNA origami design. We hypothesized, that the ability to penetrate spheroids would be affected by the size and potentially also the flexibility of the DNA origami, with smaller and more flexible structures being able to more easily pass through inter-cellular gaps, as previously suggested^[18]^. We thus constructed three different DNA origami structures, differing in size and/or flexibility. The first DNA origami is the Rothemund Rectangle Origami (rro), widely used as a molecular pegboard^[9, 13a, 19]^ (see **Figure 1a** and S1), which we already successfully employed as a nanoagent^[13a, 13b]^ before. Deviating from the rro in size, but not in structure, we designed a miniature rro (mini) from a miniature scaffold^[20]^ (see **Figure 1a** and S2). Thirdly, we designed a wireframe origami^[21]^ (wf) with similar dimensions as the rro, but with highly increased flexibility (see **Figure 1a** and S3).

**Figure 1:**
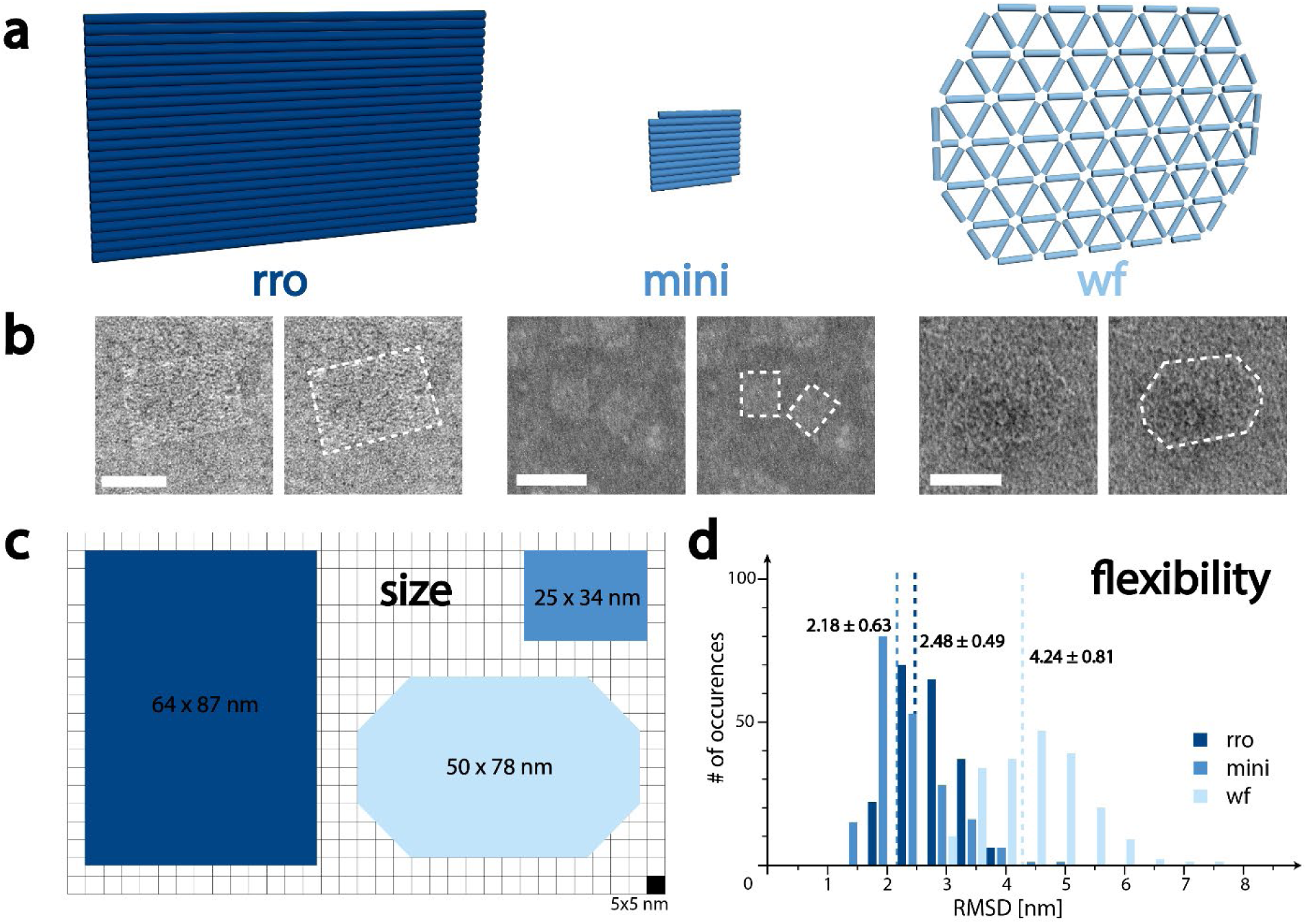
**DNA origami structures** of various sizes and flexibilities. (**a**) Rendering of DNA origami structures. Color scheme (rro: dark blue, mini: medium blue, wf: light blue) also holds for all future figures. (**b**) Characterization of DNA origami via TEM. Micrographs are duplicated and the right micrograph shows the DNA origami nanostructure outlined for better visibility. (**c**) Dimensions of the DNA origami extracted from micrographs show approximately the same dimensions for rro and wf and much smaller dimensions for the mini DNA origami. (**d**) Simulated RMSD of DNA origami structures show similar values for rro and mini and much more fluctuations, almost double in the wf structure. Scale bars in (**b**) are 50 nm.

Analysis of folded structures by transmission electron microscopy (TEM) showed well-formed structures in high agreement with the designed origami (**Figure 1b**, S4a, S5a, and S6a). The length and width of each structure were extracted from TEM micrographs and for the rro was determined as 87.4 ± 4.2 nm by 64.2 ± 3.6 nm (see **Figure 1c** and S4b), which is comparable to the length (77.9 ± 5.5 nm) and width (50.5 ± 4.2 nm) of the wf structure (Figure S6b). The mini origami structure displayed a length of 33.9 ± 3.1 nm and a width of 24.9 ± 2.1 nm, at least 4x smaller than the other two structures, as can be seen in **Figure 1b-c** and S5.

Next, to quantify the flexibility, we performed oxDNA^[22]^ simulations and extracted the root mean squared fluctuations (RMSF). The number of occurrences of fluctuations from the mean were then plotted as a histogram, seen in **Figure 1d**. In contrast to the size dimension analysis, rro and mini displayed much lower fluctuations than the wf origami. The average fluctuation of the rro was 2.48 ± 0.49 nm, similar to the average of the mini (2.18 ± 0.63 nm), whereas the wf had an average fluctuation of 4.24 ± 0.81 nm, almost twice as large, suggesting its much higher degree of flexibility.

Having three different DNA origami structures varying either in size or in flexibility, we next sought to investigate the influence of these parameters on the origami’s tumor penetration ability. N.B. we did not consider a rod- or spherical-shaped origami, due to the requirement of hosting a planar arrangement of 6 FasL with 10 nm ILD in the final nanoagent design.

### 2.2 DNA origami penetration into spheroids

To elucidate the impact of DNA origami size and flexibility on their penetration ability, we measured the respective diffusion speeds through the 3D spheroids. For this, spheroids were grown for three days after which the DNA origami structures were added for set time intervals. For subsequent visualization, spheroids were fixed, permeabilized and optically cleared to enable imaging of the entire structure by confocal microscopy.

Spheroids were produced by seeding HeLa Apo12 mGFP cells^[13a]^ in ultra-low adhesion well plates. In three days after seeding, the cells formed a large (approx. 0.5 mm) three-dimensional spheroid, visible even by the naked eye (see **Figure 2a** and S7a). After three days of growth, 0.5 pmol of the respective DNA origami was added to the culture medium and then incubated with the spheroids for different time intervals. The penetration was stopped by several washing steps and fixation of the spheroids, which were subsequently permeabilized and optically cleared. To visualize the DNA origami position in the spheroids, we applied a modified version of the recently reported origami FISH method^[23]^, where staple strands were extended from the origami for subsequent hairpin attachment (see Figure S7b). This would also allow us to indirectly infer about the DNA origami stability: if staples detach from dissociated DNA origami, we would expect a very faint homogeneous fluorescence signal throughout the whole spheroid (see Figure S7c), whereas in case of intact structures, the fluorescence signal would be more intense and spot-like, due to higher concentrations of the respective staples in the same location. Spheroids were then imaged via confocal microscopy and the average penetration depth was extracted and analyzed using a clockscan protocol^[24]^ (see Figure S8).

**Figure 2:**
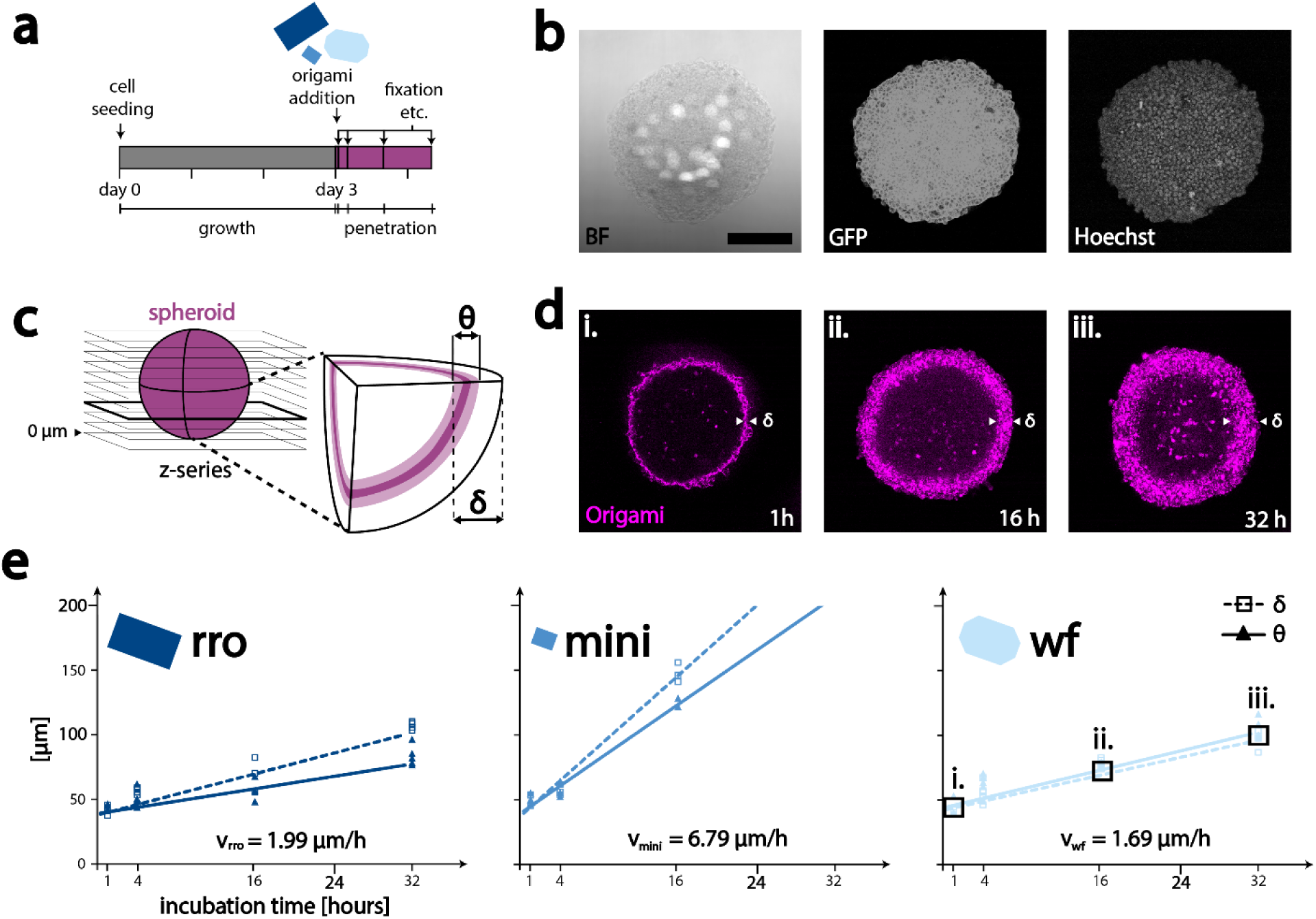
Penetration of DNA origami in spheroids. (**a**) Experimental timeline: cells were seeded and grown for 3 days. Origami (blue) was added on the third day. Spheroids were subsequently fixed after set time periods to stop penetration. (**b**) Cross section of a cleared spheroid: brightfield shows low contrast, GFP tagged FasR outlines the cell walls, and Hoechst 33342 stain indicates cell nuclei. (**c**) Schematic of image acquisition and determination of penetration depth δ and ring thickness θ. (**d**) Exemplary cross section of spheroids with origami incubated for different time periods: longer incubation shows deeper penetration into tumor tissue. Images are from wf origami incubations and time points i., ii., and iii. are also indicated in the graph below. (**e**) Penetration depths analysis of different DNA origami structures: penetration depth δ data is shown with square boxes, with a dotted trendline. Datapoints on ring thickness θ are shown as solid triangles, with a solid trendline. Respective origami structures are indicated in shades of blue. The mini origami shows the fastest penetration through the spheroid tissue, rro, and wf origami penetrate much slower. mini and rro show δ and θ deviating from each other significantly after 16 h. The scale bar in (**b**) is 0.2 mm and holds for all microscopy images. n= 3 per time-point for all structures.

Confocal microscopy in combination with optical clearing allowed us to image completely through the half-millimeter-thick spheroids. Clearing is crucial, as not clearing the spheroid would allow only for imaging approximately 100 µm into the tissue, leaving everything beyond this inaccessible to the microscope. The use of confocal microscopy allows for slicing through the spheroid, instead of imaging a projected intensity sum. As seen in **Figure 2b** we were able to image without loss of quality through a ∼ 0.5 mm thick spheroid: The GFP signal from cell wall-anchored FasR-GFP, as well as cell nuclei stained with Hoechst 33342 were visible, even in the center, with no signal quality loss, enabling a confident analysis throughout the whole spheroid.

The penetration depth of the DNA origami into the spheroid is identifiable as a fluorescent ring. The fluorescence in the ring is distributed granularly, distinctly different from the even fluorescence signal in the control (Figure S7c). With increasing incubation time, the ring becomes thicker and reaches deeper into the spheroids, indicating deeper penetration of the DNA origami (for a more in-depth analysis see Figures S9-S11). Spheroids, incubated with the respective DNA origami structure for a given time interval, and treated as discussed above, were imaged in slices as a z-series, as shown in **Figure 2c**. The penetration depth δ and the ring thickness θ were extracted from the slice at the thickest part of the spheroid. In **Figure 2d** three exemplary images of spheroids are shown, each with wf DNA origami structures incubated for different time intervals.

The penetration behavior for each DNA origami structure was analyzed and kinetic profiles were plotted. We observed the fastest penetration kinetics for the mini origami (see **Figure 2e**). rro and wf origami penetrated the spheroids much slower. Fitting linear trendlines (dotted line) and extracting the slope, the mini DNA origami showed a penetration speed v_mini_ of 6.79 µm/h, more than three times faster than the speed of the rro (v_rro_ = 1.99 µm/h) or the wf origami (v_wf_ = 1.69 µm/h). Penetration data of spheroids after 16 h incubation with the mini origami still showed the presence of a fluorescent ring, however after 32 h of incubation, a distribution of fluorescence could be observed throughout the whole spheroid (see Figure S10d). The fluorescence signal here showed the same granularity of signal across the whole spheroid, unlike the very homogeneous, low fluorescence background in the control (Figure S7c). We hypothesize that this suggests full penetration of the mini DNA origami through the spheroid. This is further supported by the fact that the calculated penetration depth after 32 h (∼220 µm), obtained from extrapolation of the data in **Figure 2e**, roughly corresponds to the spheroid radius. We attribute this behavior to the much smaller size of the mini DNA origami, allowing it to better penetrate through gaps in the cell-cell junctions, as well as increasing the general diffusion speed. Interestingly, we found that the fluorescence patterns of rro and mini looked qualitatively different from wf patterns when incubated for 16 h or more: the ring thickness θ was not equivalent to the penetration depth δ anymore, as seen in Figures S9 and S10, and plotted as solid triangles and fitted with a solid line in **Figure 2e**. Rather, θ became smaller than δ, as the outer parts of the spheroids did not show a strong fluorescence signal anymore. Since all three DNA origami behaved similarly when exposed to a serum-containing medium (Figure S12, showing high stability of all structures), we hypothesized that the difference in θ and δ may be due to origami in the tighter square lattice design interacting stronger with cells compared to the wireframe origami, as already proposed in a previous study^[18]^, resulting in cellular uptake and subsequent intracellular degradation. Degradation of the DNA origami would either render the anchor staples for FISH inoperable or lead to the unhindered diffusion of the anchor staples and thus a uniform distribution of comparably lower fluorescence signal in the spheroid (cf. Figure S7c).

We found that each DNA origami displayed unique interaction characteristics with the spheroids, allowing us to draw several conclusions: (i) origami size is the decisive factor in penetration speed. The mini origami, with a ∼4 times smaller size penetrated the spheroid with a speed of 6.79 µm/h, more than 3 times faster than rro and wf origami (1.99 and 1.69 µm/h). (ii) The influence of DNA origami flexibility appears to be negligible, as the wf and the rro origami showed very similar penetration speeds and depths. This is in contrast to previously published results, finding wf origami to penetrate better through spheroids^[18]^. Nevertheless, in the aforementioned study wf rods were compared to origami rods, whereas here we utilized flat, rectangular structures. (iii) On the other hand, the “circulation time” of DNA origami in the spheroid was found to be mainly dependent on the internal structure, rather than the origami size, as the rro and mini origami signal vanished at the edge of the spheroid after longer periods of incubation, which was not the case for the wf origami signal. Having identified the mini origami as the ideal structure for spheroid penetration, we next sought to investigate, whether the differences in penetration pattern also translate into differences in nanoagent efficacy.

### 2.3 Nanoagent design and characterization

To construct the functional nanoagents, FasL was conjugated to all three origami structures^[13a]^. For this, we developed a recombinant FasL, trimerized via an isoleucine zipper (IZ), to mimic naturally occurring, active trimeric FasL^[25]^. Through the incorporation of an additional Cys on the IZ, the protein was further functionalized with either a DNA strand or a biotin moiety, allowing for two attachment strategies to the origami structures (see **Figure 3a**). In previous work, we showed that the attachment with more flexible DNA linkers, compared to more rigid streptavidin linkers, resulted in slower apoptosis kinetics^[13a]^. To study whether this was also the case in a 3D environment, we also tested both attachment strategies here. On each of the three origami we positioned six FasL in the most optimal, hexagonal conformation with an ILD of ∼10 nm^[13a]^ (see Figures S1a,c, S2a,c, and S3a,c) either via a dsDNA linker or an intermediate neutravidin.

**Figure 3:**
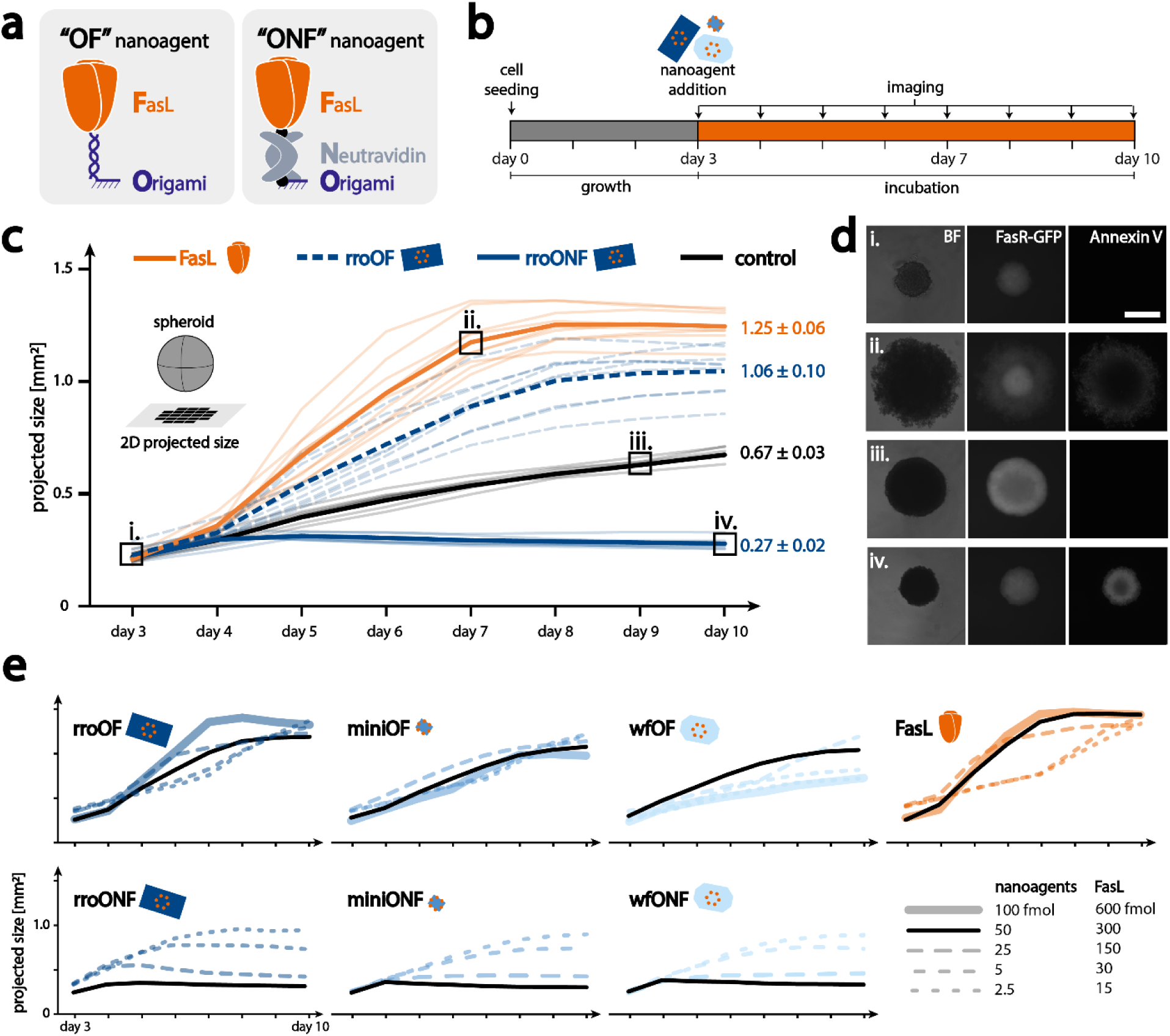
Tumor spheroid killing assay: **(a)** schematic representation of attachment strategies of FasL to the DNA origami: OF indicates direct attachment of FasL to origami via DNA hybridization, ONF indicates attachment via a neutravidin. (**b**) Experimental timeline: cells were seeded and grown for 3 days. Nanoagents (blue) or FasL (orange) were added on the third day. The spheroids were imaged daily for a whole week. (**c**) Size graphs of spheroids as projection on 2D: orange lines are spheroids incubated with 300 fmol FasL, dotted blue lines: 50 fmol OF nanoagent, solid blue lines: 50 fmol ONF nanoagent, and black lines are controls. Thin lines indicate single experiments and bold lines indicate averages (n=9). Boxes with Roman numerals indicate time points for images in (**d**). (**d**) Representative snapshots of spheroids in brightfield, GFP, and AnxV channels. Roman numerals refer to time points in (**c**), i. and iii. are from a control, ii. from a spheroid with FasL, and iv. from a spheroid with ONF nanoagent at day 10. (**e**) Average size graphs of spheroids incubated with different amounts of nanoagents, indicated by dotted lines with varying spacing. Averages were from n=9 for 50 fmol (or 300 fmol) experiments, and n=3 for the other experiments. Individual data points can be found in Figures S20-S26. The legend shows black dashed lines which are generalized for color code in different nanoagents. The scale bar in (**d**) is 0.5 mm.

Nanoagents with FasL connected via dsDNA hybridization are abbreviated OF (Origami-FasL) nanoagents, while nanoagents with a neutravidin connector are abbreviated ONF (Origami-Neutravidin-FasL) nanoagents. Successful formation of all nanoagents was verified by TEM imaging, seen in Figures S13-S15 for OF nanoagents and in Figures S16-S18 for ONF nanoagents.

### 2.4 Tumor spheroid killing assays

To investigate their efficacy, we incubated HeLa Apo12 mGFP spheroids with the different nanoagents and observed the spheroid behavior over one week by live cell imaging. The development of spheroid size, as well as the qualitative morphological changes, were examined, accompanied by Annexin V (AnxV) staining to visualize apoptotic cells. As seen in **Figure 3b**, spheroids were seeded and grown for 3 days, as described above for the penetration assays. On the third day, the respective nanoagents were added and spheroids were imaged on a fluorescence microscope once per day for one week. From those images, the spheroid size was extracted as a planar projection. In **Figure 3c** the development of spheroid size, incubated with different nanoagents, is shown. The control spheroids (see **Figure 3c** as a black line), grew steadily, undisturbed, and almost linearly from ∼ 0.2 mm^2^ to ∼ 0.65 mm^2^ (projected area). Images in the brightfield and GFP channels confirmed this observation (see **Figures 3c** and **d** i. and iii.). In both cases, the spheroid is almost perfectly round, and respectively larger after the longer incubation time. The GFP intensity was much higher on day 9, indicating more GFP proteins due to a larger number of cells. Further, there was no signal from the AnxV stain, indicating no ongoing apoptosis, as expected. As can be seen in Figure S19, this behavior is very similar for different control conditions, indicating that spheroid development is independent of the addition of buffer, non-functionalized DNA origami, and AnxV marker.

The behavior of spheroids changed drastically when soluble FasL (300 fmol) was added into the medium (orange lines in **Figure 3c** (and S20). Spheroid size increased rapidly, starting one day after FasL addition, and reached a plateau at day 7 or day 8, at ∼ 1.2 mm^2^, approximately twice the projected area as the control spheroids. The same behavior was also observed when rroOF (50 fmol) nanoagent, carrying 6 FasL, was administered (**Figure 3c**, dotted dark blue line, and S21): spheroid size increased rapidly, starting 1 day after nanoagent addition, and plateaued at day 8 / 9 at a size of ∼ 1.0 mm^2^. This increase in projected tumor size, however, is not due to increased proliferation, but rather a change in spheroid morphology, as revealed by brightfield (BF) and fluorescence images. In **Figure 3d ii.** a representative image of spheroids incubated with soluble FasL at day 7 is shown. In the BF channel, the large spheroid displayed very rough, unshaped borders. We attribute this variance to apoptotic cells partially detaching from the spheroid core. This was further supported by an observable change in fluorescence intensities of FasR-GFP and AnxV. In the GFP channel, a higher-intensity core and a lower-intensity ring could be observed. The inverse could be seen in the AnxV channel, indicating the presence of apoptotic cells in the outer ring, induced by FasL or rroOF nanoagents. Cells towards the center of the spheroid however, were not affected and showed higher GFP and no AnxV signal. N.B.: Images were recorded on a regular fluorescence microscope, and are thus not total projections of z-stacks, resulting in a general decrease in fluorescent signal in the middle of very large and thick spheroids, as light cannot penetrate through the thicker parts of the spheroid. As seen in **Figures 3c** and S20-S21, spheroids incubated with FasL or rroOF, showed a larger variance in development than the controls. Surprisingly, the underlying DNA origami structure of the nanoagent had little to no influence on the spheroid fate. As shown in the lower graphs in **Figure 3e** and also Figures S22 & S23, a similar overall behavior was also observed for the miniOF and the wfOF (50 fmol). While spheroid behavior did not change significantly with varying DNA origami architectures, we did, however, see a drastic change when varying the FasL attachment strategy.

When ONF nanoagents (50 fmol), where FasL was attached to the DNA origami via neutravidin, were administered to the spheroid, the spheroid size peaked at day 4/5 and subsequently decreased, as depicted in **Figure 3c** as a solid blue line (also shown in Figure S24). The final size of these spheroids was found to be only 0.28 mm^2^, which is less than half that of the control spheroids. As can be seen in **Figure 3c**, the spheroid incubated with rroONF at day 10 was found to have approximately the same size as at the beginning of the incubation. It furthermore displayed a round and even morphology. GFP fluorescence was low, but a very strong Annexin V signal could be detected, indicating a large amount of apoptotic cells. We suppose this is due to the strong capability of rroONF nanoagent to induce apoptosis, in agreement with our previous 2D studies^[13a]^. This behavior was also found to be independent of the underlying DNA origami structure: depicted in **Figures 3c** (and S25-S26) the miniONF and wfONF nanoagents induced the same halt in spheroid growth, with final spheroid sizes reaching only 0.26 mm^2^ and 0.29 mm^2^, respectively. The size graphs for spheroids with each ONF nanoagent were very uniform, showing very consistent behavior, as depicted in Figures S24-S26. To investigate if this behavior was concentration-dependent, and whether OF nanoagents could induce the same behavior as ONF nanoagents if concentrations were increased, we next carried out titration experiments.

### 2.5 Attachment strategy proves to be more important than origami structure

To investigate the influence of nanoagent concentration, we used the same experimental setup as before but varied the amount of all nanoagents and soluble FasL added to the spheroids. Initially, to test whether OF nanoagents or soluble FasL could be as effective as ONF nanoagents, we increased their amount to 100 fmol (OF nanoagents), or 600 fmol (soluble FasL) and subsequently analyzed the respective spheroid growth curves. Interestingly, increasing the amount of soluble FasL did not show any significant difference in final spheroid size, although a temporal difference in the growth curve could be observed (see **Figure 3e** and Figure S20). Similarly, increasing the amount of OF nanoagents did not affect the final spheroid size (see also Figure S21-23). It merely shifted the onset of the spheroid growth plateauing to an earlier time point.

Next, we investigated the effect of a decreased amount of nanoagent (ONF, OF) to 25 fmol, 5 fmol, or 2.5 fmol, and similarly of soluble FasL to 150 fmol, 30 fmol, or 15 fmol on the growth of spheroids. Interestingly, ONF nanoagents were still highly potent, completely halting tumor growth, even if only 25 fmol were added. This behavior was very similar for all ONF nanoagents (rro, mini, wf), see **Figure 3e** (lower four graphs, and Figures S24-S26). The final spheroid sizes were approximately 0.1 mm^2^ larger than that of spheroids treated with 50 fmol ONF nanoagent. Further reduction of ONF nanoagent amount to 5 or 2.5 fmol again led to a halt in tumor growth but resulted in a slightly larger final spheroid size. This behavior was found to be qualitatively distinctly different from the plateauing behavior of spheroids incubated with large amounts of OF nanoagents or FasL, suggesting a different or more potent mechanism of action (see Figure S27). Finally, reducing the amount of soluble FasL or OF nanoagents only caused a delay in the formation of the ring of apoptotic cells (see **Figure 3e** and Figures S20-23).

We summarize that contrary to our expectation, ONF nanoagents were found to not only be more effective than OF nanoagents or soluble FasL but also induced a different behavior in the spheroids, which cannot be evoked by merely increasing the amount of the less effective nanoagents. To gain more insights into the observed behavior on a cellular level, we next investigated cellular viability in the spheroids more thoroughly.

### 2.6 Quantifying the apoptosis induction efficiency

To quantify the apoptosis induction efficiencies of the different nanoagents, we devised two additional experiments: Firstly, we directly analyzed the cells of dissociated spheroids by FACS. Secondly, we dissociated the spheroids and re-seeded the cells in 2D. As spheroids only exhibited different behavior with regards to the FasL attachment, but not to the origami structure, we felt confident to only compare nanoagents based on the rro DNA origami.

After the addition of nanoagents and 10 days of incubation, spheroids were dissociated and directly analyzed via FACS (**Figure 4a**). As cells undergo apoptosis, their size and granularity change. Both can be detected with FACS, as forward scatter (FSC) and sideward scatter (SSC), respectively^[26]^. Initially, we compared the FACS distribution of cells cultivated in 2D or 3D. As can be seen from S28a, cells cultivated in 3D, cells displayed a smaller overall size (lower FSC values) compared to those grown in 2D, but still formed a visible population, which we assigned as “viable”. Secondly, upon the addition of nanoagent and induction of apoptosis, the amount of cells in the viable population decreased and the amount of smaller cell fragments, partially with higher granularity (SSC) increased (see Figure S28b). We subsequently defined gates around both populations and determined the apoptosis induction efficacy as a fraction of cells in viable or dead gates from the total number of cells in both gates, as indicated in **Figure 4b**. Here, larger proportions of dead cells indicate higher efficacy of apoptosis induction of the respective nanoagent. In agreement with previous observations, spheroids incubated with soluble FasL, rroOF, and rroONF as well as control spheroids showed drastically different ratios of dead and viable cells, consistent over two fully separate sets of measurements (Figure S29). As depicted in **Figure 4c**, cells grown in 2D showed mostly cells in the viable population (98.3 %). When cultivating cells in 3D, the viable population decreased (38.5 %) and the number of cells in the dead gate increased drastically. We attribute this behavior to two aspects: firstly, the existence of a necrotic core in the large spheroids^[15]^, and secondly, the potential starvation of cells as spheroids had been cultured with the same 50 µl of medium for 10 days at this point. The addition of 300 fmol FasL reduced the number of viable cells to 16.9 %, while the incubation with rroOF nanoagent reduced this population even further to 4.6 %. Encouragingly, for spheroids incubated with the most potent rroONF nanoagent only a negligible viable population of 1.2 % could be observed, suggesting that rroONF was indeed able to eradicate the tumor. However, since cell morphology is only an indirect measure of cell viability, we further conducted a second set of experiments, where spheroids incubated with different nanoagents were first dissociated and subsequently re-seeded in 2D. Cultivation of cells from control spheroids showed confluency after 2 days of incubation, as shown in **Figure 4d** and S30 (this was the case for each spheroid of the triplicate). Similarly, spheroids previously incubated with soluble FasL and rroOF showed regrowth, but with fewer and smaller populations. This suggests that a small fraction of viable cells remained in the spheroid, consistent with the FACS results. Importantly, spheroids exposed to rroONF did not show any regrowth. We hypothesize that the 1.2 % of viable cells observed by FACS are either background or potentially cells undergoing apoptosis at an early stage or just outliers in size of the apoptotic population. Despite inducing similar behavior qualitatively in spheroid growth (cf. **Figure 3e**), the incubation with lower amounts of rroONF (25 fmol) could still achieve the complete eradication of viable cells in most cases, while even lower amounts did not (see **Figure 4d** and S31). This suggests that 50 fmol of ONF nanoagent are required for optimal efficacy and complete eradication of viable cells in the spheroid, which is not achievable by treatment with either the soluble FasL or the OF nanoagents.

**Figure 4:**
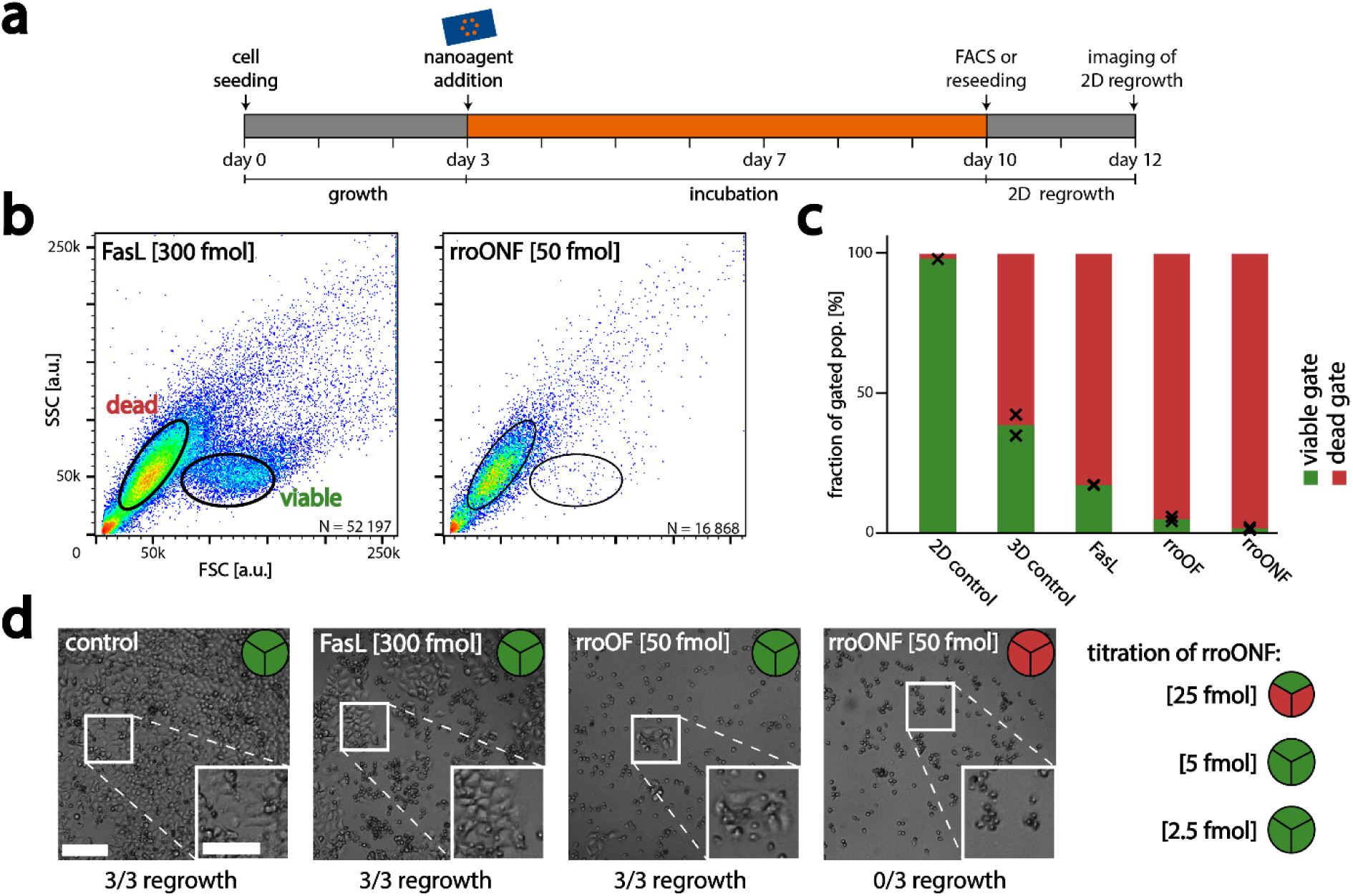
analysis of the efficacy of the apoptosis induction. (**a**) Experimental timeline: cells were seeded and grown for 3 days. Origami (blue) and/or FasL (orange) were added on the third day. The spheroids were cultivated for an additional week. After 10 days the spheroids were dissociated and either analyzed with FACS or reseeded and cultivated in 2D for two additional days. On day 12 regrowth of cells was examined. (**b**) Exemplary FACS scatter plots of cells from dissociated spheroids, incubated with FasL or rroONF nanoagent. Viable (green) and dead (red) populations differ from each other in FSC values. (**c**) Fraction of cells in viable and dead gates. Black crosses indicate the results of the single measurements. Fraction of viable cells: 2D control: 98.3 % (n=1 data set), 3D control: 38.5 % (n=2); 300 fmol FasL: 16.9 % (n=2); rroOF 4.6 % (n=2); rroONF: 1.2 % (n=2). (**d**) 2D cultures from dissociated spheroids. In 3/3 regrowth experiments (indicated with the pie chart in each image) cells survived in spheroids incubated with FasL, OF nanoagent, or low concentrations of ONF nanoagent. For 50 fmol ONF, no viable cells were observed in all three cases of spheroids. For 25 fmol ONF only cells from one out of three spheroids showed a small population of viable, adherent cell populations after 2 days of incubation. The scale bar in (**d**) for the overview images is 200 µm and for the zoom-ins is 100 µm.

## 3. Conclusion

In order to further develop nanostructured therapeutics, it is essential to understand and optimize their behavior and design for the interaction with solid tumors. We therefore investigated the penetration ability of different DNA origami nanostructures through large 3D spheroids and tested the ability of DNA origami – FasL nanoagents to diminish viable cells in the spheroids and thus eradicate them. We conclude two findings: Firstly, the ability of DNA origami to penetrate spheroid tissue is governed by their size rather than their internal structure and flexibility. Secondly, the apoptosis induction efficiency of origami - FasL nanoagents towards spheroids is dictated by the attachment strategy of FasL to the DNA origami rather than the design of the underlying origami.

When analyzing the penetration speed of different DNA origami through spheroid tissue, we found equally slow penetration for the rro in the square lattice design and the much more flexible, but similarly-sized wf origami in the wireframe design. However, the much smaller mini origami penetrated ∼3.5 times faster through the spheroid tissue. This could potentially be explained by Brownian motion being negatively proportional to the particle’s hydrodynamic radius, and the smaller size allowing for better penetration through the cell-cell junctions. However, our findings are somewhat contradictory to a prior study^[18]^, where a better penetration of more structurally flexible wireframe rods compared to DNA origami rods built in a honeycomb lattice, was observed. Taken together, we can now say that flexibility may play a small role in penetration ability, but is greatly outweighed by origami size.

Despite this faster and better penetration of small origami structures through spheroids, it surprisingly did not affect their efficacy as apoptosis-inducing nanoagents, formed via the attachment and precise spationumerical placement of FasL on the DNA origami structures. All three tested DNA origami structures were able to halt tumor growth. If FasL was attached via a rigid neutravidin-biotin linker, rroONF nanoagents were able to eradicate any viable cells in the spheroids. However, if a more flexible dsDNA linker (of ∼7 nm length) was used, nanoagents were far less efficient, unable to halt tumor growth or fully eradicate viable cells in the spheroids, even though both attachment strategies showed comparable efficiencies of FasL conjugation to the origami (ONF: 76 %^[13a]^, OF: 71 %^[25]^). Although we had previously observed a reduced efficiency of FasL nanoagents being attached via more flexible linkers in 2D cell culture^[13a]^, this effect was much more pronounced in 3D. We attribute this fact mostly to the higher flexibility of the dsDNA linker and the subsequent higher positional inaccuracy of FasL placement. A difference in the accessibility of FasL can be excluded as both the dsDNA linker and the neutravidin place the ligand at a similar distance (height) from the origami surface.

Similarly, an increased “sticking” of the positively charged FasL to the origami, owing to a more flexible dsDNA linker can be excluded as during TEM imaging of the miniOF structure, most proteins appeared next to the origami rather than directly on it. We surmise further, that the amount of flexibility added through the dsDNA connection far exceeds flexibility originating from the DNA origami structure, as we saw the same halting behavior for the more flexible wfONF nanoagents as for the rro and miniONF.

In this study, we added another dimension to the design of DNA origami-based drug development. We first provided new insight into the effects of origami design on their ability to move through tumor tissue. We then found that minute differences in nanoagent design, namely the attachment strategy of FasL to the origami, exhibit major differences in the response of the complex spheroid system. All in all our findings constitute an important further step in the future development of nanomaterial-based drugs and therapy approaches.

## 4. Methods

### 4.1 Cell Culture

HeLa Apo12 mGFP cells, used in reference^[13a]^ were cultivated in Dulbecco’s modified eagle medium (DMEM) Glutamax (gibco, cat.no.: 31966), supplemented with 10 % (v/v) fetal bovine serum (FBS) (gibco, cat.no.: 10270), and 1 % (v/v) penicillin/ streptomycin (PenStrep) (gibco, cat.no.: 15140). Cells were split into 500 000 cells or 200 000 cells in 10 ml new medium to reach ∼70 % confluency after two or 3 days of incubation, respectively. Both, 2D and 3D cultures were incubated at 37 °C, with ∼100 % humidity and 5 % CO_2_. Only cells of passage numbers 5 to 20 were used.

Spheroids were produced by seeding 800 cells in DMEM 10 % FBS 1 % PenStrep in 96 well low adhesion plates (Nunclon Sphera 96-well, cat.no.:174929) and centrifuged for 3 min at 1000 rcf to cluster cells. Spheroids were then incubated for 3 days before the nanoagents were added.

For penetration experiments, spheroids were grown as described above and starting from day 3 incubated with 500 fmol of the respective DNA origami structure with overall MgCl_2_ concentration adjusted to 5 mM. For fixation, they were washed once with phosphate-buffered saline (PBS) and then incubated with 50 µl 4 % paraformaldehyde (FA) solution for 30 min at 37 °C. After three washing steps with PBS, the spheroids were permeabilized with 50 µl PBS, 2% tween, 2 % PBST (0.05 % (w/v) NaAc, 2 % (v/v) Triton X 100 in 1X PBS) for 15 min at 37 °C. After washing three times with PBS, they were washed once with “hybridization buffer” (Molecular Instruments), and then 50 µl hairpin solution was added and incubated o/n. For the hairpin solution FISH hairpins (B1, Molecular Instruments) were heated 90 s at 95 °C and then cooled to room temperature (RT), and then diluted 100 X in hybridization buffer. After overnight incubation the spheroids were washed thrice with “wash buffer” (Molecular Instruments). Here, optionally, the spheroids were stained with 0.5 % Hoechst 33342 (ThermoFisher Scientific) for 20 min at RT and then washed once with PBS. On the following day, the spheroids were washed with probe “wash buffer” (Molecular Instruments) and then transferred to 15 well glass bottom slides (ibidi, ct.no. 81507). Excess buffer was removed and the spheroids were air-dried for 5 min to secure them into place. To optically clear the TS, 8-10 µl of RapiClear 1.47 (SunJin Biolab, cat.no.: #RC147001) was added and the spheroids were then imaged on a Leica Stellaris 8. Optionally, the brightness was adjusted for better visualization (information added in figure captions if applicable) in Fiji^[27]^ version 2.9.0 or later and the clockscan plugin^[24]^.

For spheroid-killing experiments, spheroids were grown as described above and on day 3 the respective nanoagent was added. Optionally, 1 µl of AnxV staining solution (Invitrogen, cat.no.: A13203) was added to illustrate apoptosis events. Images were taken every day on an Evos FL Auto 2 for one week. Data was extracted and analyzed with a custom Fiji script.

For reseeding experiments, spheroids were washed twice with PBS and then incubated for 30 min with 50 µl Trypsin ethylenediaminetetraacetic acid (EDTA) (TE). Spheroids were then dissociated by mechanical stimulus and reseeded into 200 µl DMEM 10 % FBS 1 % PenStrep and let grow for 2 days. Imaged on an Evos FL Auto 2.

For fluorescence-activated cell sorting (FACS) experiments, spheroids were washed twice with PBS and then incubated for 30 min with 50 µl TE. Spheroids were then dissociated by mechanical stimulus and centrifuged for 3 min at 1000 rcf to sediment. Cells were then resuspended in PBS for FACS imaging.

### 4.2 DNA origami

DNA origami were designed with either caDNAno version 2.4.10^[28]^ or vHelix^[21]^. Afterwards, they were simulated in oxDNA^[22]^ version 3.5.0 with 100 000 000 iteration steps. Staples were ordered from IDT and p7249 was produced as described previously^[29]^, p4844 was ordered from tilibit, and p1033 was produced as described previously^[20]^.

DNA Origami were synthesized with either the p7249 (RRO), p4844 (wireframe), or the p1033 (mini) scaffold. Structural staples were used in 4X excess and staples acting as handles were added in at least 8X excess, staple sequences are found in tables S1-S3. Origami were folded by heating to 65°C, holding for 5 min, then cooling down over 16 hours to 20°C and purified with ultracentrifugation (one equilibration round, two sample application rounds, and 4 wash rounds at 8000 rcf for 4 min and sample recovery for 2 min at 5000 rcf). The origami were stored at −20 °C until further use.

For transmission electron microscopy (TEM) characterization, carbon grids (Plano GmbH, cat.no.: S162-3) were first treated with an oxygen plasma for 30 s, then 10 µl 2 nM sample was incubated for 5 min, and then stained with 2 % uranyl formate for 10 s. Samples were imaged on a Jeol-JEM-1230 at 80 KeV. TEM micrographs were analyzed with Fiji^[27]^ version 2.9.0 or later, and the contrast was adjusted for better visibility.

Agarose gel electrophoresis was performed with 1 % agarose gels, TAE running buffer (40 mM Tris, 20 mM acetic acid, 1 mM EDTA, 11 mM MgCl_2_, pH. 8.0) precast with SYBR safe nucleic acid stain (Invitrogen, cat.no.: S33102) for 90 min at constant 70 V on ice. The gel was scanned with a Typhoon FLA 9000 laser scanner and analyzed with Fiji version 2.14.0.

### 4.3 FasL functionalization

Potential disulfide bridges on FasL were reduced by the addition of tris(2-carboxyethyl)phosphine (TCEP) to a total concentration of 2 mM for 30 min. FasL was then functionalized by the addition of either a maleimide functionalized DNA oligomer (biomers) or a Maleimide-C6-Biotin (Sigma-Aldrich cat.no.: B1267) in 5X or 30X excess over the trimerized FasL, respectively, and incubated over-night. Functionalized FasL was purified by 10 rounds of ultracentrifugation (with one round of equilibration, one round of sample application, and ∼10 rounds of washing with 1X PBS 8 000 rcf for 4 min, and sample recovery at 5000 rcf for 2 min)

Concentrations were determined by nanodrop absorption curves, either by the absorbance at 280 nm (A280) value, or the ssDNA33 value under the assumption of three attached ssDNA handles due to the trimerized FasL structure. FasL was then either frozen in liquid N_2_ with the addition of glycerol to 10 % or directly used to functionalize DNA Origami. FasL used for incubation was filtered through a 0.22 µm mesh (Merck Millipore, cat.no.: UFC30GV, at 5000 rcf for 2 min) to sterilize the sample before freezing with liquid N_2_ and storing at −80 °C.

### 4.4 Nanoagent synthesis

For ONF nanoagents, the neutravidin (Thermo Scientific, cat.no.: 31000) was added in 50X excess over each binding site and incubated at 4 °C o/n. To remove potential aggregates, the sample was the next day filtered through a 0.22 µm mesh, as described above, and then purified with High-performance liquid chromatography (HPLC), similar to reference^[30]^. The sample was then concentrated using ultrafiltration.

For all nanoagents, the FasL was added in 5X excess over each binding site and incubated at 4 °C o/n. To remove potential aggregates, on the next day, the sample was filtered through a 0.22 µm mesh, as described above, and then purified with HPLC, similar to reference^[30]^. The sample was then concentrated using ultrafiltration. Then glycerol was added to a total concentration of 10 %. To sterilize the sample, it was filtered through a 0.22 µm mesh, as described above, and then frozen in liquid N_2_.

## Supporting information

Supporting Information is available from the Wiley Online Library or from the author.

## Data availability

All data supporting the key findings of this study are available within the main text and supplementary information files. Additional data used in this study are available from the corresponding author upon reasonable request.

## Author contributions

J.M.W. performed the research, supported by R.N. and M.Z.K.. X.S. and C.M. designed and created the trimerized FasL and provided the HeLa Apo12 mGFP cell line. J.M.W. and A.H.-J. wrote the manuscript. J.M.W. designed the figures. J.M.W. and A.H.-J. designed the experiments. A.H.-J. conceived the study and provided the funding. All authors discussed and edited the final manuscript draft.

## Competing interests

The authors declare no conflict of interest.

## Acknowledgments

A.H-J acknowledges financial support from the German Research Foundation (DFG) through SFB1032 (Nanoagents) project A06 and the Emmy Noether program (project no. 427981116). CM was supported by the German Research Foundation (DFG) – project number 267205415 - CRC 1208, project A12 and project number 458090666 - CRC 1535, project A09 as well as the Freigeist Fellowship by the Volkswagen Foundation.

We thank S. Suppman, J. Basquin, Y. Xiao, and J. Scholz for their help with protein expression. We thank M. Pinner, M. N. Honemann, and H. Dietz for providing the p1033 scaffold. We thank M. Braun and U. Weber for their help with TEM imaging.

## Notes

### Competing Interest Statement

The authors have declared no competing interest.

