## Supplementary Files for "Effects of DNA Origami-Based Nanoagent Design on Apoptosis Induction in a Large 3D Spheroid Model"

**Table of figures**

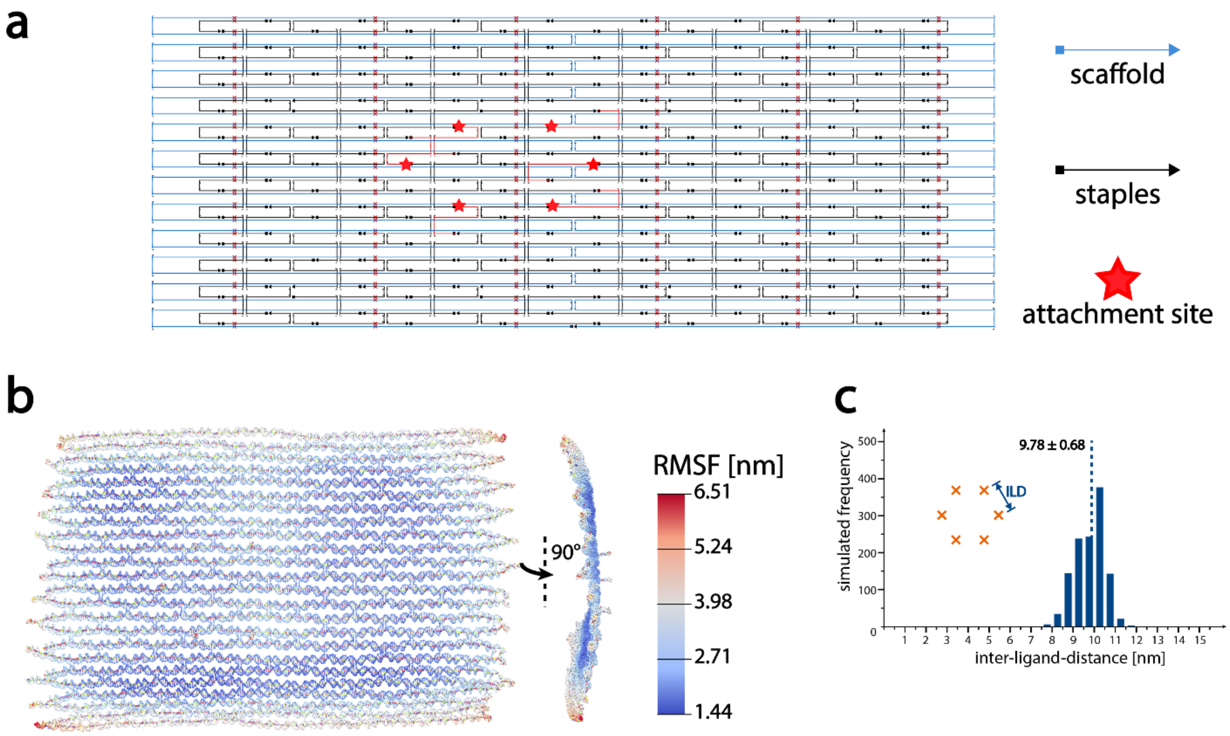

**Figure S1: design and simulation of the rro DNA origami** (**a**) caDNAno screenshot of the rro layout: blue lines indicate scaffold routing, black lines indicate staple routings and red stars indicate attachment sites. (**b**) oxDNA simulation of the rro: front and side view of the average structure, indicated by a heatmap is the RMSF of the structure. In the oxDNA simulation, the dimensions of the rro DNA origami are approximately 90 nm x 59 nm. (**c**) In-silico analysis of ILD on the respective DNA origami. n=1200 ILDs at different time points in the simulation for each structure. The ILD extracted from the simulation of the rro origami is 9.78 ± 0.68 nm.

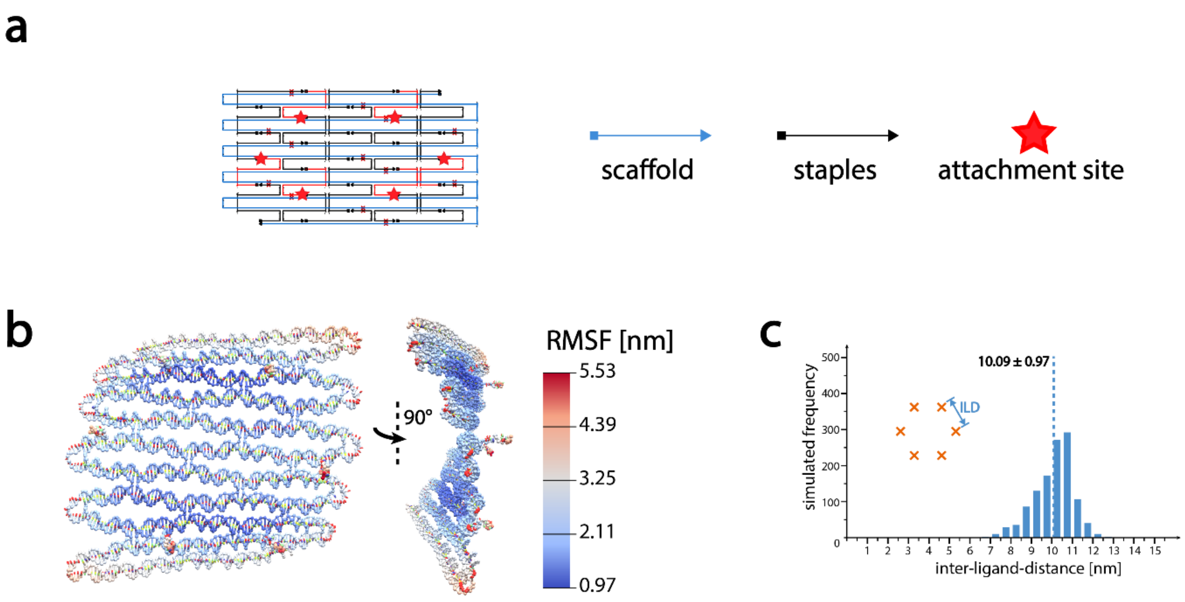

**Figure S2: design and simulation of the mini DNA origami** (**a**) caDNAno screenshot of the mini layout: blue lines indicate scaffold routing, black lines indicate staple routings and red stars indicate attachment sites. (**b**) oxDNA simulation of the mini: front and side view of the average structure, indicated by a heatmap is the RMSF of the structure. In the oxDNA simulation, the dimensions of the mini DNA origami are approximately 28 nm x 24 nm. (**c**) In-silico analysis of ILD on the respective DNA origami. n=1200 ILDs at different time points in the simulation for each structure. The ILD extracted from the simulation of the mini origami is 10.09 ± 0.97 nm.

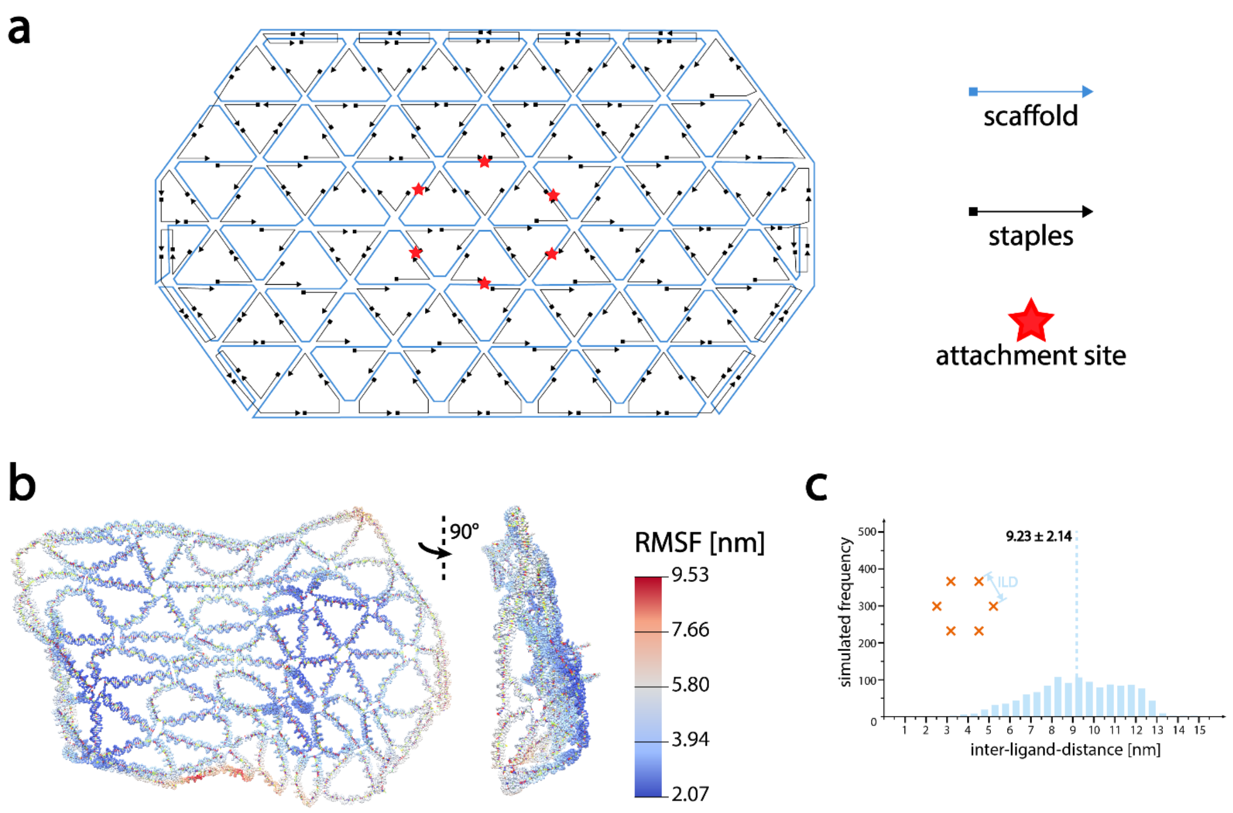

**Figure S3: design and simulation of the wf DNA origami** (**a**) scaffold and staple layout of the wf layout: blue lines indicate scaffold routing, black lines indicate staple routings and red stars indicate attachment sites. (**b**) oxDNA simulation of the wf: front and side view of the average structure, indicated by a heatmap is the RMSF of the structure. In the oxDNA simulation, the dimensions of the wf DNA origami are approximately 67 nm x 45 nm. (**c**) In-silico analysis of ILD on the respective DNA origami. n=1200 ILDs at different time points in the simulation for each structure. The ILD extracted from the simulation of the wf origami is 9.23 ± 2.14 nm.

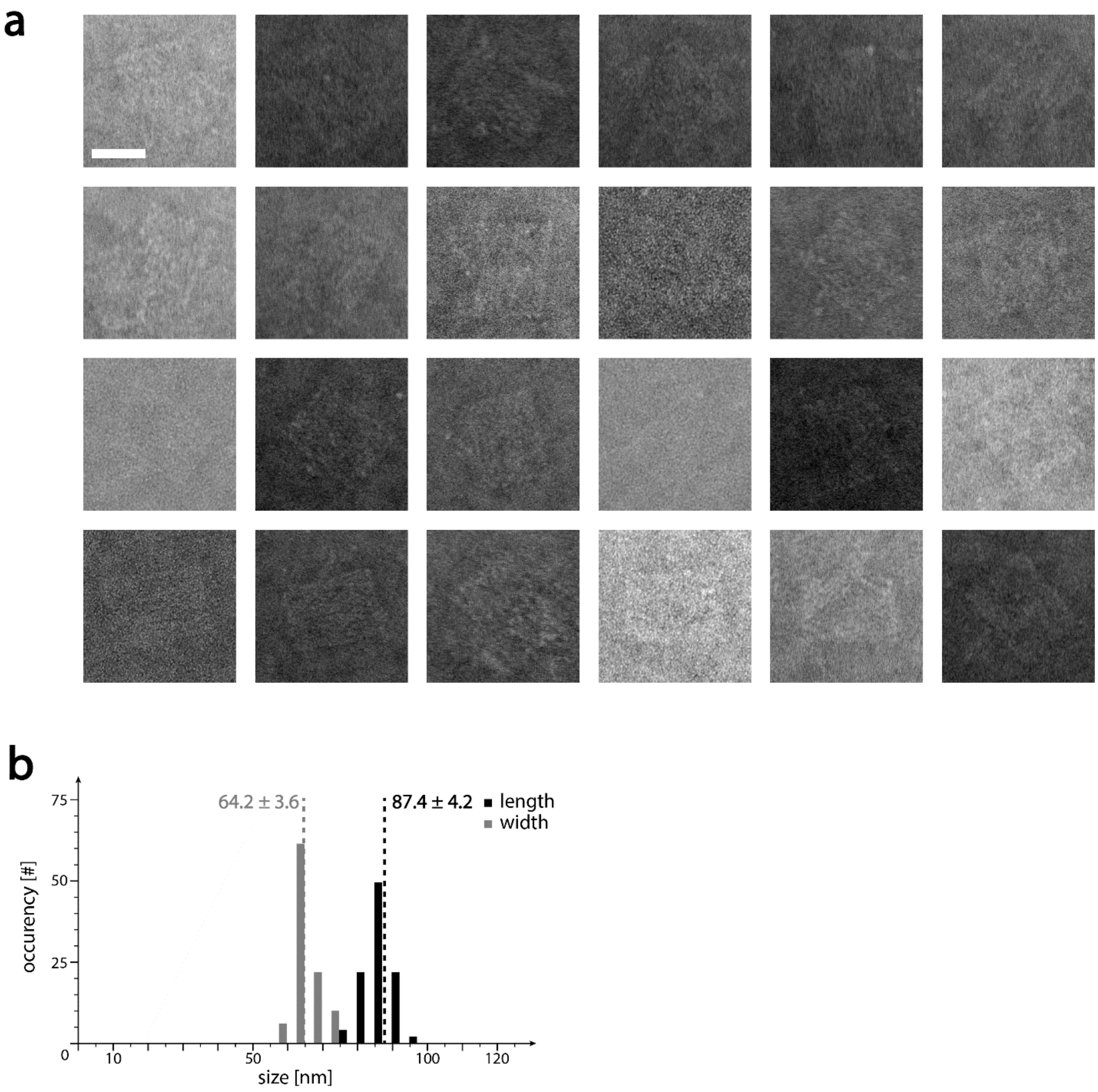

**Figure S4: Transmission electron microscopy (TEM) characterization of the rro DNA origami**: (**a**) Cropped micrographs show single rro origami. (b) Histograms of length (black) and width (grey) of rro origami extracted from TEM micrographs. N>30 structures, normalized to 100. Length is 87.4 ± 4.2 nm and width is 64.2 ± 3.6 nm. Images from the same experiment were used in Figure 1b. The scale bar in (**a**) is 50 nm and holds for all micrographs.

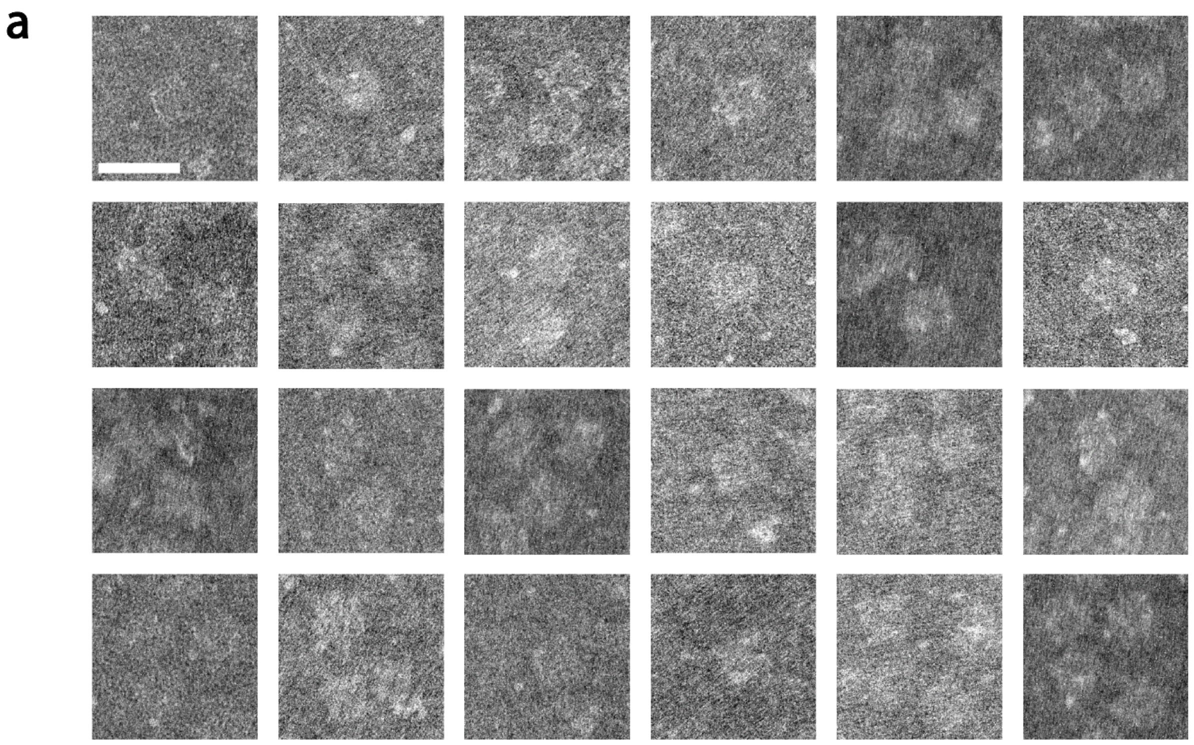

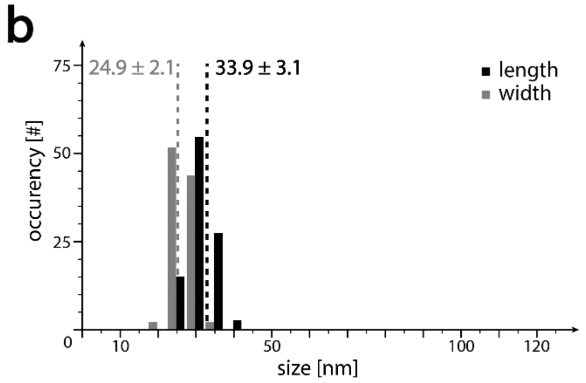

**Figure S5: TEM characterization of the mini DNA origami**: (**a**) Cropped micrographs show mini origami. (b) Histograms of length (black) and width (grey) of mini origami extracted from TEM micrographs. N>30 structures, normalized to 100. Length is 33.9 ± 3.1 nm and width is 24.9 ± 2.1 nm. Images from the same experiment were used in Figure 1b. The scale bar in (**a**) is 50 nm and holds for all micrographs.

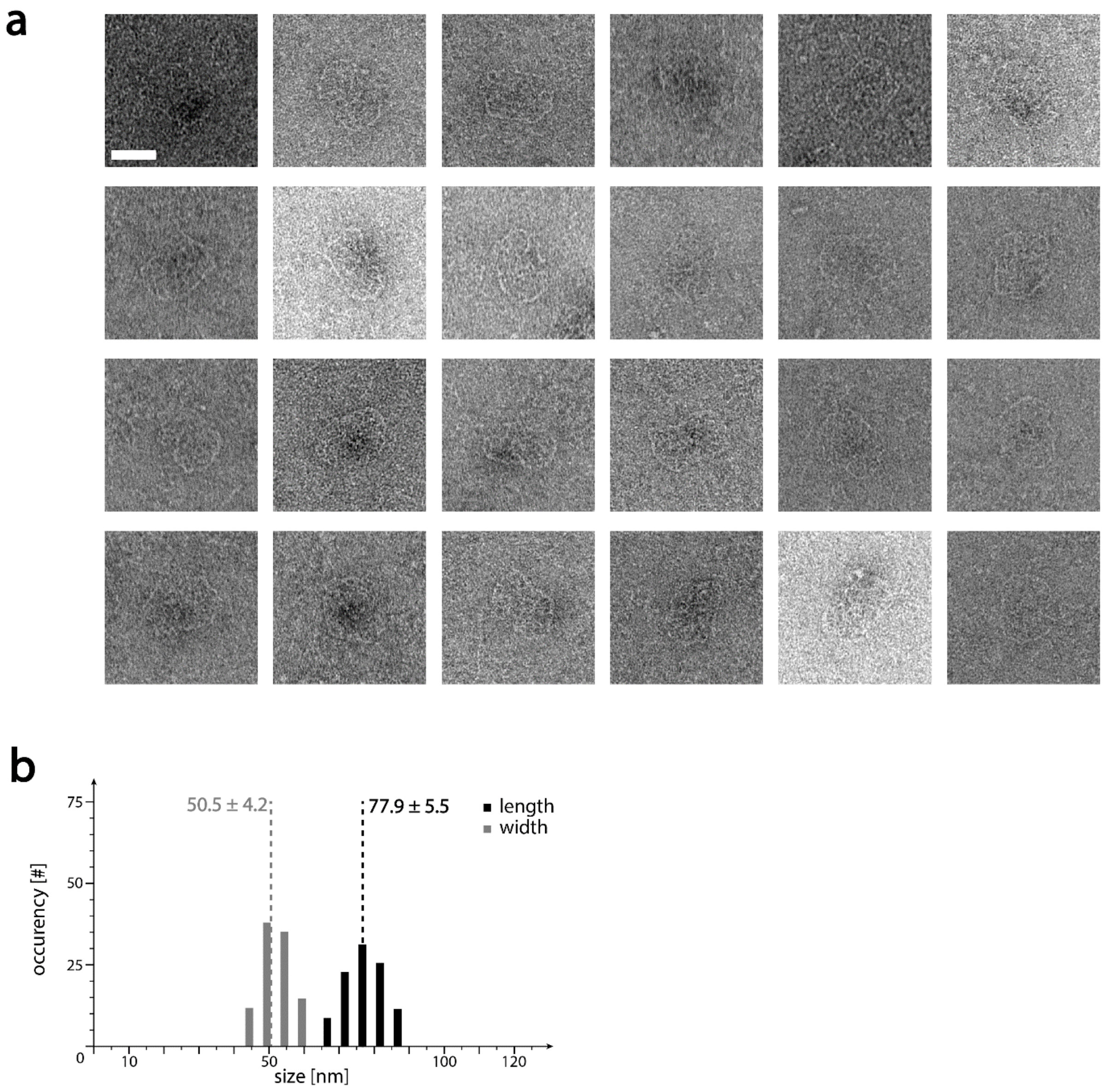

**Figure S6: TEM characterization of the wf DNA origami**: (**a**) Cropped micrographs show single wf origami. (b) Histograms of length (black) and width (grey) of wf origami extracted from TEM micrographs. N>30 structures, normalized to 100. Length is 77.9 ± 5.5 nm and width is 50.5 ± 4.2 nm. Images from the same experiment were used in Figure 1b. The scale bar in (**a**) is 50 nm and holds for all micrographs.

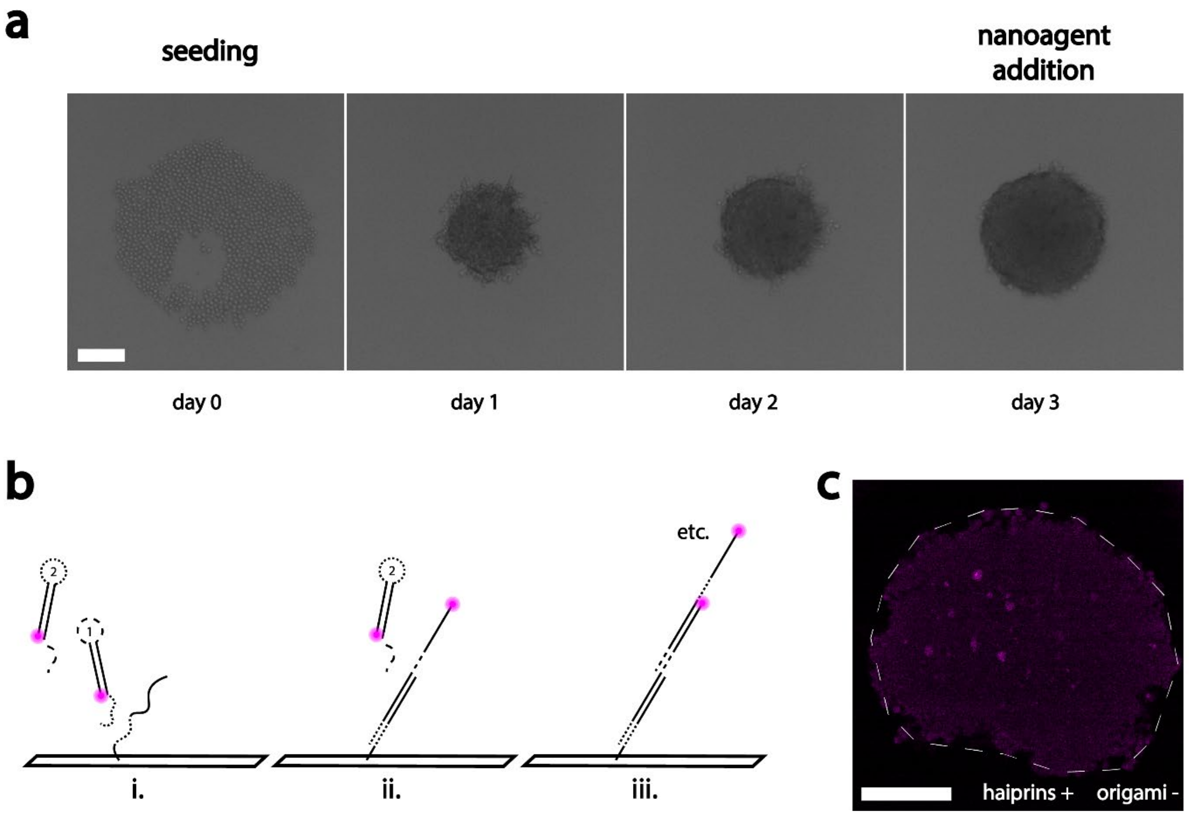

**Figure S7: Spheroid seeding and origami FISH** (**a**) first three days in the spheroid development. On day 0 the cells were seeded and in a flat layer on the low adhesion plate. In the following three days, the cells proliferate and form a spheroid with sharp contours. (**b**) Schematic of origami FISH: staples extended with an anchor sequence are protruding from a DNA origami. (i.) To a toehold region (dotted part) on the anchor sequence, hairpin 1 hybridizes and displaces the stem-loop, making a toehold (stripes) and an attachment site for hairpin 2 accessible. (ii.) Displacement of the stem-loop in hairpin 2, again makes a toehold and an attachment site for hairpin 1 accessible, starting a chain reaction, which leads to (iii.) an accumulation of fluorophores around the initial attachment site. (**c**) Control spheroid without DNA origami, but with fluorescent hairpins shows a very dim, homogeneous fluorescence signal throughout the entire spheroid. The contrast was adjusted for better visibility. The scale bar in (**a**) is 100 µm and in (**c**) is 500 µm.

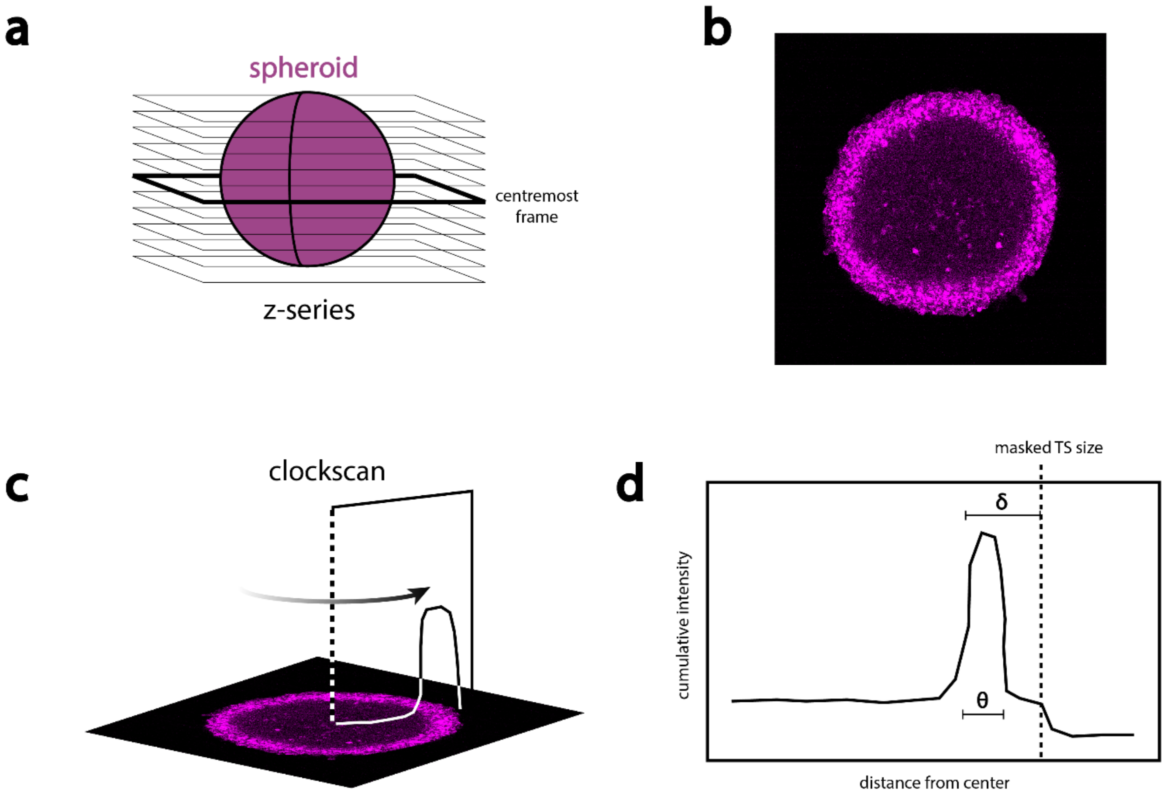

**Figure S8: Schematic of the clockscan protocol** used to extract the penetration depth from fluorescence images. (**a**) A z-stack of a cleared spheroid was recorded with a confocal microscope and the thickest part on the centermost frame (**b**) was determined. On the thickest part of the spheroid a clockscan (**c**) was performed, averaging the fluorescence signal from the center towards the outside. The fluorescence intensity was then plotted (**d**) and the penetration depth and the ring thickness were extracted.

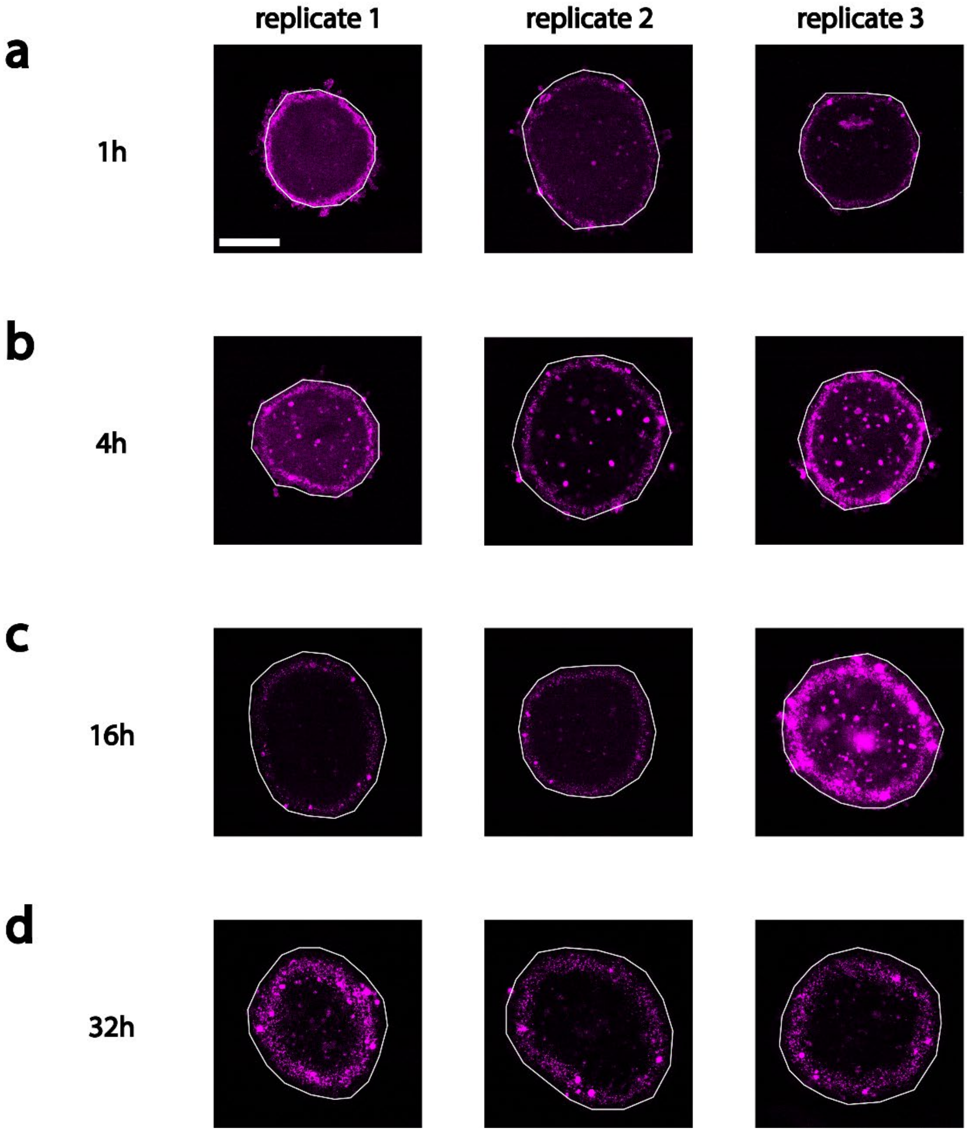

**Figure S9: rro origami penetration through spheroids** after (**a**) 1 hour, (**b**) 4 hours, (**c**) 16 hours, and (**d**) 32 hours, as triplicates of the same condition. Spheroid outlines, extracted from the GFP channel, are indicated as white lines. Graphs in Figure 2e are based on these images. The contrast was adjusted in some images for better visibility of the fluorescent rings. The scale bar is 200 µm and holds for all images.

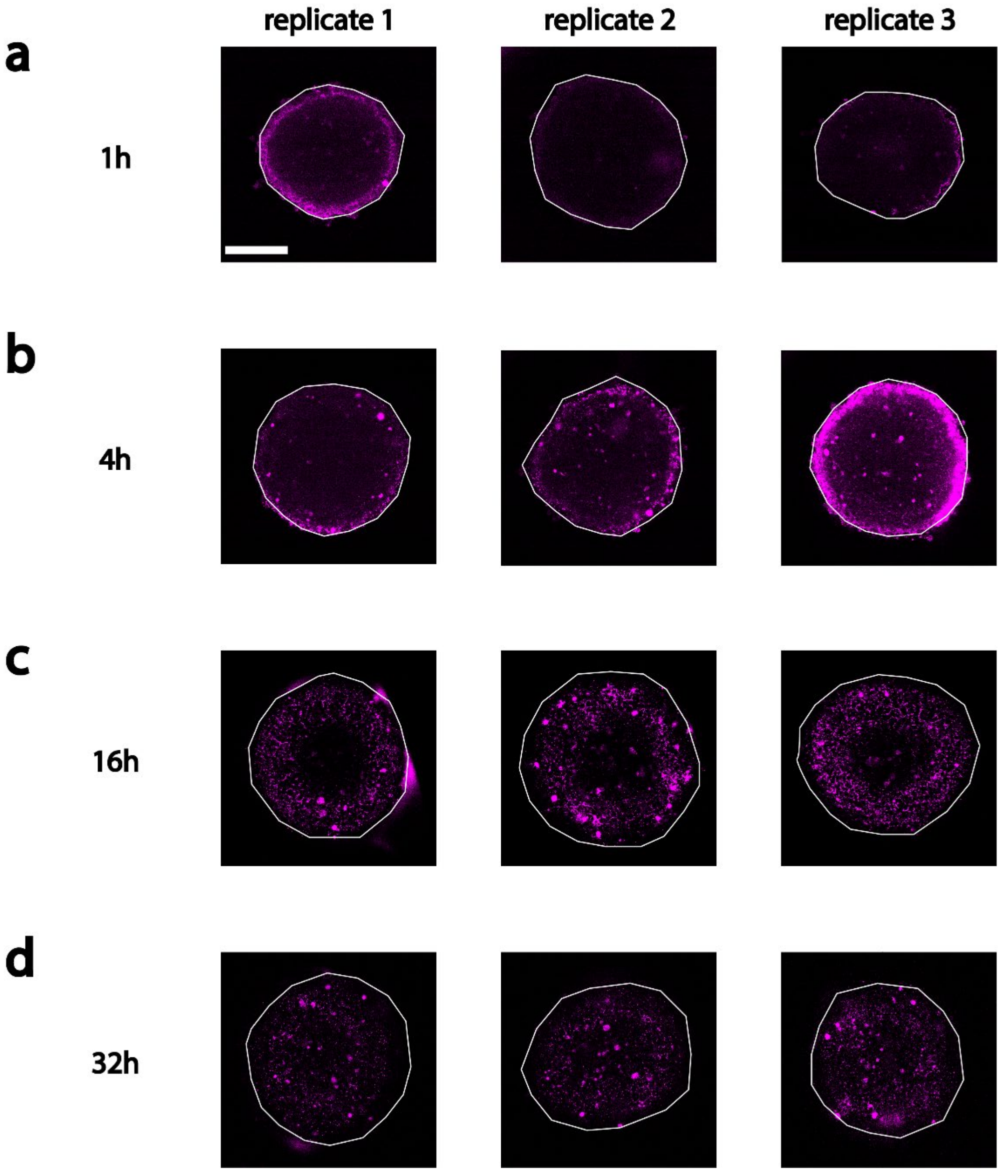

**Figure S10: mini origami penetration through spheroids** after (**a**) 1 hour, (**b**) 4 hours, (**c**) 16 hours, and (**d**) 32 hours, as triplicates of the same condition. Spheroid outlines, extracted from the GFP channel, are indicated as white lines. Graphs in Figure 2e are based on these images. The contrast was adjusted in some images for better visibility of the fluorescent rings. The scale bar is 200 µm and holds for all images.

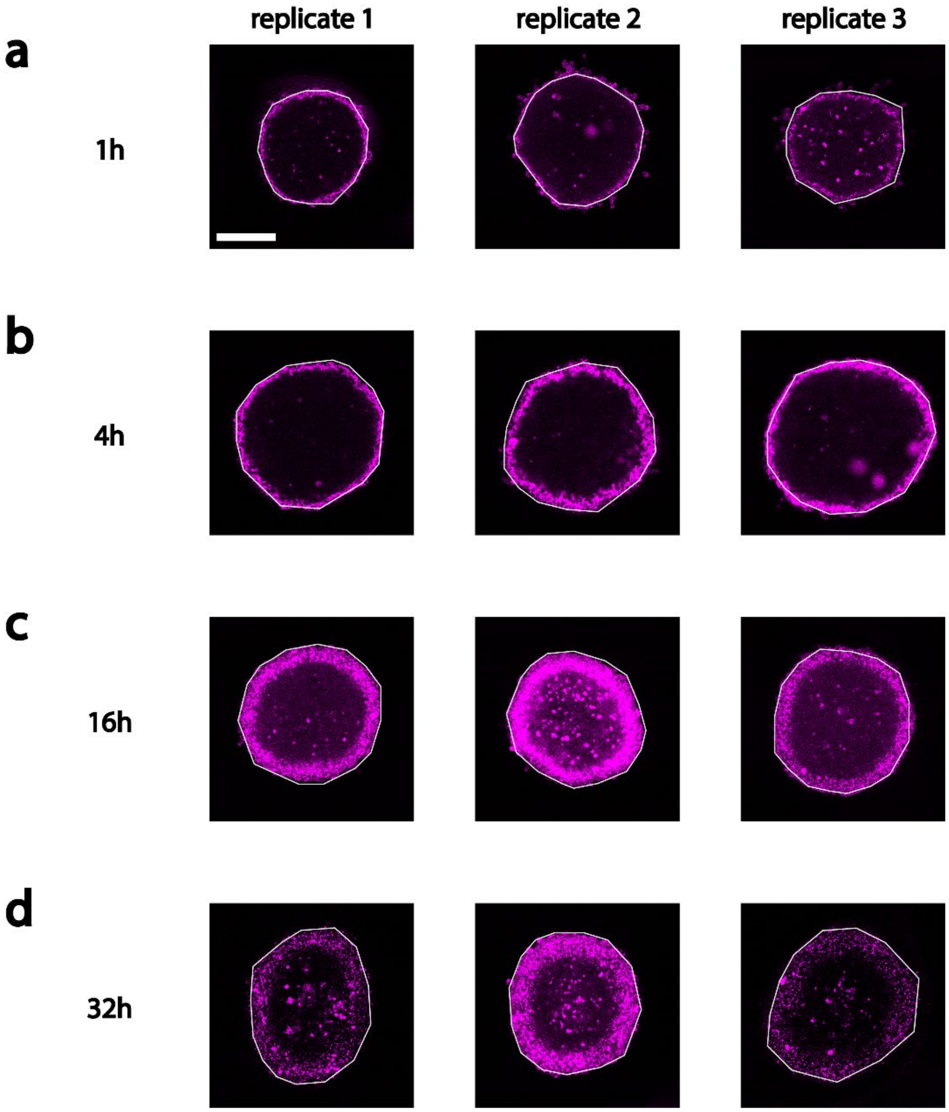

**Figure S11: wf origami penetration through spheroids** after (**a**) 1 hour, (**b**) 4 hours, (**c**) 16 hours, and (**d**) 32 hours, as triplicates of the same condition. Spheroid outlines, extracted from the GFP channel, are indicated as white lines. Images are partially used in Figure 2d. Graphs in Figure 2e are based on these images. The contrast was adjusted in some images for better visibility of the fluorescent rings. The scale bar is 200 µm and holds for all images.

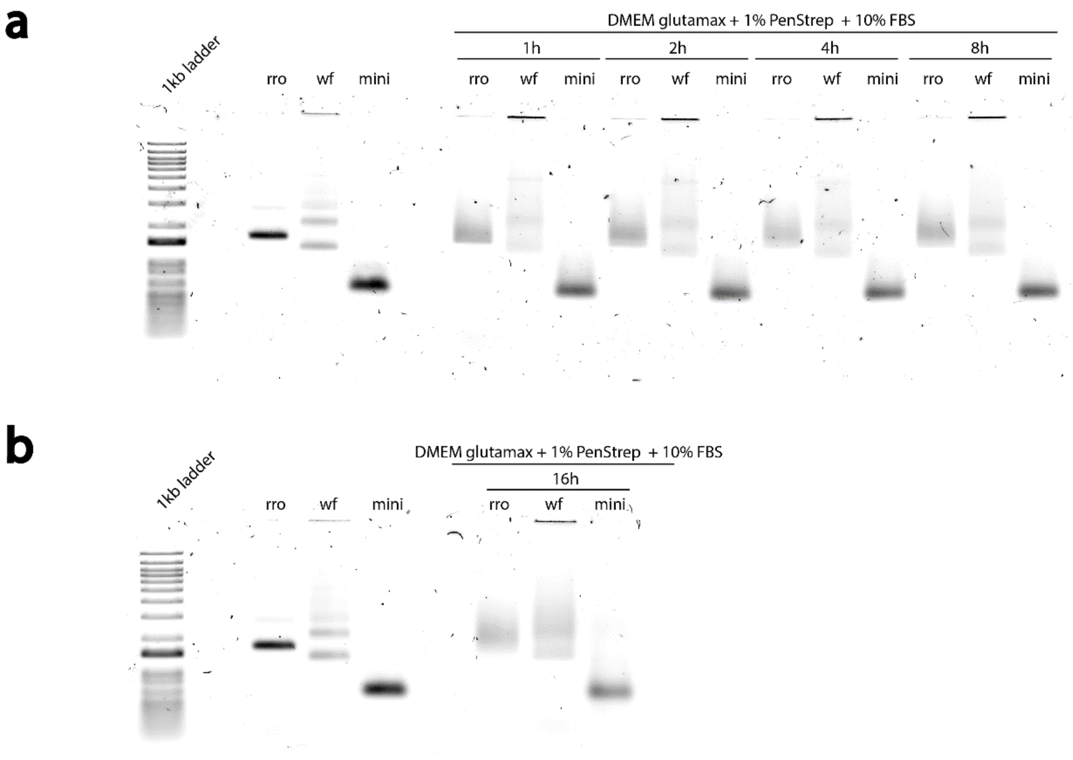

**Figure S12: stability test of DNA origami in serum-containing medium using agarose gel electrophoresis.** The three types of DNA origami (rro, mini, wf) were exposed to cell culture medium (Dulbecco's modified eagle medium (DMEM) gluatamax, supplemented with penicillin/ streptomycin (PenStrep) to 1 % and fetal bovine serum (FBS) to 10 %) for (**a**) 1 to 8 hours and (**b**) 16 hours at 37 °C. The DNA origami migrate slower with increasing structure size. Incubation in cell culture medium led to an increase in smear and, in case of wf origami an increase of aggregates. 200 ng of each DNA origami were used, respectively.

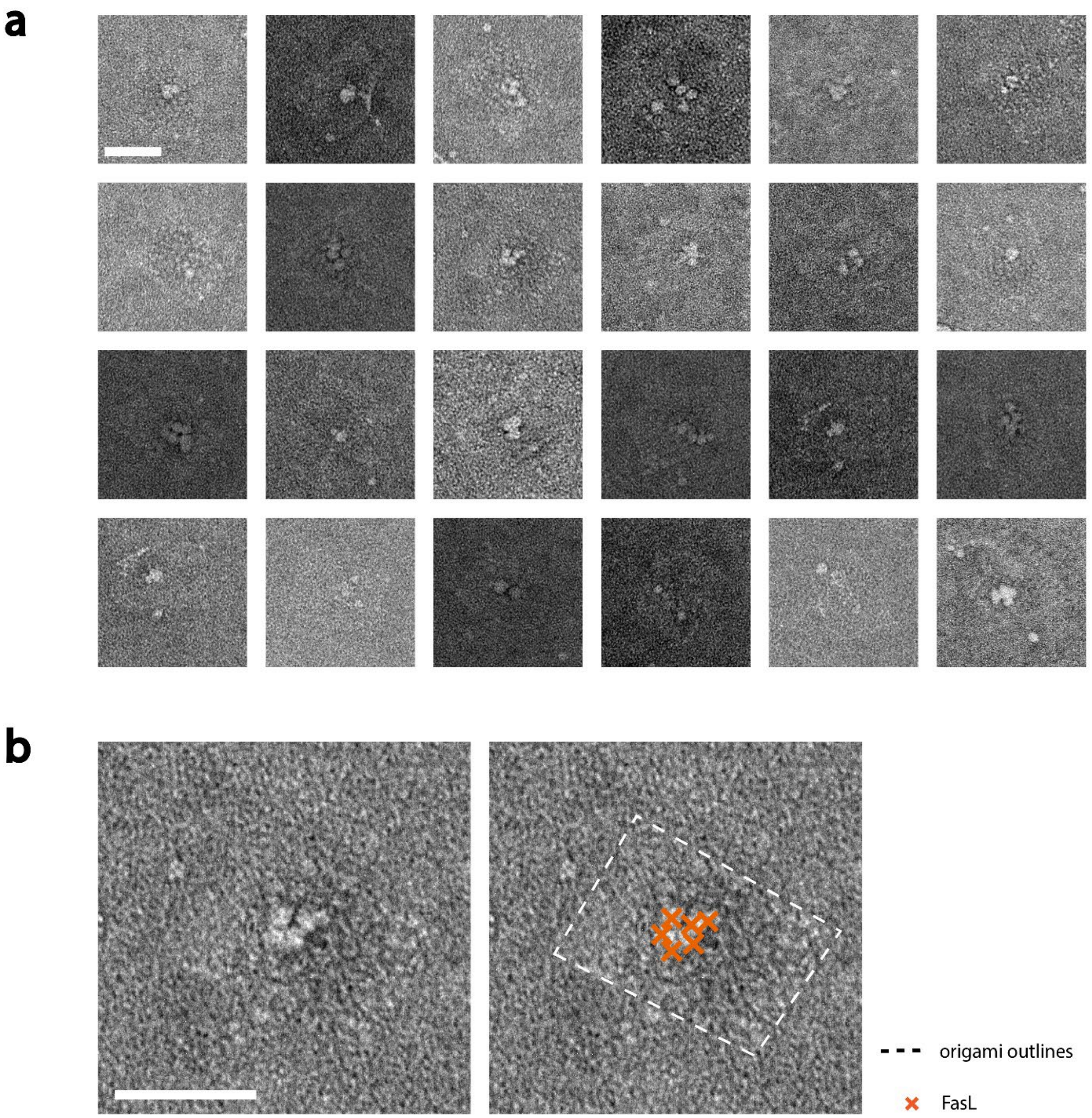

**Figure S13: TEM characterization of the rroOF nanoagents**. (**a**) Cropped micrographs show single rro DNA origami with FasL attached to them. FasL proteins are identifiable as white spots. (**b**) Zoom-in of one rroOF nanoagent, duplicate on the right with DNA origami outlines and proteins marked. The attachment efficiency of FasL to the nanoagent was determined to be 71 % in a previous publication[1]. The scale bars are 50 nm and hold for all micrographs of the respective subfigure.

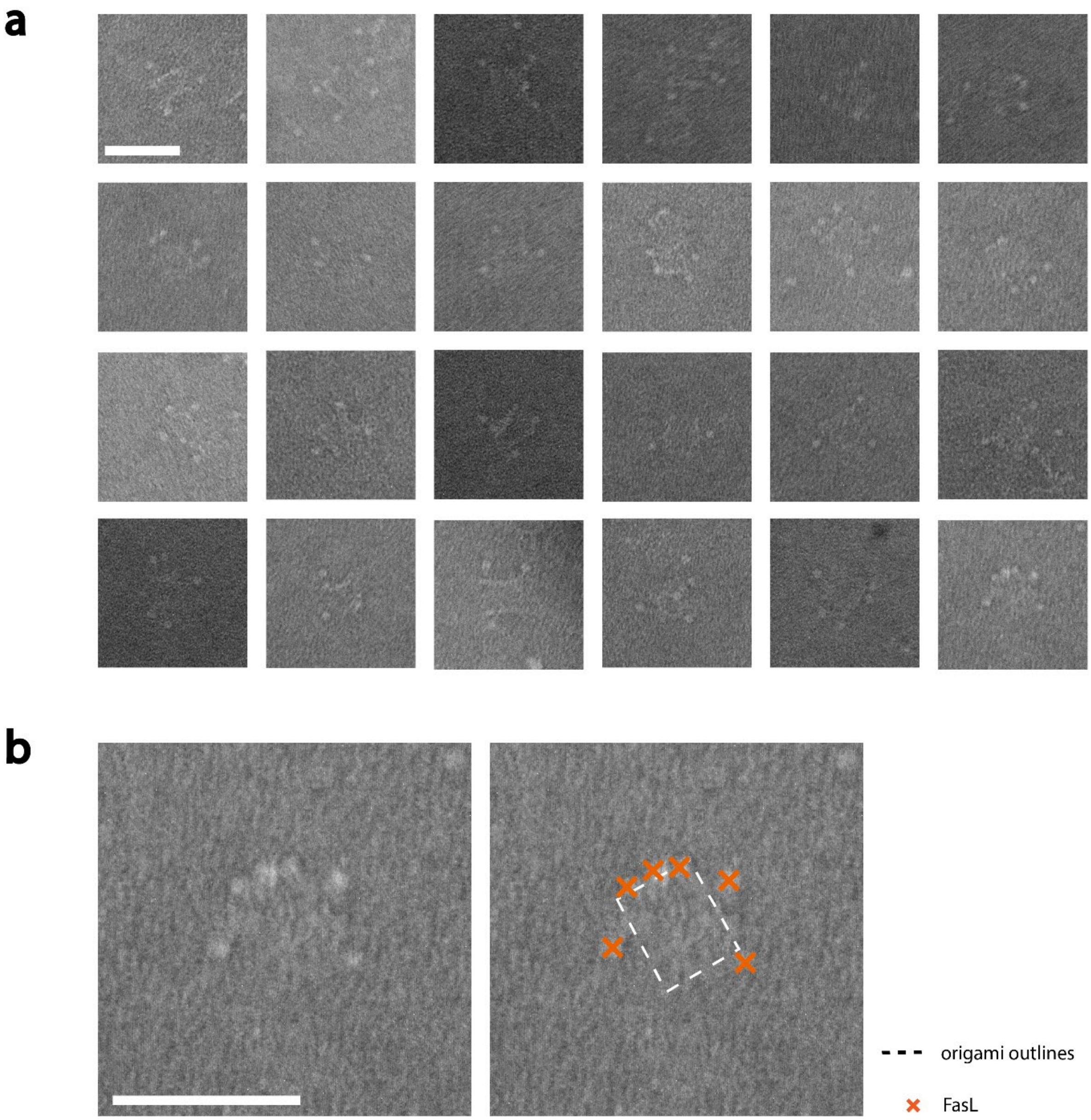

**Figure S14: TEM characterization of the miniOF nanoagents**. (**a**) Cropped micrographs show single mini DNA origami with FasL attached to them. FasL proteins are identifiable as white spots. (**b**) Zoom-in of one miniOF nanoagent, duplicate on the right with DNA origami outlines and proteins marked. The attachment efficiency of FasL to the nanoagent was determined to be 71 % in a previous publication[1]. The scale bars are 50 nm and hold for all micrographs of the respective subfigure.

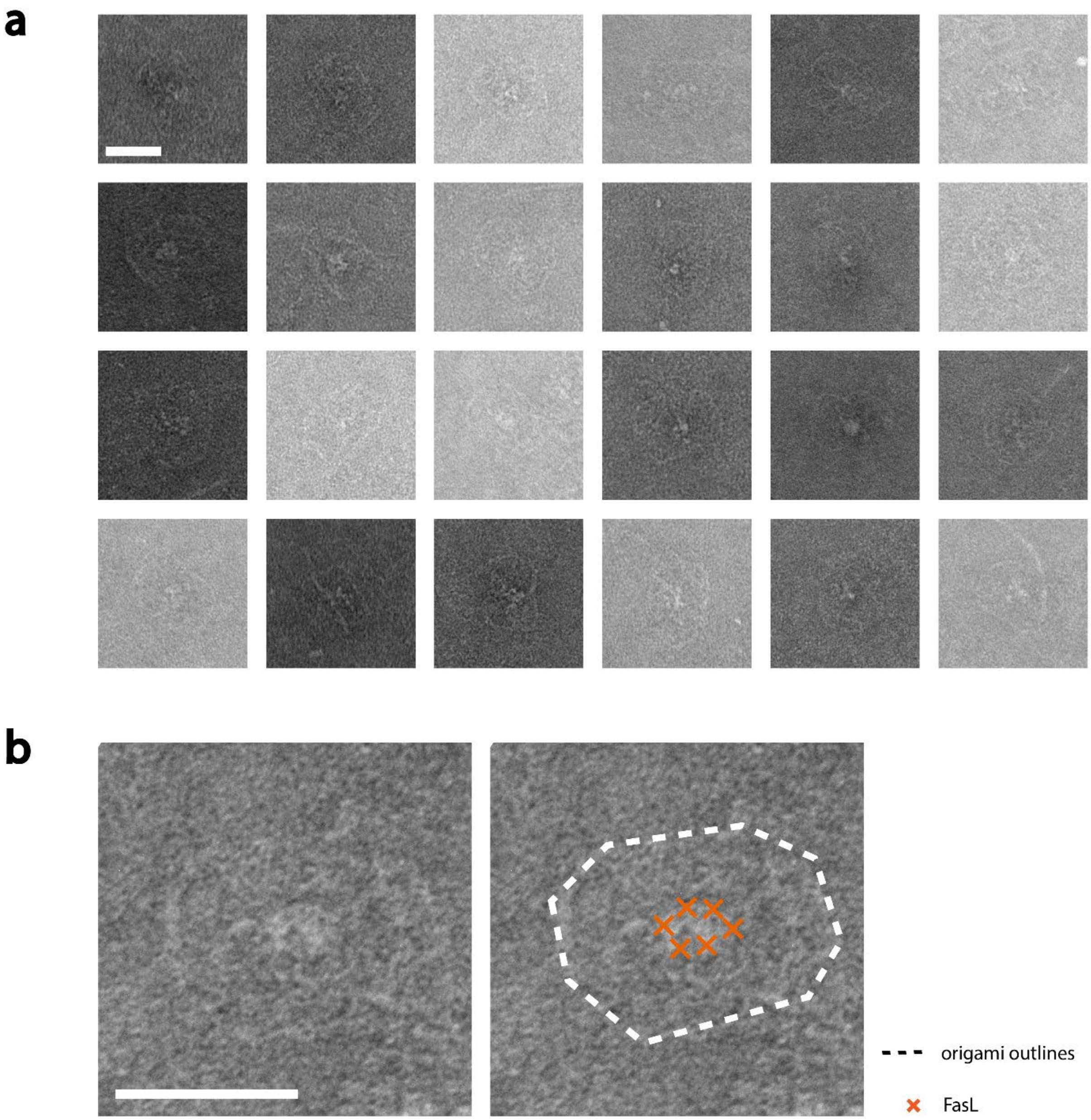

**Figure S15: TEM characterization of the wfOF nanoagents:** (**a**) Cropped micrographs show single wf DNA origami with FasL attached to them. FasL proteins are identifiable as white spots. (**b**) Zoom-in of one wfOF nanoagent, duplicate on the right with DNA origami outlines and proteins marked. The attachment efficiency of FasL to the nanoagent was determined to be 71 % in a previous publication[1]. The scale bars are 50 nm and hold for all micrographs of the respective subfigure.

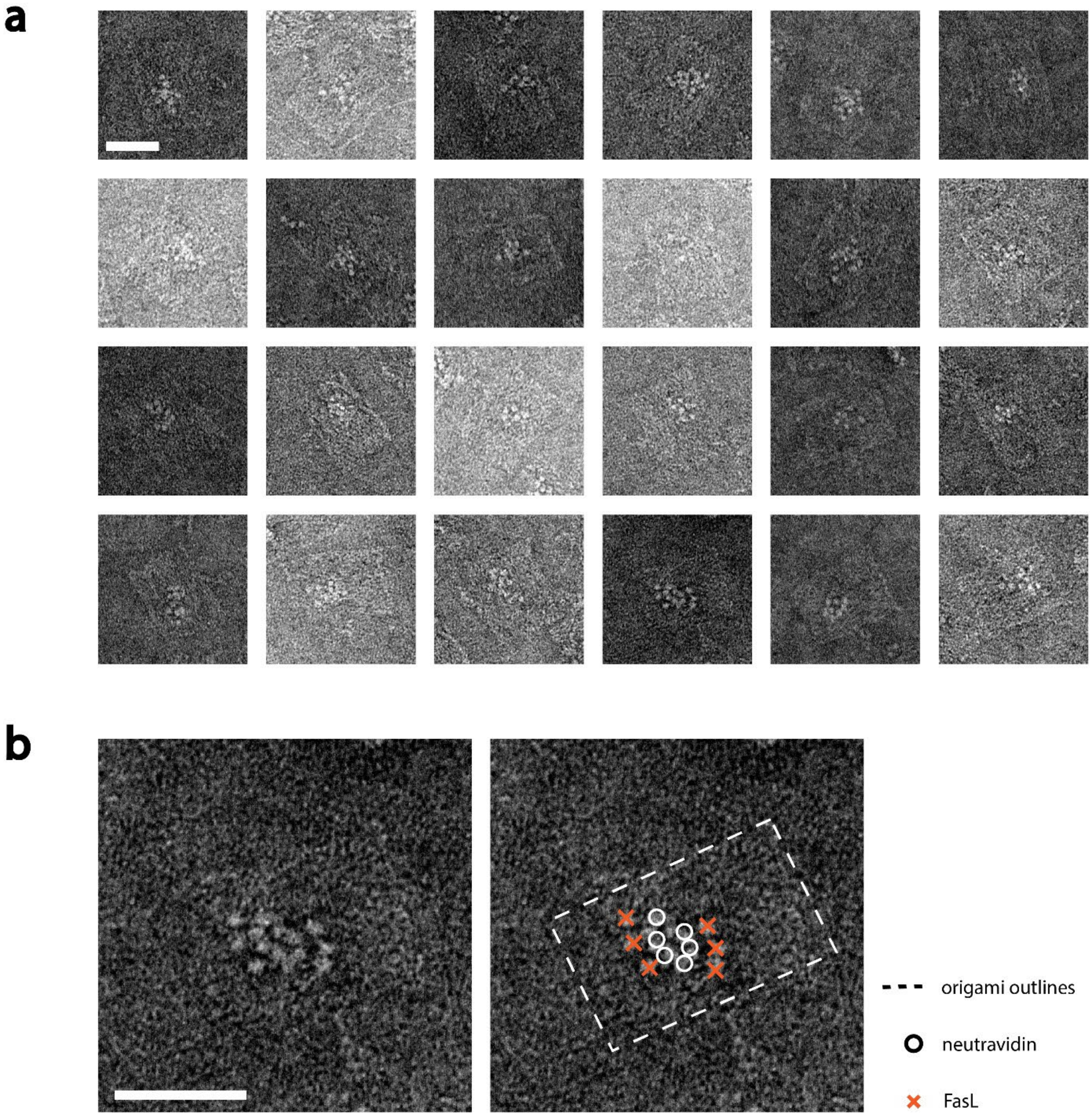

**Figure S16: TEM characterization of the rroONF nanoagents:** (**a**) Cropped micrographs show single rro DNA origami with neutravidin and FasL attached to them. Neutravidin and FasL proteins are identifiable as white spots. (**b**) Zoom-in of one rroONF nanoagent, duplicate on the right with DNA origami outlines and proteins marked. The attachment efficiency of FasL to the nanoagent was determined to be 76 % in a previous publication[2]. The scale bars are 50 nm and hold for all micrographs of the respective subfigure.

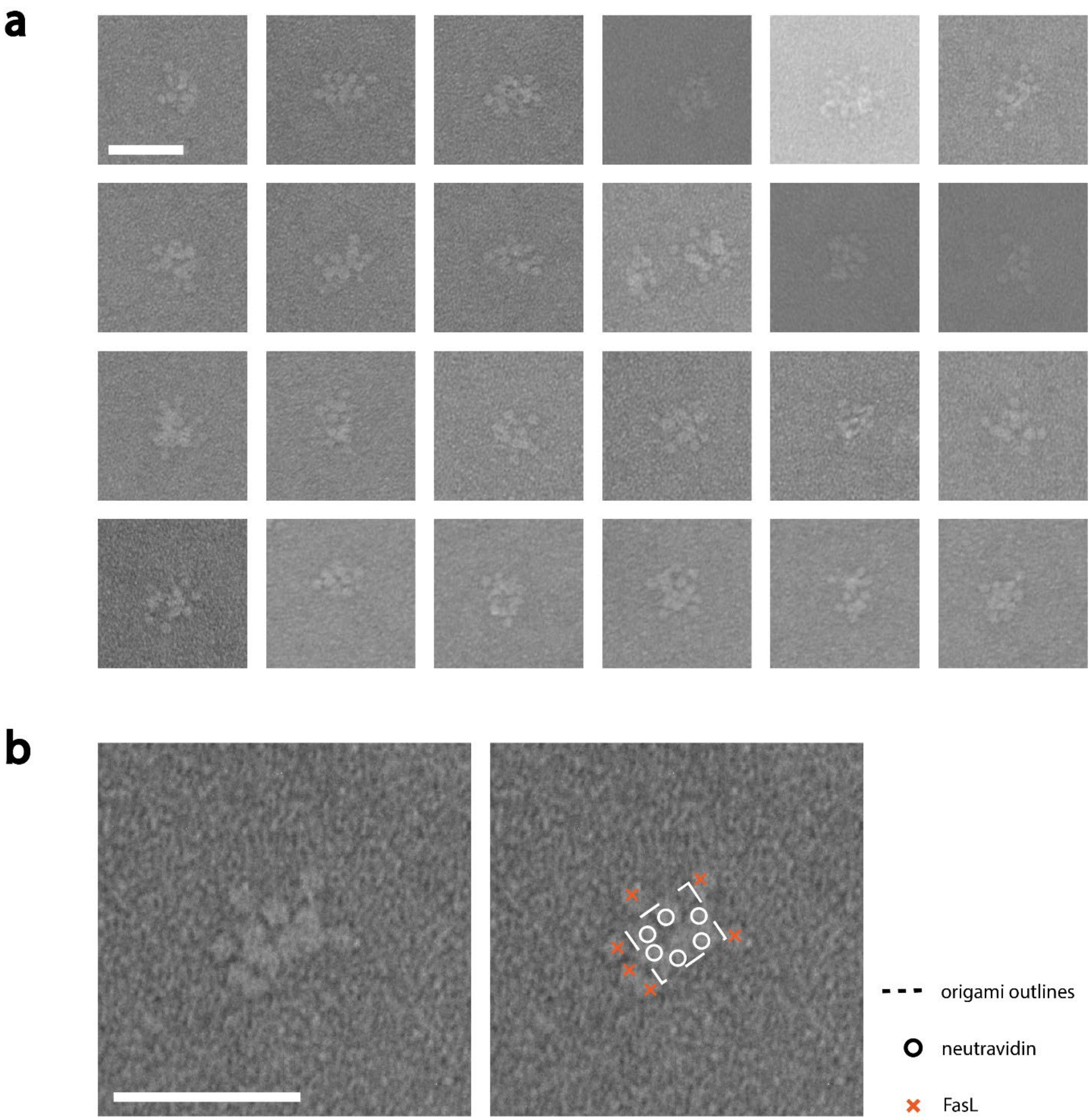

**Figure S17: TEM characterization of the miniONF nanoagents:** (**a**) Cropped micrographs show single mini DNA origami with neutravidin and FasL attached to them. Neutravidin and FasL proteins are identifiable as white spots. (**b**) Zoom-in of one miniONF nanoagent, duplicate on the right with DNA origami outlines and proteins marked. The attachment efficiency of FasL to the nanoagent was determined to be 76 % in a previous publication[2]. The scale bars are 50 nm and hold for all micrographs of the respective subfigure.

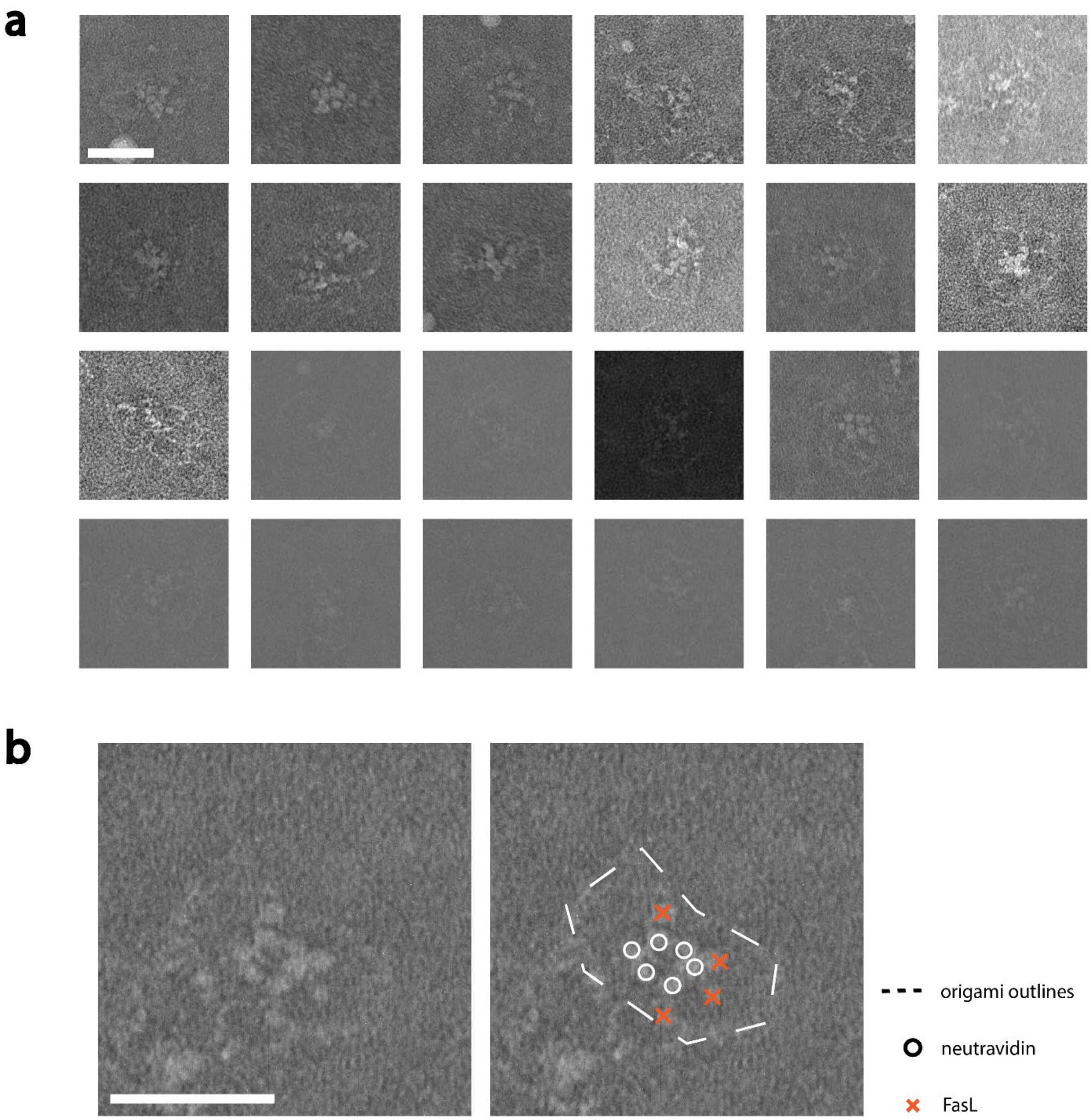

**Figure S18: TEM characterization of the wfONF nanoagents:** (**a**) Cropped micrographs show single wf DNA origami with neutravidin and FasL attached to them. Neutravidin and FasL proteins are identifiable as white spots. (**b**) Zoom-in of one wfONF nanoagent, duplicate on the right with DNA origami outlines and proteins marked. The attachment efficiency of FasL to the nanoagent was determined to be 76 % in a previous publication[2]. The scale bars are 50 nm and hold for all micrographs of the respective subfigure.

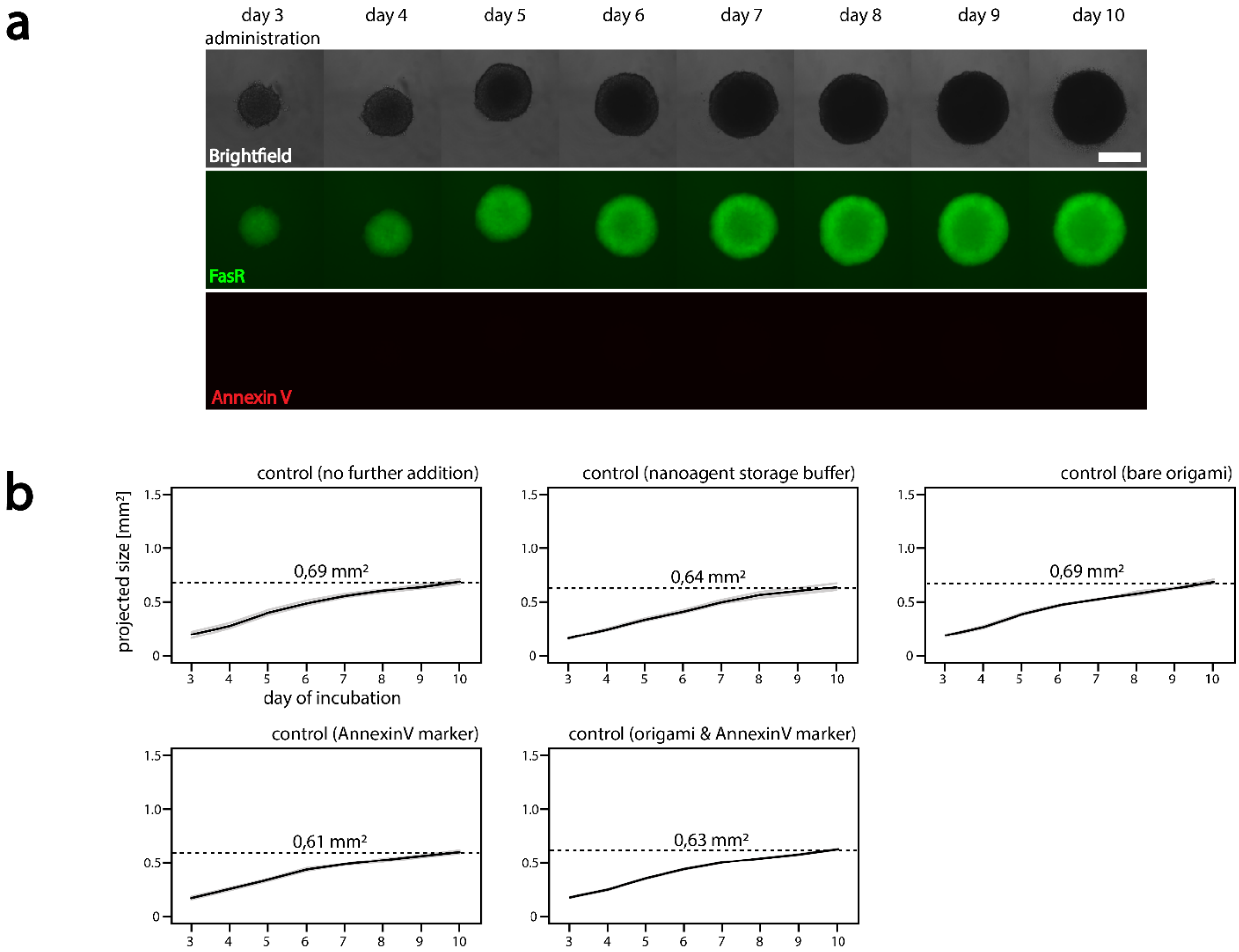

**Figure S19: development curves of control spheroids** (**a**) Brightfield, GFP (FasR), and Texas Red (Annexin V) images of a spheroid over 7 d (from day 3 to day 10). (**b**) Size projections of spheroids incubated with no additives, nanoagent storage buffer, DNA origami, AnnexinV marker, and the combination of Annexin V marker and DNA origami. Data were also used in Figure 3. The scale bar in (**a**) is 500 µm and holds for all images. Thin grey lines indicate single experiments and thick black lines indicate averages of those (n=3).

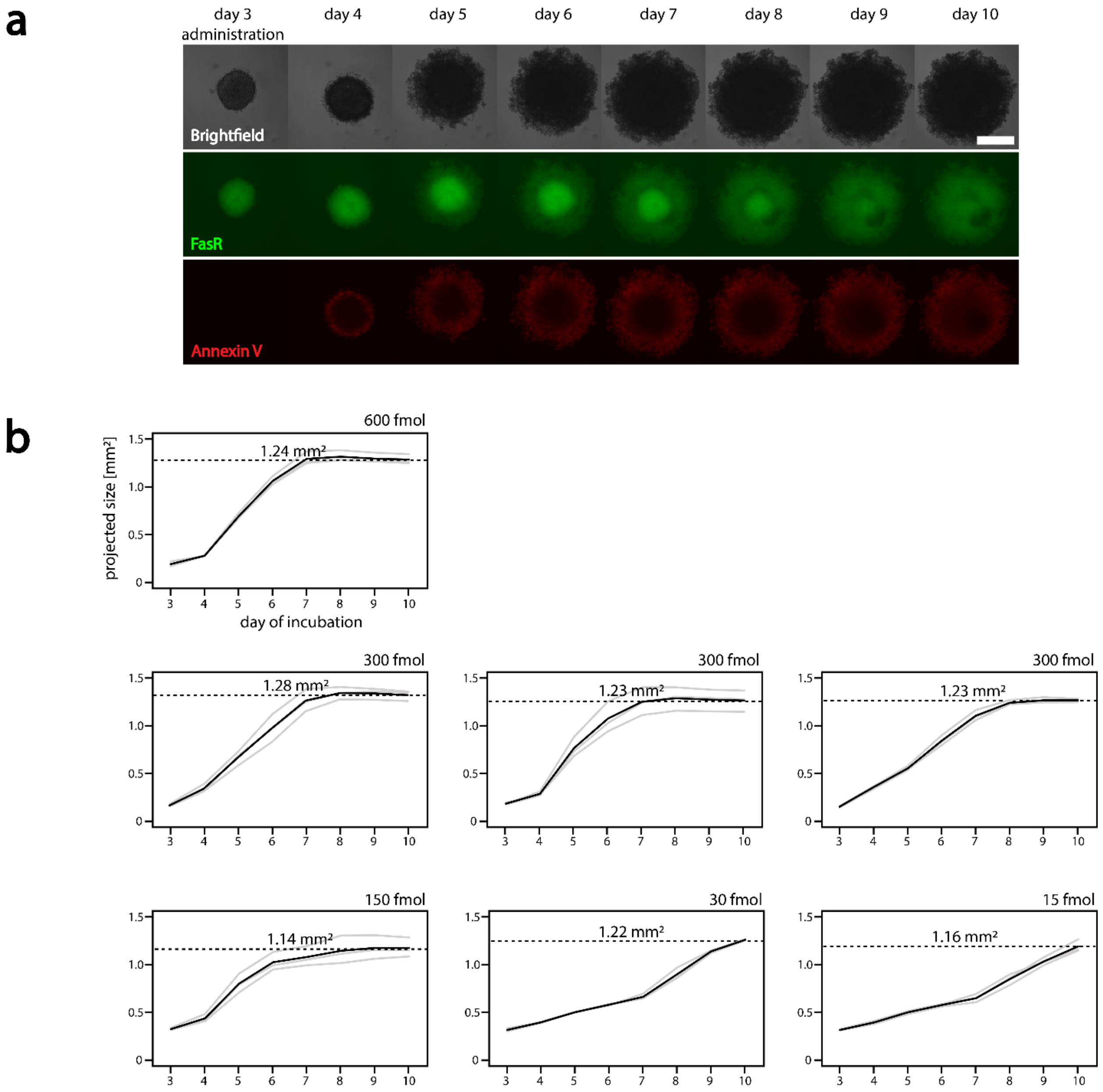

**Figure S20: development curves of spheroids with FasL** (**a**) Brightfield, GFP (FasR), and Texas Red (Annexin V) images of a spheroid incubated with 300 fmol of FasL over 7 d (from day 3 to day 10). (**b**) Size projections of spheroids incubated with 600 fmol, 300 fmol (triplicate of triplicates), 150 fmol, 30 fmol, and 15 fmol nanoagent. Data were also used in Figure 3. The scale bar in (**a**) is 500 µm and holds for all images. Thin grey lines indicate single experiments and thick black lines indicate averages of those (n=3).

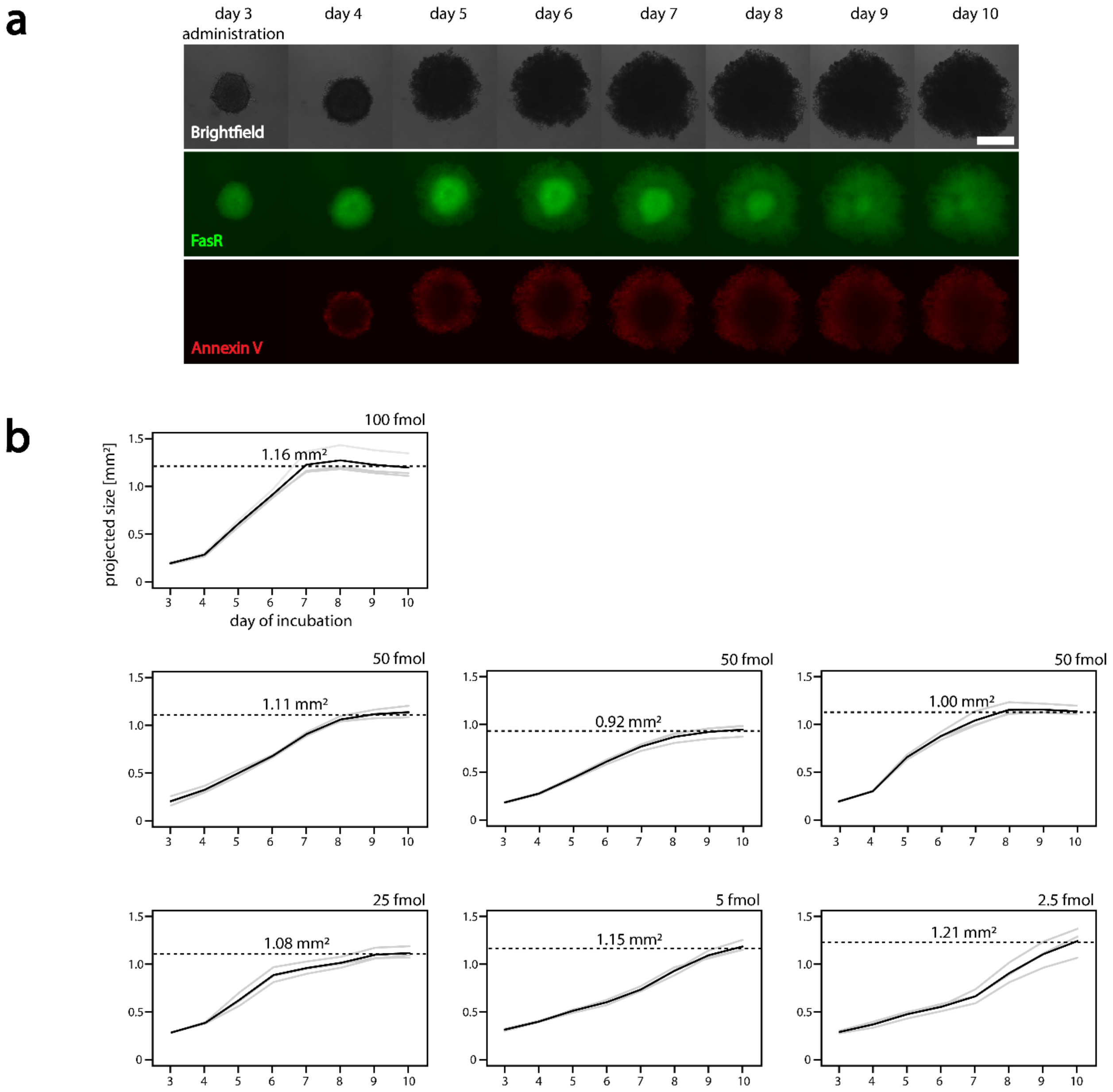

**Figure S21: development curves of spheroids with rroOF nanoagent** (**a**) Brightfield, GFP (FasR), and Texas Red (Annexin V) images of a spheroid with 50 fmol of rroOF nanoagent over 7 d (from day 3 to day 10). (**b**) Size projections of spheroids incubated with 100 fmol, 50 fmol (triplicate of triplicates), 25 fmol, 5 fmol, and 2.5 fmol nanoagent. Data were also used in Figure 3. The scale bar in (**a**) is 500 µm and holds for all images. Thin grey lines indicate single experiments and thick black lines indicate averages of those (n=3).

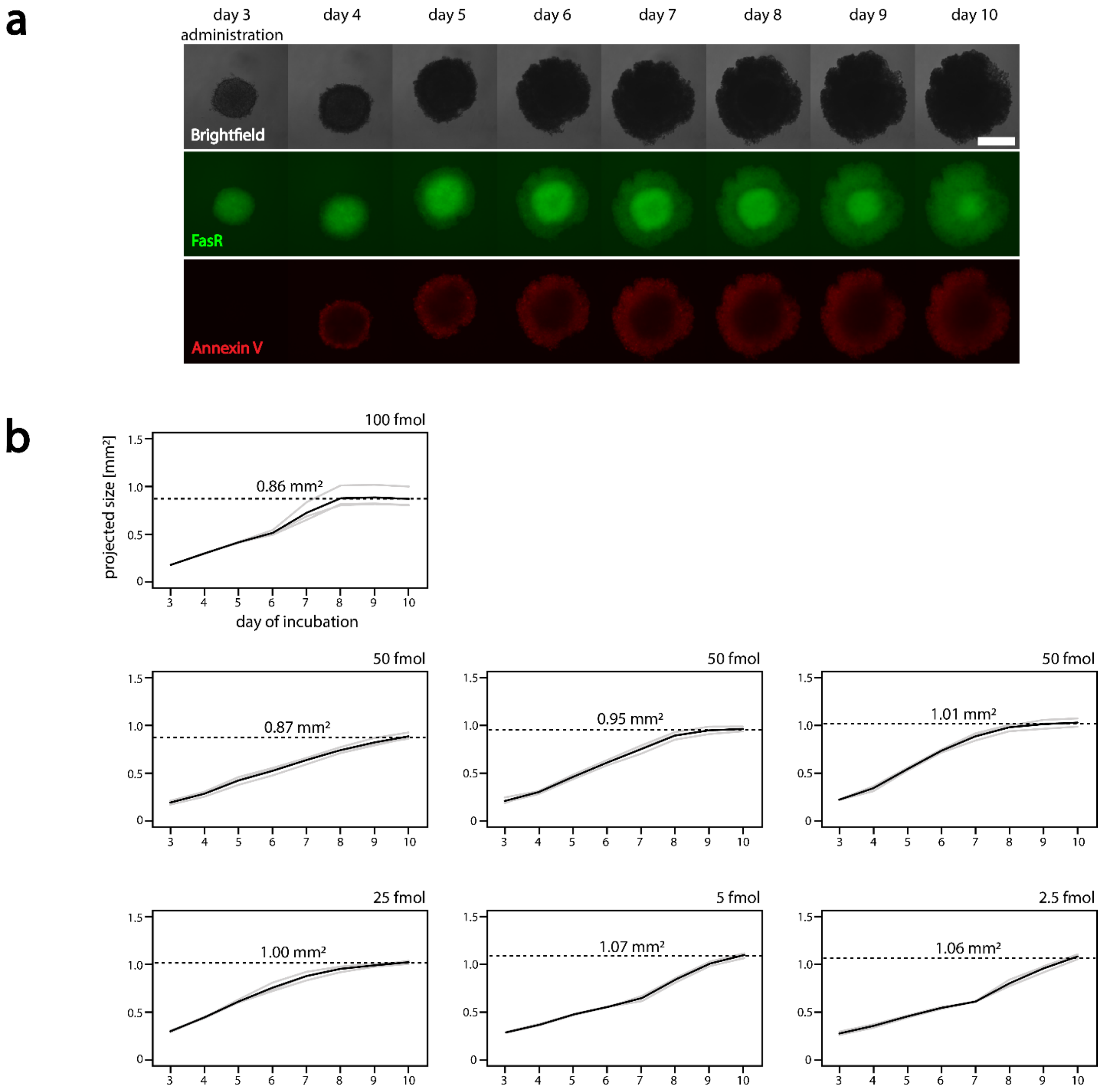

**Figure S22: development curves of spheroids with miniOF nanoagent** (**a**) Brightfield, GFP (FasR), and Texas Red (Annexin V) images of a spheroid with 50 fmol of miniOF nanoagent over 7 d (from day 3 to day 10). (**b**) Size projections of spheroids incubated with 100 fmol, 50 fmol (triplicate of triplicates), 25 fmol, 5 fmol, and 2.5 fmol nanoagent. Data were also used in Figure 3. The scale bar in (**a**) is 500 µm and holds for all images. Thin grey lines indicate single experiments and thick black lines indicate averages of those (n=3).

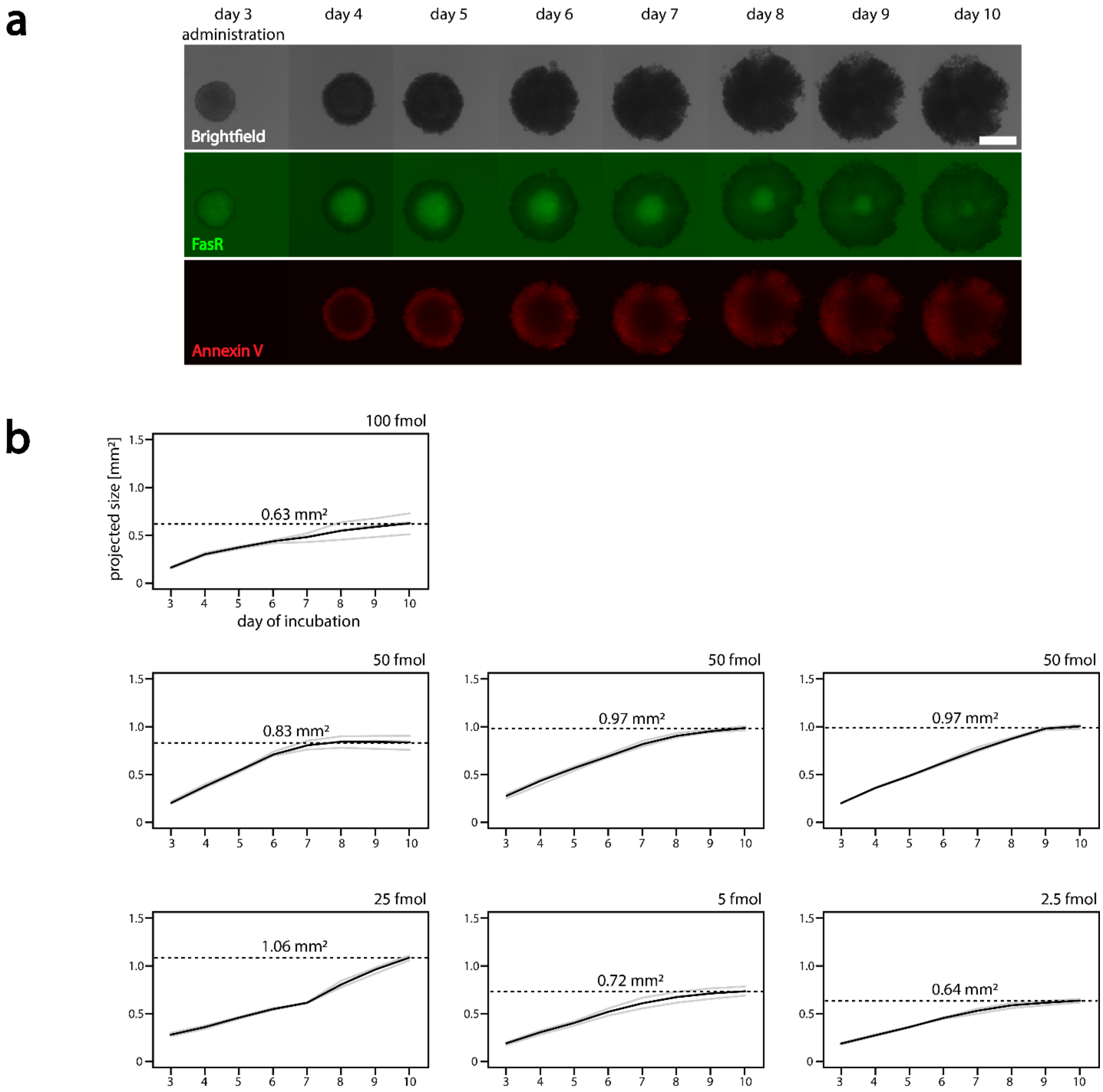

**Figure S23: development curves of spheroids with wfOF nanoagent** (**a**) Brightfield, GFP (FasR), and Texas Red (Annexin V) images of a spheroid with 50 fmol of wfOF nanoagent over 7 d (from day 3 to day 10). (**b**) Size projections of spheroids incubated with 100 fmol, 50 fmol (triplicate of triplicates), 25 fmol, 5 fmol, and 2.5 fmol nanoagent. Data were also used in Figure 3. The scale bar in (**a**) is 500 µm and holds for all images. Thin grey lines indicate single experiments and thick black lines indicate averages of those (n=3).

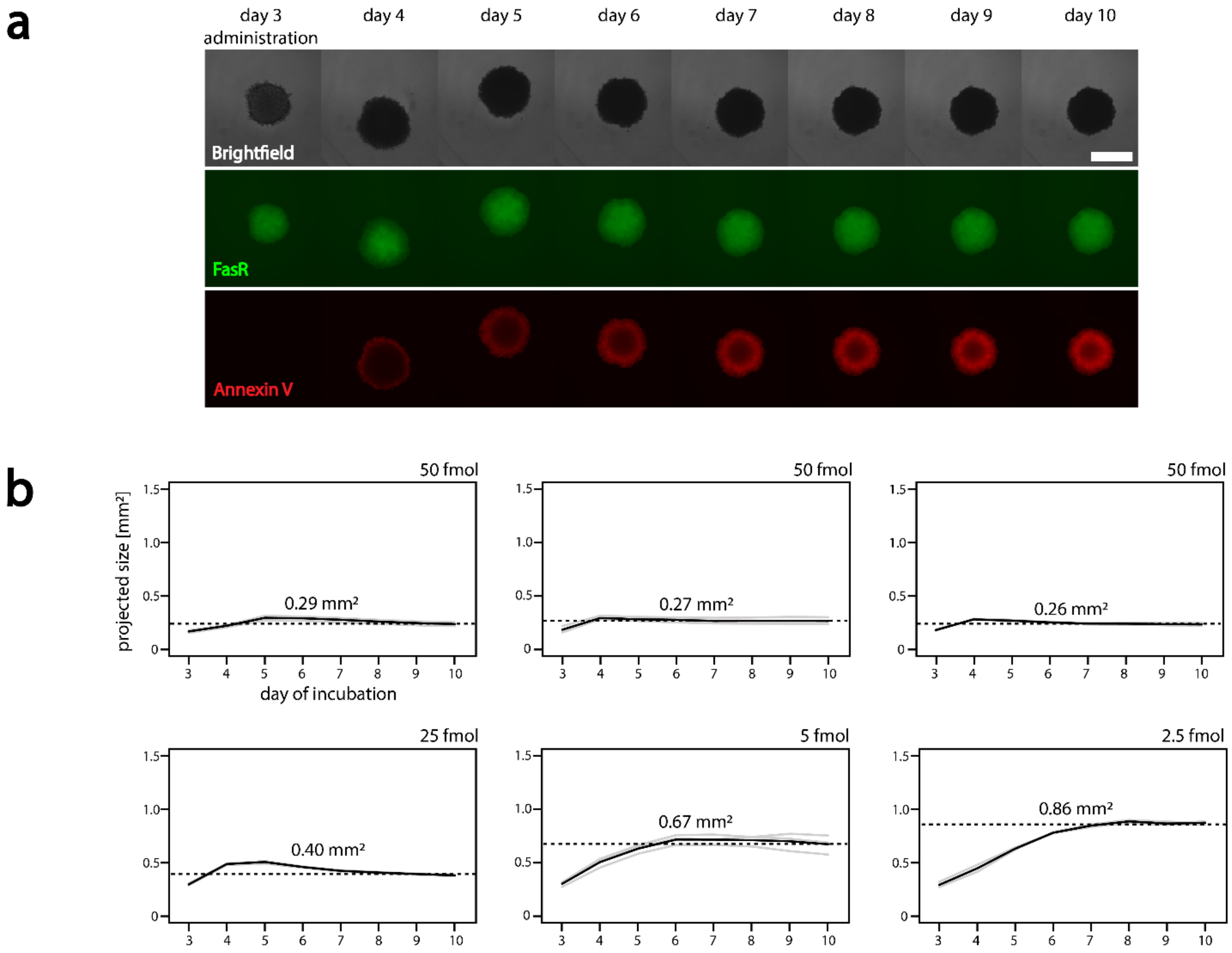

**Figure S24: development curves of spheroids with rroONF nanoagent** (**a**) Brightfield, GFP (FasR), and Texas Red (Annexin V) images of a spheroid with 50 fmol of rroONF nanoagent over 7 d (from day 3 to day 10). (**b**) Size projections of spheroids incubated with 50 fmol (triplicate of triplicates), 25 fmol, 5 fmol, and 2.5 fmol nanoagent. Data were also used in Figure 3. The scale bar in (**a**) is 500 µm and holds for all images. Thin grey lines indicate single experiments and thick black lines indicate averages of those (n=3).

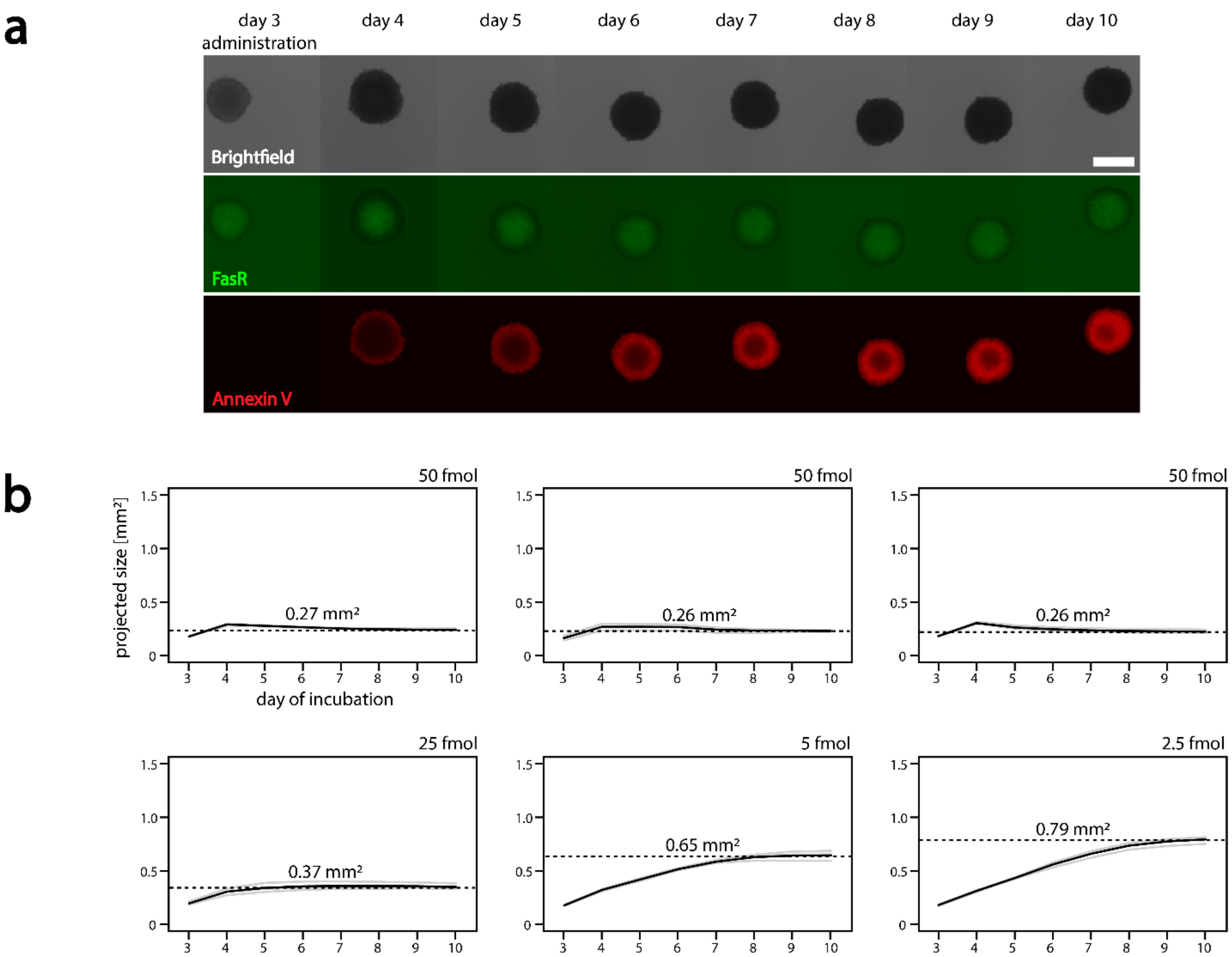

**Figure S25: development curves of spheroids with miniONF nanoagent** (**a**) Brightfield, GFP (FasR), and Texas Red (Annexin V) images of a spheroid with 50 fmol of miniONF nanoagent over 7 d (from day 3 to day 10). (**b**) Size projections of spheroids incubated with 50 fmol (triplicate of triplicates), 25 fmol, 5 fmol, and 2.5 fmol nanoagent. Data were also used in Figure 3. The scale bar in (**a**) is 500 µm and holds for all images. Thin grey lines indicate single experiments and thick black lines indicate averages of those (n=3).

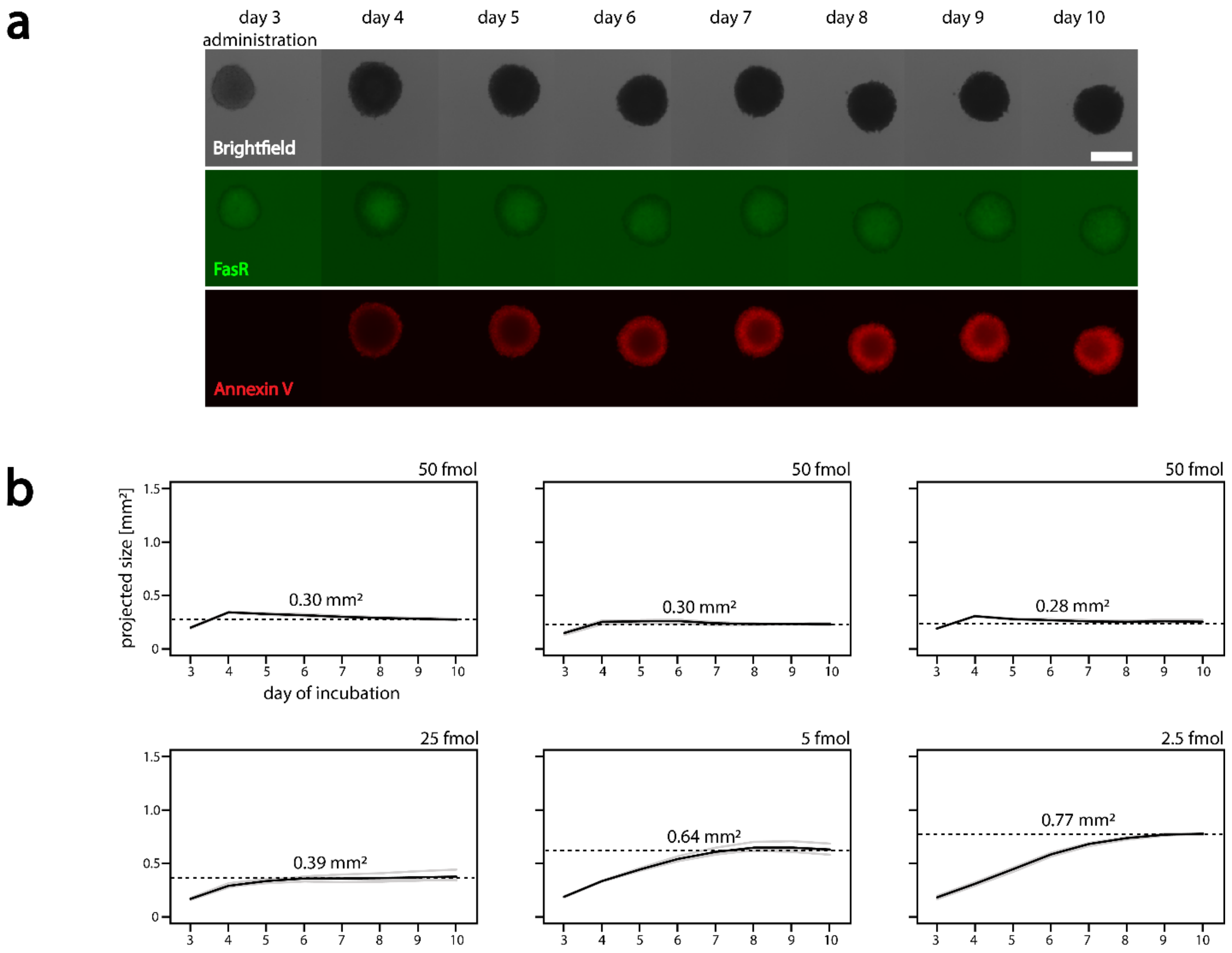

**Figure S26: development curves of spheroids with wfONF nanoagent** (**a**) Brightfield, GFP (FasR), and Texas Red (Annexin V) images of a spheroid with 50 fmol of wfONF nanoagent over 7 d (from day 3 to day 10). (**b**) Size projections of spheroids incubated with 50 fmol (triplicate of triplicates), 25 fmol, 5 fmol, and 2.5 fmol nanoagent. Data were also used in Figure 3. The Scale bar in (**a**) is 500 µm. Thin grey lines indicate single experiments and thick black lines indicate averages of those (n=3).

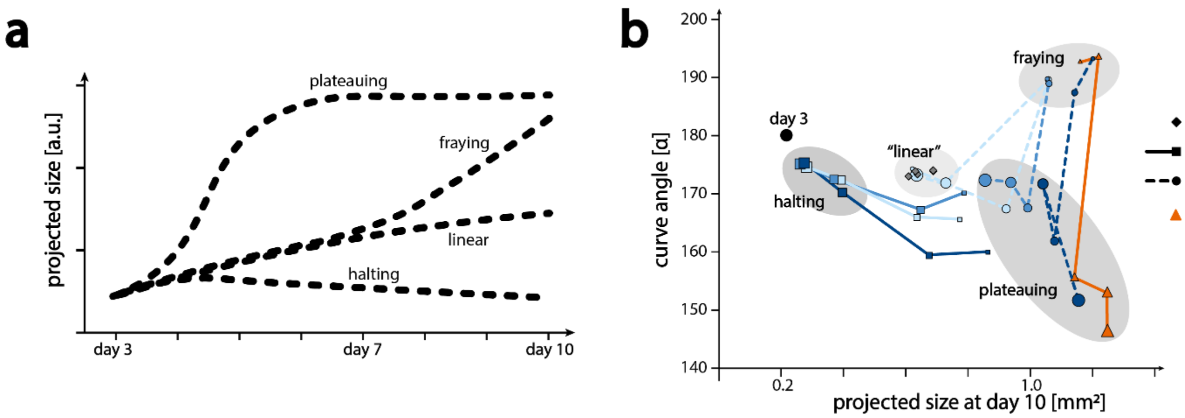

**Figure S27: pseudo-phase diagram of spheroid behavior** (**a**) Different qualitative behaviors of spheroids, plateauing was observed for large amounts of FasL or OF nanoagents, lower amounts of those led to fraying, then to linear growth, similar to the controls. Halting behavior was observed for larger amounts of ONF nanoagents. (**b**) Pseudo-phase diagram, of spheroid behavior, where the opening angle of the curves in (**a**) are plotted against their final spheroid size (averages from Figures S19-S26). Concentrations of nanoagents are indicated by the size of the respective markers. Several distinct populations are observed. Linear growth is observed for all controls. FasL and OF nanoagents induce plateauing and with lower concentrations, fraying behavior. The halting behavior, induced by ONF nanoagents, is distinctly different from the others.

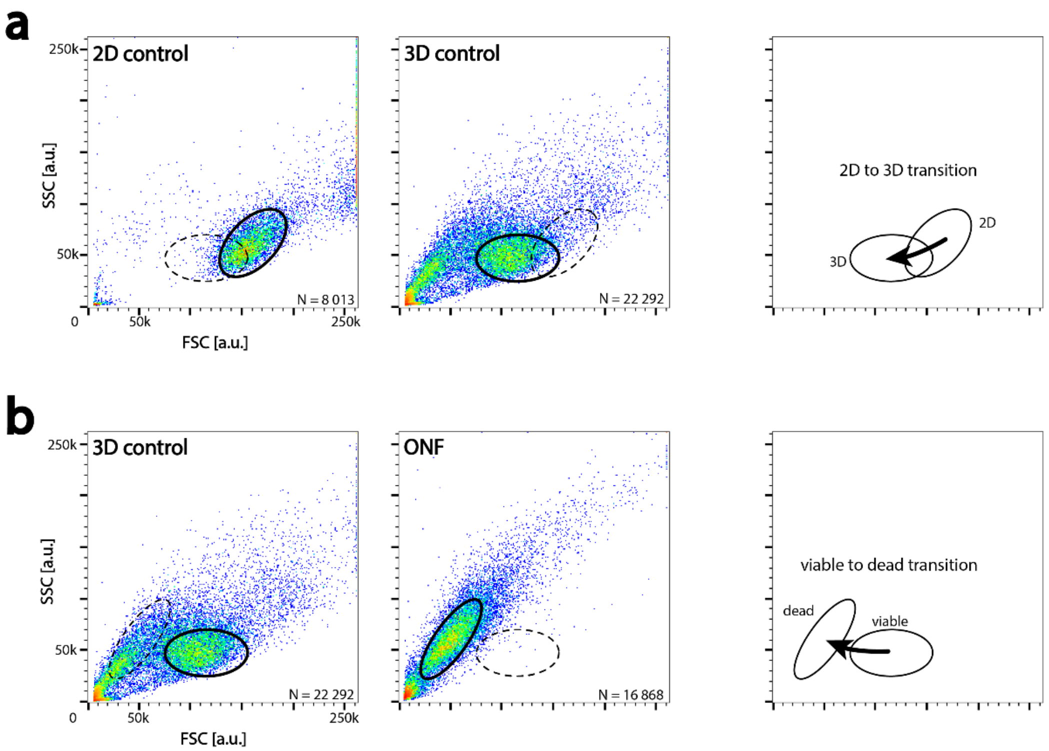

**Figure S28: population shifts in fluorescence activated cell sorting (FACS)** are seen when (**a**) comparing cells cultured in 2D vs. 3D : the 3D control shows a shift in FSC, indicating a size decrease of individual cells cultured in 3D. (**b**) The effectiveness of the respective nanoagent was measured as the ratio between events in viable and dead gates. The higher the ratio of cells in the dead gate compared to the viable gate, the more effective the nanoagent.

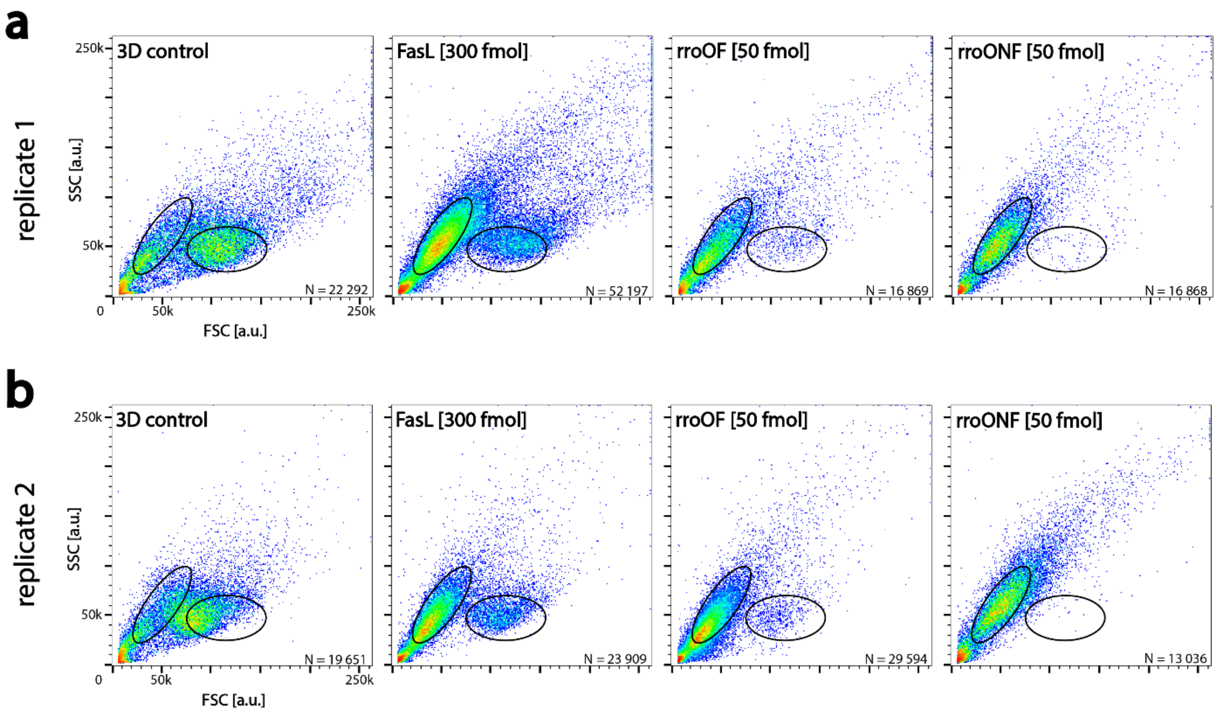

**Figure S29: FACS data of dissolved spheroids** in two replicates (a) and (b). Incubated with 300 fmol FasL, 50 fmol of rroOF or rroONF nanoagent or without addition, as controls. Both replicates, (**a**) and (**b**), show similar distributions depending on which nanoagent was added: The ratio of events in the viable gates to dead gates is largest in the control, and decreases when the spheroids were incubated with FasL. When OF nanoagent was added, the ratio was again lower, and with rroONF nanoagent, nearly no events remained in the viable gate. Data from the same experiment was used for Figure 4.

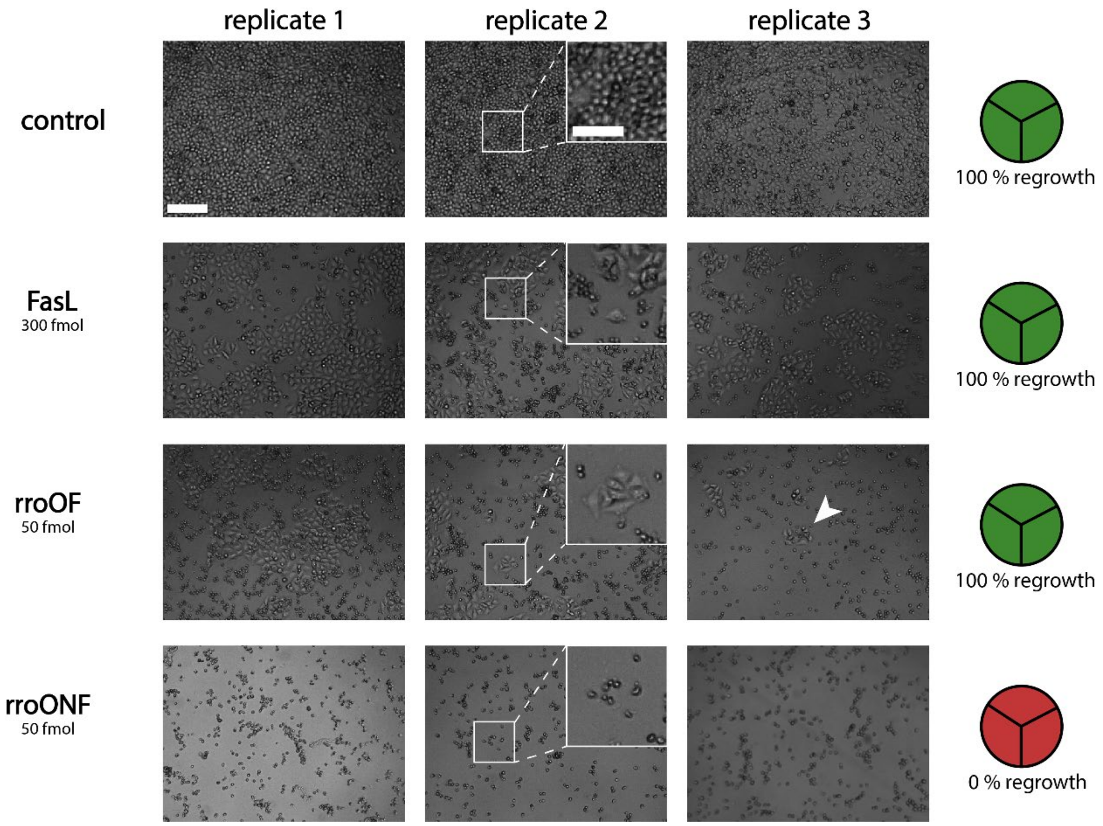

**Figure S30: 2D regrowth of dissolved spheroids** after 10 d of incubation with the respective additive, dissociation, and seeding on a cell culture-treated dish. Images were taken after 2 d of incubation, and badly visible colonies were marked with a white arrow. rroONF data is the same as in Figure S28. Data was also used in Figure 4. The scale bar is 200 µm and holds for every overview image, and the scale bar in the zoom-in is 100 µm.

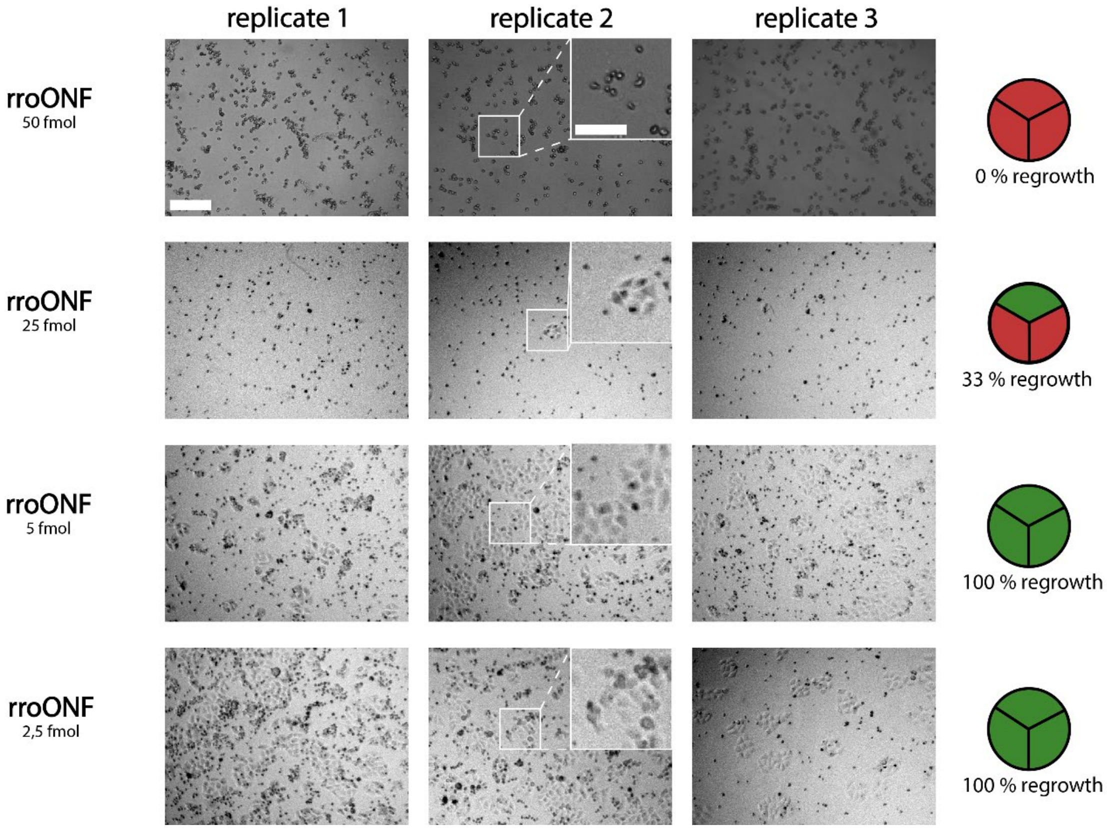

**Figure S31: 2D regrowth of dissolved spheroids, titrating concentrations:** After 10 d of incubation with the respective nanoagent, dissociation, and seeding on a cell culture-treated dish. Images were taken after 2 d of incubation. rroONF data is the same as in Figure S27, data was also used in Figure 4. The scale bar is 200 µm and holds for every overview image, and the scale bar in the zoom-in is 100 µm.

### Table S1: rro DNA origami staples

| name | sequence |
|---|---|
| rro_core_001 | TTTTCACTCAAAGGGCGAAAAACCATCACC |
| rro_core_002 | GTCGACTTCGGCCAACGCGCGGGGTTTTTC |
| rro_core_003 | TGCATCTTTCCCAGTCACGACGGCCTGCAG |
| rro_core_004 | TAATCAGCGGATTGACCGTAATCGTAACCG |
| rro_core_005 | AACGCAAAATCGATGAACGGTACCGGTTGA |
| rro_core_006 | AACAGTTTTGTACCAAAAACATTTTATTTC |
| rro_core_007 | TTTACCCCAACATGTTTTAAATTTCCATAT |
| rro_core_008 | TTTAGGACAAATGCTTTAAACAATCAGGTC |
| rro_core_009 | CATCAAGTAAAACGAACTAACGAGTTGAGA |
| rro_core_010 | AATACGTTTGAAAGAGGACAGACTGACCTT |
| rro_core_011 | AGGCTCCAGAGGCTTTGAGGACACGGGTAA |
| rro_core_012 | AGAAAGGAACAACTAAAGGAATTCAAAAAAA |
| rro_core_013 | CAAATCAAGTTTTTTGGGGTCGAAACGTGGA |
| rro_core_014 | CTCCAACGCAGTGAGACGGGCAACCAGCTGCA |
| rro_core_015 | TTAATGAACTAGAGGATCCCCGGGGGGTAACG |
| rro_core_016 | CCAGGGTTGCCAGTTTGAGGGGACCCGTGGGA |
| rro_core_017 | ACAAACGGAAAAGCCCCAAAAACACTGGAGCA |
| rro_core_018 | AACAAGAGGGATAAAAATTTTTAGCATAAAGC |
| rro_core_019 | TAAATCGGGATTCCCAATTCTGCGATATAATG |
| rro_core_020 | CTGTAGCTTGACTATTATAGTCAGTTCATTGA |
| rro_core_021 | ATCCCCCTATACCACATTCAACTAGAAAAATC |
| rro_core_022 | TACGTTAAAGTAATCTTGACAAGAACCGAACT |
| rro_core_023 | GACCAACTAATGCCACTACGAAGGGGGTAGCA |
| rro_core_024 | ACGGCTACAAAAGGAGCCTTTAATGTGAGAAT |
| rro_core_025 | AGCTGATTGCCCTTCAGAGTCCACTATTAAAGGGTGCCGT |
| rro_core_026 | GTATAAGCCAACCCGTCGGATTCTGACGACAGTATCGGCCGCAAGGCG |
| rro_core_027 | TATATTTTGTCATTGCCTGAGAGTGGAAGATT |
| rro_core_028 | GATTTAGTCAATAAAGCCTCAGAGAACCCTCA |
| rro_core_029 | CGGATTGCAGAGCTTAATTGCTGAAACGAGTA |
| rro_core_030 | ATGCAGATACATAACGGGAATCGTCATAAATAAAGCAAAG |
| rro_core_031 | TTTATCAGGACAGCATCGGAACGACACCAACCTAAAACGAGGTCAATC |
| rro_core_032 | ACAACTTTCAACAGTTTCAGCGGATGTATCGG |
| rro_core_033 | AAAGCACTAAATCGGAACCCTAATCCAGTT |
| rro_core_034 | TGGAACAACCGCCTGGCCCTGAGGCCCGCT |
| rro_core_035 | TTCCAGTCGTAATCATGGTCATAAAAGGGG |
| rro_core_036 | GATGTGCTTCAGGAAGATCGCACAATGTGA |
| rro_core_037 | GCGAGTAAAAATATTTAAATTGTTACAAAG |
| rro_core_038 | GCTATCAGAAATGCAATGCCTGAATTAGCA |
| rro_core_039 | AAATTAAGTTGACCATTAGATACTTTTGCG |
| rro_core_040 | GATGGCTTATCAAAAAGATTAAGAGCGTCC |

| | |
|---|---|
| rro_core_041 | AATACTGCCCAAAAGGAATTACGTGGCTCA |
| rro_core_042 | TTATACCACCAAATCAACGTAACGAACGAG |
| rro_core_043 | GCGCAGACAAGAGGCAAAAGAATCCCTCAG |
| rro_core_044 | CAGCGAAACTTGCTTTCGAGGTGTTGCTAA |
| rro_core_045 | AGCAAGCGTAGGGTTGAGTGTTGTAGGGAGCC |
| rro_core_046 | CTGTGTGATTGCGTTGCGCTCACTAGAGTTGC |
| rro_core_047 | GCTTTCCGATTACGCCAGCTGGCGGCTGTTTC |
| rro_core_048 | ATATTTTGGCTTTCATCAACATTATCCAGCCA |
| rro_core_049 | TAGGTAAACTATTTTTGAGAGATCAAACGTTA |
| rro_core_050 | AATGGTCAACAGGCAAGGCAAAGAGTAATGTG |
| rro_core_051 | TAAGAGCAAATGTTTAGACTGGATAGGAAGCC |
| rro_core_052 | TCATTCAGATGCGATTTTAAGAACAGGCATAG |
| rro_core_053 | ACACTCATCCATGTTACTTAGCCGAAAGCTGC |
| rro_core_054 | AAACAGCTTTTTGCGGGATCGTCAACACTAAA |
| rro_core_055 | TAAATGAATTTTCTGTATGGGATTAATTTCTT |
| rro_core_056 | CCCGATTTAGAGCTTGACGGGGAAAAAGAATA |
| rro_core_057 | GCCCGAGAGTCCACGCTGGTTTGCAGCTAACT |
| rro_core_058 | CACATTAAAATTGTTATCCGCTCATGCGGGCC |
| rro_core_059 | TCTTCGCTGCACCGCTTCTGGTGCGGCCTTCC |
| rro_core_060 | GAGGGTAGGATTCAAAAGGGTGAGACATCCAA |
| rro_core_061 | TAAATCATATAACCTGTTTAGCTAACCTTTAA |
| rro_core_062 | AATAGTAAACACTATCATAACCCTCATTGTGA |
| rro_core_063 | ATTACCTTTGAATAAGGCTTGCCCAAATCCGC |
| rro_core_064 | GACCTGCTCTTTGACCCCCAGCGAGGGAGTTA |
| rro_core_065 | AAGGCCGCTGATACCGATAGTTGCGACGTTAG |
| rro_core_066 | CCCAGCAGGCGAAAAATCCCTTATAAATCAAGCCGGCG |
| rro_core_067 | TAAATCAAAATAATTCGCGTCTCGGAAACCAGGCAAAGGGAAGG |
| rro_core_068 | GAGACAGCTAGCTGATAAATTAATTTTTGT |
| rro_core_069 | TTTGGGGATAGTAGTAGCATTAAAAGGCCG |
| rro_core_070 | GCTTCAATCAGGATTAGAGAGTTATTTTCA |
| rro_core_071 | CGTTTACCAGACGACAAAGAAGTTTTGCCATAATTCGA |
| rro_core_072 | TGACAACTCGCTGAGGCTTGCATTATACCAAGCGCGATGATAAA |
| rro_core_073 | TCTAAAGTTTTGTCGTCTTTCCAGCCGACAA |
| rro_core_074 | TCAATATCGAACCTCAAATATCAATTCCGAAA |
| rro_core_075 | GCAATTCACATATTCCTGATTATCAAAGTGTA |
| rro_core_076 | AGAAAACAAAGAAGATGATGAAACAGGCTGCG |
| rro_core_077 | ATCGCAAGTATGTAAATGCTGATGATAGGAAC |
| rro_core_078 | CCAATAGCTCATCGTAGGAATCATGGCATCAA |
| rro_core_079 | AGAGAGAAAAAAATGAAAATAGCAAGCAAACT |
| rro_core_080 | GCAAGGCCTCACCAGTAGCACCATGGGCTTGA |
| rro_core_081 | TTGACAGGCCACCACCAGAGCCGCGATTTGTA |
| rro_core_082 | TTAGGATTGGCTGAGACTCCTCAATAACCGAT |
| rro_core_083 | TCCACAGACAGCCCTCATAGTTAGCGTAACGA |

| | |
|---|---|
| rro_core_084 | AACGTGGCGAGAAAGGAAGGGAAACCAGTAA |
| rro_core_085 | TCGGCAAATCCTGTTTGATGGTGGACCCTCAA |
| rro_core_086 | AAGCCTGGTACGAGCCGGAAGCATAGATGATG |
| rro_core_087 | CAACTGTTGCGCCATTCGCCATTCAAACATCA |
| rro_core_088 | GCCATCAAGCTCATTTTTTAACCACAAATCCA |
| rro_core_089 | CAACCGTTTCAAATCACCATCAATTCGAGCCA |
| rro_core_090 | CCAACAGGAGCGAACCAGACCGGAGCCTTTAC |
| rro_core_091 | CTTTTGCAGATAAAAACCAAAATAAAGACTCC |
| rro_core_092 | GATGGTTTGAACGAGTAGTAAATTTACCATTA |
| rro_core_093 | TCATCGCCAACAAAGTACAACGGACGCCAGCA |
| rro_core_094 | ATATTCGGAACCATCGCCCACGCAGAGAAGGA |
| rro_core_095 | TAAAAGGGACATTCTGGCCAACAAAGCATC |
| rro_core_096 | ACCTTGCTTGGTCAGTTGGCAAAGAGCGGA |
| rro_core_097 | ATTATCATTCAATATAATCCTGACAATTAC |
| rro_core_098 | CTGAGCAAAAATTAATTACATTTTGGGTTA |
| rro_core_099 | TATAACTAACAAAGAACGCGAGAACGCCAA |
| rro_core_100 | CATGTAATAGAATATAAAGTACCAAGCCGT |
| rro_core_101 | TTTTATTTAAGCAAATCAGATATTTTTTGT |
| rro_core_102 | TTAACGTCTAACATAAAAACAGGTAACGGA |
| rro_core_103 | ATACCCAACAGTATGTTAGCAAATTAGAGC |
| rro_core_104 | CAGCAAAAGGAAACGTCACCAATGAGCCGC |
| rro_core_105 | CACCAGAAAGGTTGAGGCAGGTCATGAAAG |
| rro_core_106 | TATTAAGAAGCGGGGTTTTGCTCGTAGCAT |
| rro_core_107 | TCAACAGTTGAAAGGAGCAAATGAAAAATCTAGAGATAGA |
| rro_core_108 | TCAAATATAACCTCCGGCTTAGGTAACAATTTCATTTGAAGGCGAATT |
| rro_core_109 | GTAAAGTAATCGCCATATTTAACAAAACTTTT |
| rro_core_110 | TATCCGGTCTCATCGAGAACAAGCGACAAAAG |
| rro_core_111 | TTAGACGGCCAAATAAGAAACGATAGAAGGCT |
| rro_core_112 | CGTAGAAAATACATACCGAGGAAACGCAATAAGAAGCGCA |
| rro_core_113 | GCGGATAACCTATTATTCTGAAACAGACGATTGGCCTTGAAGAGCCAC |
| rro_core_114 | TCACCAGTACAAACTACAACGCCTAGTACCAG |
| rro_core_115 | ACCCTTCTGACCTGAAAGCGTAAGACGCTGAG |
| rro_core_116 | AGCCAGCAATTGAGGAAGGTTATCATCATTTT |
| rro_core_117 | GCGGAACATCTGAATAATGGAAGGTACAAAAT |
| rro_core_118 | CGCGCAGATTACCTTTTTTAATGGGAGAGACT |
| rro_core_119 | ACCTTTTTATTTTAGTTAATTTCATAGGGCTT |
| rro_core_120 | AATTGAGAATTCTGTCCAGACGACTAAACCAA |
| rro_core_121 | GTACCGCAATTCTAAGAACGCGAGTATTATTT |
| rro_core_122 | ATCCCAATGAGAATTAACTGAACAGTTACCAG |
| rro_core_123 | AAGGAAACATAAAGGTGGCAACATTATCACCG |
| rro_core_124 | TCACCGACGCACCGTAATCAGTAGCAGAACCG |
| rro_core_125 | CCACCCTCTATTCACAAACAAATACCTGCCTA |
| rro_core_126 | TTTCGGAAGTGCCGTCGAGAGGGTGAGTTTCG |

| | |
|---|---|
| rro_core_127 | CTTTAGGGCCTGCAACAGTGCCAATACGTG |
| rro_core_128 | CTACCATAGTTTGAGTAACATTTAAAATAT |
| rro_core_129 | CATAAATCTTTGAATACCAAGTGTTAGAAC |
| rro_core_130 | CCTAAATCAAAATCATAGGTCTAAACAGTA |
| rro_core_131 | ACAACATGCCAACGCTCAACAGTCTTCTGA |
| rro_core_132 | GCGAACCTCCAAGAACGGGTATGACAATAA |
| rro_core_133 | AAAGTCACAAAATAAACAGCCAGCGTTTTA |
| rro_core_134 | AACGCAAAGATAGCCGAACAAACCCTGAAC |
| rro_core_135 | TCAAGTTTCATTAAAGGTGAATATAAAAGA |
| rro_core_136 | TTAAAGCCAGAGCCGCCACCCTCGACAGAA |
| rro_core_137 | GTATAGCAAACAGTTAATGCCCAATCCTCA |
| rro_core_138 | AGGAACCCATGTACCGTAACACTTGATATAA |
| rro_core_139 | GCACAGACAATATTTTTGAATGGGGTCAGTA |
| rro_core_140 | TTAACACCAGCACTAACAACTAATCGTTATTA |
| rro_core_141 | ATTTTAAAATCAAAATTATTTGCACGGATTCG |
| rro_core_142 | CCTGATTGCAATATATGTGAGTGATCAATAGT |
| rro_core_143 | GAATTTATTTAATGGTTTGAAATATTCTTACC |
| rro_core_144 | AGTATAAAGTTCAGCTAATGCAGATGTCTTTC |
| rro_core_145 | CTTATCATTCCCGACTTGCGGGAGCCTAATTT |
| rro_core_146 | GCCAGTTAGAGGGTAATTGAGCGCTTTAAGAA |
| rro_core_147 | AAGTAAGCAGACACCACGGAATAATATTGACG |
| rro_core_148 | GAAATTATTGCCTTTAGCGTCAGACCGGAACC |
| rro_core_149 | GCCTCCCTCAGAATGGAAAGCGCAGTAACAGT |
| rro_core_150 | GCCCGTATCCGGAATAGGTGTATCAGCCCAAT |
| rro_core_151 | AGATTAGAGCCGTCAAAAAACAGAGGTGAGGCCTATTAGT |
| rro_core_152 | GTGATAAAAAGACGCTGAGAAGAGATAACCTTGCTTCTGTTCGGGAGA |
| rro_core_153 | GTTTATCAATATGCGTTATACAAACCGACCGT |
| rro_core_154 | GCCTTAAACCAATCAATAATCGGCACGCGCCT |
| rro_core_155 | GAGAGATAGAGCGTCTTTCCAGAGGTTTTGAA |
| rro_core_156 | GTTTATTTTGTCACAATCTTACCGAAGCCCTTTAATATCA |
| rro_core_157 | CAGGAGGTGGGGTCAGTGCCTTGAGTCTCTGAATTTACCGGGAACCAG |
| rro_core_158 | CCACCCTCATTTTCAGGGATAGCAACCGTACT |
| rro_core_159 | CTTTAATGCGCGAACTGATAGCCCCACCAG |
| rro_core_160 | CAGAAGATTAGATAATACATTTGTCGACAA |
| rro_core_161 | CTCGTATTAGAAATTGCGTAGATACAGTAC |
| rro_core_162 | CTTTTACAAAATCGTCGCTATTAGCGATAG |
| rro_core_163 | CTTAGATTTAAGGCGTTAAATAAAGCCTGT |
| rro_core_164 | TTAGTATCACAATAGATAAGTCCACGAGCA |
| rro_core_165 | TGTAGAAATCAAGATTAGTTGCTCTTACCA |
| rro_core_166 | ACGCTAACACCCACAAGAATTGAAAATAGC |
| rro_core_167 | AATAGCTATCAATAGAAAATTCAACATTCA |
| rro_core_168 | ACCGATTGTCGGCATTTTCGGTCATAATCA |
| rro_core_169 | AAATCACCTTCCAGTAAGCGTCAGTAATAA |

| | |
|---|---|
| rro_core_170 | GTTTTAACTTAGTACCGCCACCCAGAGCCA |
| rro_anchor_01 | CATTCTCCTATTACTACCTTGTGTCGTGACGAGAAACACCAAATTTCAACTTTAAT |
| rro_anchor_02 | CATTCTCCTATTACTACCGCGATCGGCAATTCCACACAACAGGTGCCTAATGAGTG |
| rro_anchor_03 | CATTCTCCTATTACTACCCACCCTCAGAAACCATCGATAGCATTGAGCCATTTGGGAA |
| rro_anchor_04 | CATTCTCCTATTACTACCAACAATAACGTAAAACAGAAATAAAAATCCTTTGCCCGAA |
| rro_anchor_05 | CATTCTCCTATTACTACCATTAAGTTTACCGAGCTCGAATTCGGGAAACCTGTCGTGC |
| rro_anchor_06 | CATTCTCCTATTACTACCCACCCTCAGAAACCATCGATAGCATTGAGCCATTTGGGAA |
| rro_anchor_07 | CATTCTCCTATTACTACCATAAGGGAACCGGATATTCATTACGTCAGGACGTTGGGAA |
| rro_anchor_08 | CATTCTCCTATTACTACCAGCCACCACTGTAGCGCGTTTTCAAGGGAGGGAAGGTAAA |
| rro_FasL_handle_01 | CGAAAGACTTTGATAAGAGGTCATATTTCGCA TT TTCATTCTCCTATTACTACC |
| rro_FasL_handle_02 | TGTAGCCATTAAAATTCGCATTAAATGCCGGA TT TTCATTCTCCTATTACTACC |
| rro_FasL_handle_03 | TTGCTCCTTTCAAATATCGCGTTTGAGGGGGT TT TTCATTCTCCTATTACTACC |
| rro_FasL_handle_04 | GTAATAAGTTAGGCAGAGGCATTTATGATATT TT TTCATTCTCCTATTACTACC |
| rro_FasL_handle_05 | TTATTACGAAGAACTGGCATGATTGCGAGAGG TT TTCATTCTCCTATTACTACC |
| rro_FasL_handle_06 | TTCTACTACGCGAGCTGAAAAGGTTACCGCGC TT TTCATTCTCCTATTACTACC |
| rro_biotin_handle_01 | CGAAAGACTTTGATAAGAGGTCATATTTCGCA TT [biotin] |
| rro_biotin_handle_02 | TGTAGCCATTAAAATTCGCATTAAATGCCGGA TT [biotin] |
| rro_biotin_handle_03 | TTGCTCCTTTCAAATATCGCGTTTGAGGGGGT TT [biotin] |
| rro_biotin_handle_04 | GTAATAAGTTAGGCAGAGGCATTTATGATATT TT [biotin] |
| rro_biotin_handle_05 | TTATTACGAAGAACTGGCATGATTGCGAGAGG TT [biotin] |
| rro_biotin_handle_06 | TTCTACTACGCGAGCTGAAAAGGTTACCGCGC TT [biotin] |
| rro_FISH_handle_01 | CGAAAGACTTTGATAAGAGGTCATATTTCGCA GCATTCTTTCTTGAGGAGGGCAGCAAACGGGAAGAG |
| rro_FISH_handle_02 | TGTAGCCATTAAAATTCGCATTAAATGCCGGA GCATTCTTTCTTGAGGAGGGCAGCAAACGGGAAGAG |
| rro_FISH_handle_03 | TTGCTCCTTTCAAATATCGCGTTTGAGGGGGT GCATTCTTTCTTGAGGAGGGCAGCAAACGGGAAGAG |
| rro_FISH_handle_04 | GTAATAAGTTAGGCAGAGGCATTTATGATATT GCATTCTTTCTTGAGGAGGGCAGCAAACGGGAAGAG |
| rro_FISH_handle_05 | TTATTACGAAGAACTGGCATGATTGCGAGAGG GCATTCTTTCTTGAGGAGGGCAGCAAACGGGAAGAG |
| rro_FISH_handle_06 | TTCTACTACGCGAGCTGAAAAGGTTACCGCGC GCATTCTTTCTTGAGGAGGGCAGCAAACGGGAAGAG |

### Table S2: mini DNA origami staples

| name | sequence |
|---|---|
| mini_core_01 | ACTCTCGGGTTAAAGAGCACCATCCGGCGGC |
| mini_core_02 | ATCACTCCGCGAACAGTTTCACTGGTGCATAG |
| mini_core_03 | AGCCCGACTAGCTAATAAGCTCCTGAAACAAGTGGCGCAGTGCAGTA |
| mini_core_04 | CAGACGGTATTTGCCGTCAAATGGAGTCTGT |
| mini_core_05 | GATCGCTATATGTTCTATACCCACGTTAAGTT |
| mini_core_06 | CCTGACACCCGAGCATGTTACATTGGGAGCA |
| mini_core_07 | TGTCCGTATGGAGATATAGAACCCTTTCAGAG |
| mini_core_08 | AGGACCCGCCACGCCCTCGCTGCCATTATAC |
| mini_core_09 | GCTTTGAGCTCTCCTGTGTTGTGCGGGTTAGT |
| mini_core_10 | CGCCGGTCTCAGAAGGCCCAAACAGTGTATATCGAATCGCGGAAGTCT |
| mini_core_11 | GCTCCGTGAAGCAGCCGTGCTCCATCTTCGAT |
| mini_core_12 | TCGGGAGGAAGGACACTGTTATCCGTCCGGC |
| mini_core_13 | ATATTCATGGATCCAACCAATTTATTGGAGCT |
| mini_core_14 | CGTTTGACGAAGCTTGATTTAAGGCTTACCC |
| mini_core_15 | TGTGCCGGGAGTATTCCGATGAAAGGTATGT |
| mini_core_16 | GTTGATGCCTCTAGGTACGGATGGTTCAAAG |
| mini_core_17 | CTGCTCGCACGATCGATGGCTGATTAGTGCGG |
| mini_core_18 | TGGAGTTCGTCCGCATGGAGGGCCGTTCTTA |
| mini_core_19 | CGCCATATAGAGAACTGGTTCATTTTGCCAGC |
| mini_core_20 | ATTGGCCATTTACGGGACGCCGCACCGTACT |
| mini_FasL_handle_01 | GATCTACCAGTCATCGTCGTGCAATAACACGG TT TTCATTCTCCTATTACTACC |
| mini_FasL_handle_02 | CTTAAGCCATTGTTCAGGGAGTACAGGCTTG TT TTCATTCTCCTATTACTACC |
| mini_FasL_handle_03 | ACCAAGAACTCCGCTTGCAGAGGCAAAGGTT TT TTCATTCTCCTATTACTACC |
| mini_FasL_handle_04 | TACTTCAGTATCAGTAGTCCCTAAGGCTATGT TT TTCATTCTCCTATTACTACC |
| mini_FasL_handle_05 | GAATTTGACACGGCAGACATCGCGACTGACGC TT TTCATTCTCCTATTACTACC |
| mini_FasL_handle_06 | AATGACGTACGAGGGAATCCACTCCCACATGC TT TTCATTCTCCTATTACTACC |
| mini_biotin_handle_01 | GATCTACCAGTCATCGTCGTGCAATAACACGG TT [biotin] |
| mini_biotin_handle_02 | CTTAAGCCATTGTTCAGGGAGTACAGGCTTG TT [biotin] |
| mini_biotin_handle_03 | ACCAAGAACTCCGCTTGCAGAGGCAAAGGTT TT [biotin] |
| mini_biotin_handle_04 | TACTTCAGTATCAGTAGTCCCTAAGGCTATGT TT [biotin] |
| mini_biotin_handle_05 | GAATTTGACACGGCAGACATCGCGACTGACGC TT [biotin] |
| mini_biotin_handle_06 | AATGACGTACGAGGGAATCCACTCCCACATGC TT [biotin] |
| mini_FISH_handle_01 | GATCTACCAGTCATCGTCGTGCAATAACACGGGCATTCTTTCTTGAGGAGGGCAGCAAACGGGAAGAG |
| mini_FISH_handle_02 | CTTAAGCCATTGTTCAGGGAGTACAGGCTTG GCATTCTTTCTTGAGGAGGGCAGCAAACGGGAAGAG |
| mini_FISH_handle_03 | ACCAAGAACTCCGCTTGCAGAGGCAAAGGTT GCATTCTTTCTTGAGGAGGGCAGCAAACGGGAAGAG |
| mini_FISH_handle_04 | TACTTCAGTATCAGTAGTCCCTAAGGCTATGT GCATTCTTTCTTGAGGAGGGCAGCAAACGGGAAGAG |
| mini_FISH_handle_05 | GAATTTGACACGGCAGACATCGCGACTGACGCGCATTCTTTCTTGAGGAGGGCAGCAAACGGGAAGAG |
| mini_FISH_handle_06 | AATGACGTACGAGGGAATCCACTCCCACATGCGCATTCTTTCTTGAGGAGGGCAGCAAACGGGAAGAG |

### Table S3: wf DNA origami staples

| name | sequence |
|---|---|
| WF_core_001 | CGCCGCCAGCATTGACACCCCCCGTTCAGCCC |
| WF_core_002 | GGTTTGGCTCTTAGGGGAACCACCACCAGAGC |
| WF_core_003 | CCCTCAGAGCCGCCACCACACCGGAACCAGA |
| WF_core_004 | TCAGACGATTGGCCTTGCCACCCTCAGAGCCACCA |
| WF_core_005 | CGCCACCCTCAGAACCGATATTCACAAAC |
| WF_core_006 | TGGCTCCGCCTCCCTCAGAGC |
| WF_core_007 | GCCACCACCGGAAATCGGCATTTTCGG |
| WF_core_008 | TTTTCATAATCAAAATACTGTGAAGACGC |
| WF_core_009 | TTGGAAGGTCAGAATTAGCGTTTGCCATC |
| WF_core_010 | TCATAGCCCCCTTCCGTAATCAGTAG |
| WF_core_011 | ACTGTAGCGCGTTTTCTTTGATGATACAG |
| WF_core_012 | GTGCCTTGAGTAACAGTGTTTGCCTTTAGCGTCAG |
| WF_core_013 | CGACAGAATCAAGAGCACCATTACCA |
| WF_core_014 | ACCATCGATAGCAGCAAGCTGTCAACTGGGTT |
| WF_core_015 | AAGTAGGAGTTAAAGCAAACGTCACCAATGAA |
| WF_core_016 | TTAGCAAGGCCGGAATTATCACCGTC |
| WF_core_017 | CAGCAAAATCACCAGTCCCGTATAAACAGTTA |
| WF_core_018 | GTATCACCGTACTCAGATTTGGGAATTAGAGC |
| WF_core_019 | ACCGACTTGAGCCCATTCAACCGATT |
| WF_core_020 | TTATTCATTAAAGGTGTCCTTAGTTACTT |
| WF_core_021 | ACTCCTTATTACGTAAATATTGACGGAAA |
| WF_core_022 | GAGGGAGGGAAGGTTTGTCACAATCA |
| WF_core_023 | AAAGACAAAAGGGCGAGAGGTTTAGTACC |
| WF_core_024 | CCCTCAGAGCCACCACCCTCAATGGTTTACCAGCGCC |
| WF_core_025 | ATAGAAAATTCATTTTTCAGGGATAGCAAGCC |
| WF_core_026 | GGAATAAGTTTATCAGTATGTTAGCA |
| WF_core_027 | CAATAGGAACCCATGTACGCAAAGACACCAC |
| WF_core_028 | AACATATAAAAGAAACCGTAACACTGAGTTTCGTC |
| WF_core_029 | CCAAGTCGTATGGCTACATACATAAAGGTGGC |
| WF_core_030 | AACGTAGAAAATATCACCCTTCTTTCCGT |
| WF_core_031 | TAAGTAGACGCTACGGTGGCATGATTAAG |
| WF_core_032 | GATGCTACCGGTTCGTGTGCGGTGGCGTATGA |
| WF_core_033 | GTCAATGAAGTCTCCTCCTCTCCATGAAA |
| WF_core_034 | CAGTAGCACCAGATGGAGCTGGAGTAGTC |
| WF_core_035 | TCAAGATTGTTGATGGCGGTACCACATAC |
| WF_core_036 | TCTGTGGCAGCTTAGTTTCCCGCAGA |
| WF_core_037 | GTCTTACAAATGTAAAGTCCTGAACATTTACCTTC |
| WF_core_038 | GCACACCCCCTGCTAACCATACTAACTTT |
| WF_core_039 | CTAAGCCAGTATTATGCGATTGGTGA |
| WF_core_040 | ACCAGTACAAACTACAACTGTAACGATAGGCAAA |

| | |
|---|---|
| WF_core_041 | ACGGGTCGTACGCGCCTGTAGCATTCCACAGA |
| WF_core_042 | ATCTGCATCGCAACAATGCACTTTAT |
| WF_core_043 | CAGCCCTCATAGTTAGCGCGTCCGAGTTGTT |
| WF_core_044 | TACTCCGTGGTTACTTTAACGATCTAAAGTTT |
| WF_core_045 | TTTTCTGTATGGGCTCGTAAGAGATAGGA |
| WF_core_046 | AATGAGAATAACATTTAAAAAGGCCGTAA |
| WF_core_047 | TATCCAGCTGAACGGTGTTTAGTATACCC |
| WF_core_048 | ATGGTAATCCCTTATTACTTTTCTCCATTTTAGC |
| WF_core_049 | TCCAACAACCATTCATCTGAGGGCCC |
| WF_core_050 | TGCCTTTTGACTCATCTCACGCTGCGCGT |
| WF_core_051 | CGCCGCTACAGGGATATGATTTCTGCTTG |
| WF_core_052 | GGCGGCCATAGTCGGCCTGTTAAGTG |
| WF_core_053 | CGCCTACTGCGCTCGCTCAGCGATCC |
| WF_core_054 | GACCGCTGCGCCTTATGCAGCTCCCCACTAGA |
| WF_core_055 | GAGGAACTCTGGTAGGCGGTGCTACAGAG |
| WF_core_056 | TTCTTGAAGTGGTGGCGCCGCGCTTAATG |
| WF_core_057 | AACCACCACACCCCTTGATCCGGCAAACA |
| WF_core_058 | AACCACCGCTGGTAGCCGCTAGGGCGCTG |
| WF_core_059 | CGAAAGGAGCGGGCGCTCAGTGGAACGAA |
| WF_core_060 | CGTGGCGAGAAAGTGCCGTAAAGCAC |
| WF_core_061 | AACTCACGTTAAGGGAGAAAGCCGGCGAA |
| WF_core_062 | TTTAGAGCTTGACGGGATGAGTAAACTTGGTC |
| WF_core_063 | GCGGCATCAGCACCTTTAAAGGGAGCCCCCGA |
| WF_core_064 | TAAATCGGAACCCTGGAACCTCTTAC |
| WF_core_065 | GTGCCCGATCAAGAATCTCGATAACT |
| WF_core_066 | TCATTATGGTGAAAGTGTCGCCTTGCGTATAATAT |
| WF_core_067 | TGTCCATATTGGCCGGTAGTGATCTTATT |
| WF_core_068 | CAAAAAATACGCCATCAACGGTGGTA |
| WF_core_069 | TATCCAGTGATTTCTGGTTATAGGTACATTGA |
| WF_core_070 | GATGCCATTGGGATATCACGTTTAAATCAAAA |
| WF_core_071 | AACAAGGGTGAACACTTCAAAATGTTCTTTAC |
| WF_core_072 | GCAACTGACTGAAATGCCACTTGTGCTTATT |
| WF_core_073 | TTTCTTTACGGTCGTTTTGCTAAACAACT |
| WF_core_074 | AAGGCCGGATAAAATCCCATATCACCAGCTCA |
| WF_core_075 | TTCAACAGTTTCAGCGAAGAATGTGAATA |
| WF_core_076 | CATTCATCAGGCGGGCAAGGCTCCAAAAGGAGC |
| WF_core_077 | CTTTAATTGTATCGGTCGAAATTCCGGATGAG |
| WF_core_078 | CCGTCTTTCATTGCCATAAGAGCGATGAAAA |
| WF_core_079 | CTGGTGAAACTCACCCTCATGGAAAACGGTGT |
| WF_core_080 | CGTTTCAGTTTGCGCGAATAT |
| WF_core_081 | TCGTGGTATTCACTCCTTATCAGCTTGCTTTCG |
| WF_core_082 | AGGTGAATTTCGTGAGAAACTGCCGGAAATCG |
| WF_core_083 | ATGTGTAGCAAAAGGCCAGCA |

| | |
|---|---|
| WF_core_084 | CACGCCACATCTTAGGGATTGGCTGA |
| WF_core_085 | AAAGGCCAGGAACGTTTTCACCGTAA |
| WF_core_086 | TAGGGAAATAGGCCAGCGTAAAAAGGCCGCGTTGC |
| WF_core_087 | TTAAGCATTCTGCCGATTCTCAATAAACCCTT |
| WF_core_088 | GACGAAAAACATAGGGGCGAAGAAGT |
| WF_core_089 | TTGCCCATGGTGAAAACGCATGGAAGCCATC |
| WF_core_090 | TGACAGCTCGAGGCTTTGAACCTGAATCGCCA |
| WF_core_091 | ACAAACGGCATGACGATATCAAATTACGCCCC |
| WF_core_092 | TGGCGTTTTTCCATAGGCGTACTGTTGTAATTCA |
| WF_core_093 | GCCCTGCCACTCATCGCATCCGCCCCCCTGACGAGCA |
| WF_core_094 | ACAGGAGTCCAAGCGAGCTGGATTCTCACCAA |
| WF_core_095 | TCACAAAAATCGACGCTCGTCATTACTGGATCTATCA |
| WF_core_096 | AAATCCAGATGGAGTTCTGAGAAGTCAGAGGTGGCGAAACCC |
| WF_core_097 | GACAGGACTATAAAGATACCAGCGGCAACCGAGCGTTCTGAAC |
| WF_core_098 | TAAAAAACGCCCGAAGTTTTAAATCA |
| WF_core_099 | ATCTAAAGTATATTTTTGGTCATGAG |
| WF_core_100 | TTTTAAATTAAAAATGGGCGTTTCCCCCTGGA |
| WF_core_101 | AGCTCCCTCGTGCGCTATCTTCACCTAGATCC |
| WF_core_102 | ATTATCAAAAAGGTCCTTTGATCTTTT |
| WF_core_103 | CTACGGGGTCTGAGGTGGTTTTTTTGT |
| WF_core_104 | AAGGATCTCAAGAAGACTCCTGTTCCGACCCT |
| WF_core_105 | GCCGCTTACCGGATACATTACGCGCAGAAAAA |
| WF_core_106 | TTGCAAGCAGCAGTACCTTCGGAAAA |
| WF_core_107 | AGAGTTGGTAGCTCTAACTACGGCTA |
| WF_core_108 | CTCTGCTGAAGCCAGTCTGTCCGCCTTTCTCC |
| WF_core_109 | CTTCGGGAAGCGTGGCGTATTTGGTATCTGCG |
| WF_core_110 | CACTAGAAGAACAAACAGGATTAGCA |
| WF_core_111 | GAGCGAGGTATGTCCGGTAACTATCG |
| WF_core_112 | CCACTGGCAGCAGCCACTGGTGCTTTCTCATAGCTCACGCTGT |
| WF_core_113 | AGGTATCTCAGTTCGGTGTAGCCGGTAAGACACGACTTATCG |
| WF_core_114 | TCTTGAGTCCAACGTCGTTCGCTCCAAGC |
| WF_core_115 | TGGGCTGTGTGCACGAAGGAGGTTGAGGCAGG |
| WF_core_116 | GAAAATCTCCAAAAAAAAGAGTGAGAATAGAAAGGAA |
| WF_core_117 | TGTCGTCTTTCCAGACATAATAATTTTTTCACGTT |
| WF_core_118 | CAACTAAAGGAATTGCGAGTTAGTAAATGAA |
| WF_core_119 | CGGGGTTTTGCTCAGTACCCGCCACCCTCAGAACCGCCA |
| WF_core_120 | GCCACCCTCAGAACAGGCGGATAAGTGCCGTC |
| WF_core_121 | ATGCCCCCTGCCTATTATAGCCCGGAATAGGT |
| WF_core_122 | GAGAGGGTTGATATAAGTAGAGAAGGATTAGGATTAG |
| WF_core_123 | CTGAGACTCCTCATCGGAACCTATTA |
| WF_core_124 | GGAAAGCGCAGTCAAGTATTAAGAGG |
| WF_core_125 | TTCTGAAACATGAATAAGTTTTAACGGGGTCA |
| WF_core_126 | GAGTGTACTGGTATCTGAATTTACCG |

| | |
|---|---|
| WF_core_127 | GAATGGATCCTCACATACA |
| WF_core_128 | TTCCAGTAAGCGTTTAAAGCCAGAAT |
| WF_core_129 | ATACACAGAGTTATCGGATAGAACTTCT |
| WF_core_130 | ACTCGCGATAACCGTGTAGTAATTTATTT |
| WF_core_131 | CGTCACCTGGAGACGACGGGGGATTCA |
| WF_core_132 | ACGAAAACTTAAAGCAGACGAAGGGAAGAAAG |
| WF_core_133 | TTCCCCGAAAAGGTCGAGGACGACTACGGTCT |
| WF_FasL_handle_01 | TATTTTAATTCTAGGCGCCACGGCA TTTTCATTCTCCTATTACTACC |
| WF_FasL_handle_02 | AACCCAAAAGAACTCTCAGATACGTG TTTTCATTCTCCTATTACTACC |
| WF_FasL_handle_03 | AGTTTACAAGGAGCCCAGCATTGGCTACGCTAAG TTTTCATTCTCCTATTACTACC |
| WF_FasL_handle_04 | TTCCTTAGCTCCTGAAGCTATCCTAACGCA TTTTCATTCTCCTATTACTACC |
| WF_FasL_handle_05 | GCGCAGTTTTTCCGTTCCGCGCACA TTTTTCATTCTCCTATTACTACC |
| WF_FasL_handle_06 | GCAAGTGTAGCGGCATCCAGCAACGG TTTTCATTCTCCTATTACTACC |
| WF_biotin_handle_01 | TATTTTAATTCTAGGCGCCACGGCA TT [biotin] |
| WF_biotin_handle_02 | AACCCAAAAGAACTCTCAGATACGTG TT [biotin] |
| WF_biotin_handle_03 | AGTTTACAAGGAGCCCAGCATTGGCTACGCTAAG TT [biotin] |
| WF_biotin_handle_04 | TTCCTTAGCTCCTGAAGCTATCCTAACGCA TT [biotin] |
| WF_biotin_handle_05 | GCGCAGTTTTTCCGTTCCGCGCACA TT [biotin] |
| WF_biotin_handle_06 | GCAAGTGTAGCGGCATCCAGCAACGG TT [biotin] |
| WF_FISH_handle_01 | TATTTTAATTCTAGGCGCCACGGCA GCATTCTTTCTTGAGGAGGGCAGCAAACGGGAAGAG |
| WF_FISH_handle_02 | AACCCAAAAGAACTCTCAGATACGTG GCATTCTTTCTTGAGGAGGGCAGCAAACGGGAAGAG |
| WF_FISH_handle_03 | AGTTTACAAGGAGCCCAGCATTGGCTACGCTAAGGCATTCTTTCTTGAGGAGGGCAGCAAACGGGAAGAG |
| WF_FISH_handle_04 | TTCCTTAGCTCCTGAAGCTATCCTAACGCA GCATTCTTTCTTGAGGAGGGCAGCAAACGGGAAGAG |
| WF_FISH_handle_05 | GCGCAGTTTTTCCGTTCCGCGCACA GCATTCTTTCTTGAGGAGGGCAGCAAACGGGAAGAG |
| WF_FISH_handle_06 | GCAAGTGTAGCGGCATCCAGCAACGG GCATTCTTTCTTGAGGAGGGCAGCAAACGGGAAGAG |
